# Optimal information loading into working memory in prefrontal cortex explains dynamic coding

**DOI:** 10.1101/2021.11.16.468360

**Authors:** Jake P. Stroud, Kei Watanabe, Takafumi Suzuki, Mark G. Stokes, Máté Lengyel

## Abstract

Working memory involves the short-term maintenance of information and is critical in many tasks. The neural circuit dynamics underlying working memory remain poorly understood, with different aspects of prefrontal cortical (PFC) responses explained by different putative mechanisms. By mathematical analysis, numerical simulations, and using recordings from monkey PFC, we investigate a critical but hitherto ignored aspect of working memory dynamics: information loading. We find that, contrary to common assumptions, optimal loading of information into working memory involves inputs that are largely orthogonal, rather than similar, to the persistent activities observed during memory maintenance, naturally leading to the widely observed phenomenon of dynamic coding in PFC. Using a novel, theoretically principled metric, we show that PFC exhibits the hallmarks of optimal information loading. We also find that optimal loading emerges as a general dynamical strategy in task-optimized recurrent neural networks. Our theory unifies previous, seemingly conflicting theories of memory maintenance based on attractor or purely sequential dynamics, and reveals a normative principle underlying dynamic coding.

## Introduction

Working memory requires the ability to temporarily hold information in mind, and it is essential to performing cognitively demanding tasks^1,2^. A widely observed neural correlate of the maintenance of information in working memory is selective persistent activity. For example, in the paradigmatic memory-guided saccade task^3–13^, subjects must maintain the location of one out of several cues during a delay period after which they must respond with a saccade to the correct location (Fig. 1a). Cells in the lateral prefrontal cortex (lPFC) show elevated levels of activity that persist during the delay period and that is selective to the location of the now-absent cue^3–5,9^. However, neurons typically only reach a steady, persistent level of activity late in the delay period of a trial^6,8,10,11,14–20^. In contrast, during the cue and early delay period, neurons in lPFC often exhibit strong transient dynamics during a variety of working memory tasks^3,8,10,11,14–24^.

**Fig. 1.**
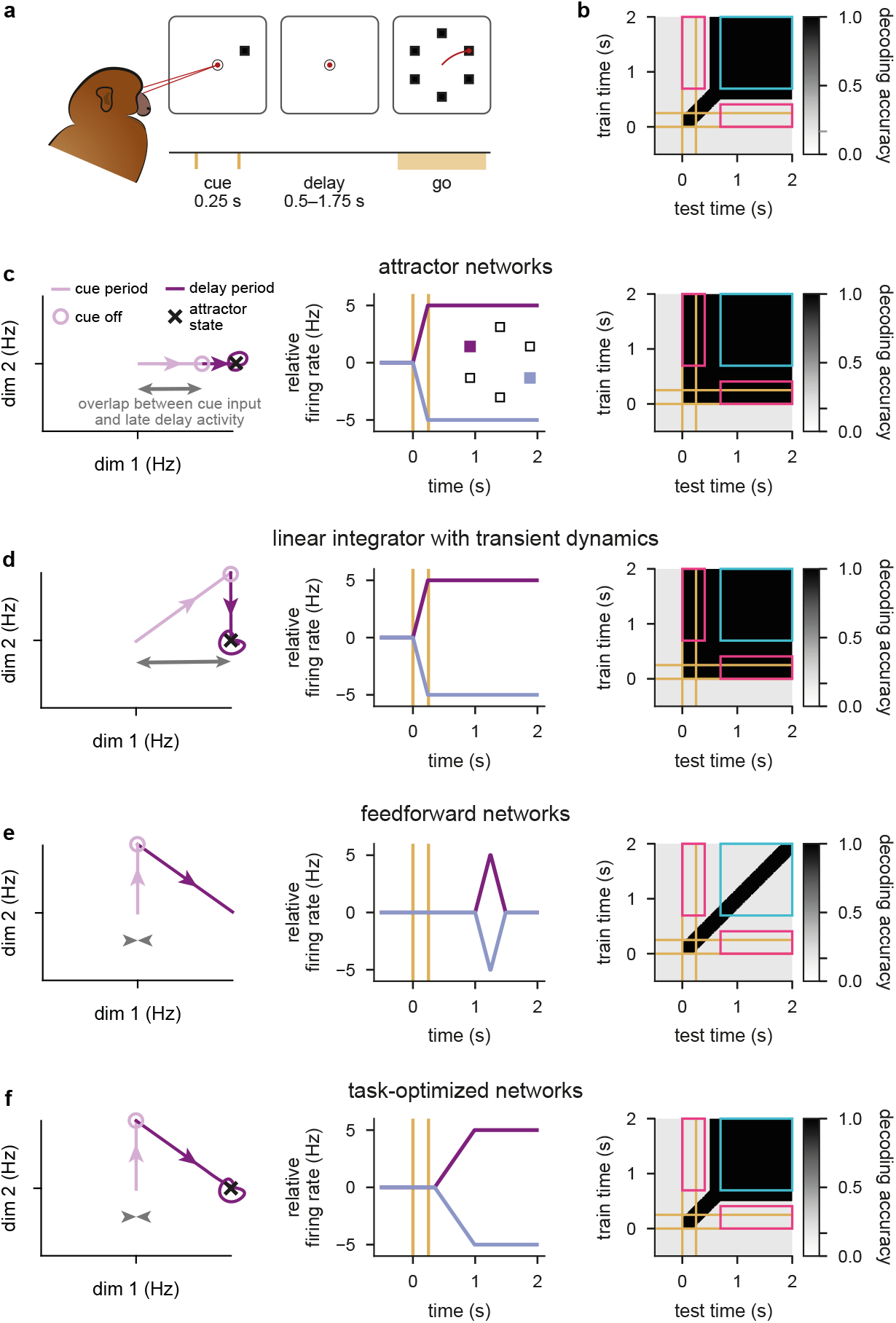
Neural dynamics in data and models during working memory. **a**, Illustration of the memory-guided saccade task. Time line of task events in a trial (bottom), with the corresponding displays (top). Top: black circle and squares show fixation ring, and the arrangement of visually cued saccade target locations, respectively (not to scale), red dots and line illustrate gaze positions during fixations and saccade, respectively. Bottom: yellow ticks show timing of stimulus cue onset and offset, yellow bar shows interval within which the go cue can occur. **b**, Schematic pattern of cross-temporal decoding when applied to neural recordings from the lPFC during working memory tasks^8,10,14–16,25^. Gray scale map shows accuracy of decoding cue identity (one out of 6) when the decoder is trained on neural activities recorded at a particular time in the trial (y-axis) and tested at another time (x-axis). Yellow lines indicate cue onset and offset times. Note poor generalization between time points inside the pink rectangle (i.e. dynamic coding), but good generalization between time points inside the cyan square (i.e. stable coding). The gray tick on the color bar indicates chance-level decoding. **c**, Schematic of neural network dynamics in an attractor network performing the task shown in **a** (see also Extended Data Fig. 1a,b). Left: trajectory in a low-dimensional projection of neural state space in a single cue condition during the cue period (pale purple line, ending in pale purple circle) and delay period (dark purple line). Purple arrow heads indicate direction of travel along the trajectory, black cross shows attractor state, gray arrow shows overlap between cue input and late delay activity. Center: time course of firing rates (relative to across-condition mean) of a neuron aligned with dim 1 from left panel for two cue conditions (purple vs. blue, see also inset). Yellow lines indicate cue onset and offset times. Right: cross-temporal decoding of neural activity in the network (cf. **b**; see also Extended Data Fig. 1a,b). **d–f**, Same as **c**, but for a linear integrator network with added transient dynamics^6,26^ (**d**; see also Extended Data Fig. 1c), a feedforward network that generates sequential activities^21,27^ (**e**; see also Extended Data Fig. 1d), and for a network optimized to perform the task shown in **a** (**f**; see also Extended Data Fig. 1e).

It remains unknown what mechanism underlies the combination of persistent and dynamically changing neural activities in lPFC—especially in light of recent population-level analyses. These analyses, using the technique of ‘cross-temporal decoding’, place particularly stringent constraints on any candidate neural mechanism of working memory maintenance. Cross-temporal decoding measures how well information about the cue location can be decoded from neural responses when a decoder is trained and tested on any pair of time points during a trial^8,10,11,14,15,25^ (Fig. 1b). These analyses reveal a consistent but somewhat puzzling set of results. First, when decoder training and testing times are identical, decodability is high (Fig. 1b, dark along the diagonal), confirming that information about cue location is indeed present in the population at all times. Decodability is also high when both training and testing occurs during the late delay period, suggesting that even if there are changes in neural responses during this period, the coding of cue location remains stable (Fig. 1b, black inside cyan square). However, decoding performance remains low when a decoder is trained during the cue or early delay period and tested during the late delay period, and vice-versa (Fig. 1b, light gray inside pink rectangles). This demonstrates that the neural code for cue location undergoes substantial change between these these two periods—a phenomenon that has been called ‘dynamic coding’^8,10,14–16,25^.

Classically, the neural mechanism of working memory maintenance is thought to rely on attractor network dynamics. Attractor networks^5,7,12,28–33^, and closely related ‘integrator’ networks^34,35^, naturally account for selective persistent activity (Fig. 1c, left and middle). However, in these models, neurons show limited transient activity during the delay period, and crosstemporal decoding reveals stable coding throughout the whole trial, lacking the characteristic dynamic coding seen in experimental data (compare Fig. 1b to c, right). This behavior emerges across several variants of attractor networks, whether they express a continuum of persistent activity patterns (‘ring’ or ‘bump’ attractor networks) or a finite number of discrete patterns (Extended Data Fig. 1a–b; see also Supplementary Information S1). Critically, even when external inputs were specifically chosen so that neural activity showed longer transient dynamics^6,26^ (Fig. 1d), these inputs still relied on a large overlap with the desired persistent state (Fig. 1d, left). As a result, these models also exhibited strongly stable stimulus coding over time (Fig. 1d, right and Extended Data Fig. 1c) and the transient dynamics were regarded as being purely epiphenomenal^6,26^.

To capture transient dynamics more naturally, a very different class of models have been developed based on mechanisms that generate neural activity sequences. These models typically rely either on effectively feedforward network connectivity^21,27^ or chaotic network dynamics^24,36–38^. The dynamics of such models rapidly transition between orthogonal subspaces over time (Fig. 1e, left), thus cross-temporal decoding is high only between neighbouring time-points (Fig. 1e, black along diagonal). Although such models are ideally suited to capturing transient neural responses (Fig. 1e, center), they fail to exhibit persistent activities and stable coding during the late delay period (Fig. 1e, right; gray inside blue square). Therefore, previous work leaves open two interrelated key questions: how can a neural circuit exhibit early sequential dynamics followed by stable late-delay dynamics, and more importantly, why would it use such a counterintuitive dynamical regime?

In order to study the network mechanisms underlying the combination of persistent and dynamic neural activities during working memory, we build on recent advances in using task-optimized neural networks^13,17,20,24,36,39–41^. We find that the behaviour of such task-optimized networks unifies attractor and sequential activity models, showing both early transient dynamics and late persistent activities, giving rise to dynamic coding (Fig. 1f). To understand the principles and functional significance of this dynamical behavior, we focus on a hitherto ignored aspect of the operation of attractor networks: optimal information loading. Through numerical simulations and mathematical analyses, we show that inputs that most efficiently drive network activities into a desired attractor state tend to be orthogonal to the attractor state itself (Fig. 1f, left). Critically, this results in an initial period of strong transient dynamics with dynamic coding (Fig. 1f, right), which are thus fundamental and functionally useful features of attractor dynamics when used with optimal inputs. Based on our theoretical results, we develop a specific neural measure for assessing whether a network uses optimal information loading. Using this measure, we demonstrate key signatures of optimal information loading in neural recordings from lPFC. Finally, we show that optimal information loading emerges naturally in task-optimized neural networks with a variety of architectures, including linear integrators, as well as nonlinear discrete and ring attractor models.

Our results offer a novel, normative perspective on a core but hitherto ignored component of attractor networks dynamics—information loading— and challenge long-held assumptions about pattern completion-like mechanisms in neural circuits.

## Results

### Pattern completion and optimal information loading in attractor networks

Traditional approaches to studying attractor networks used models in which the connectivity between neurons was constrained to be effectively symmetric^5,7,28,30,32,34,35,42–45^, making the analysis of their dynamics mathematically more convenient^28,34,42,46^. Thus, we first replicated results with such symmetric networks that were optimized to perform the working memory task shown in Fig. 1a. In particular, we defined optimal information loading to be achieved by a set of inputs when they maximize the performance of a network in terms of how well the cue can be decoded from its neural activities at the end of the delay period. For simplicity, we only modelled the intrinsic dynamics of the network during the delay period and the effect of the cue was captured by cue-specific initial neural activities (i.e. neural activities at the beginning of the delay period^35,42,43^; Fig. 2b). To study optimal information loading, we optimized these initial activities for cue-decodability at the end of the delay period (Methods 1.3.1).

**Fig. 2.**
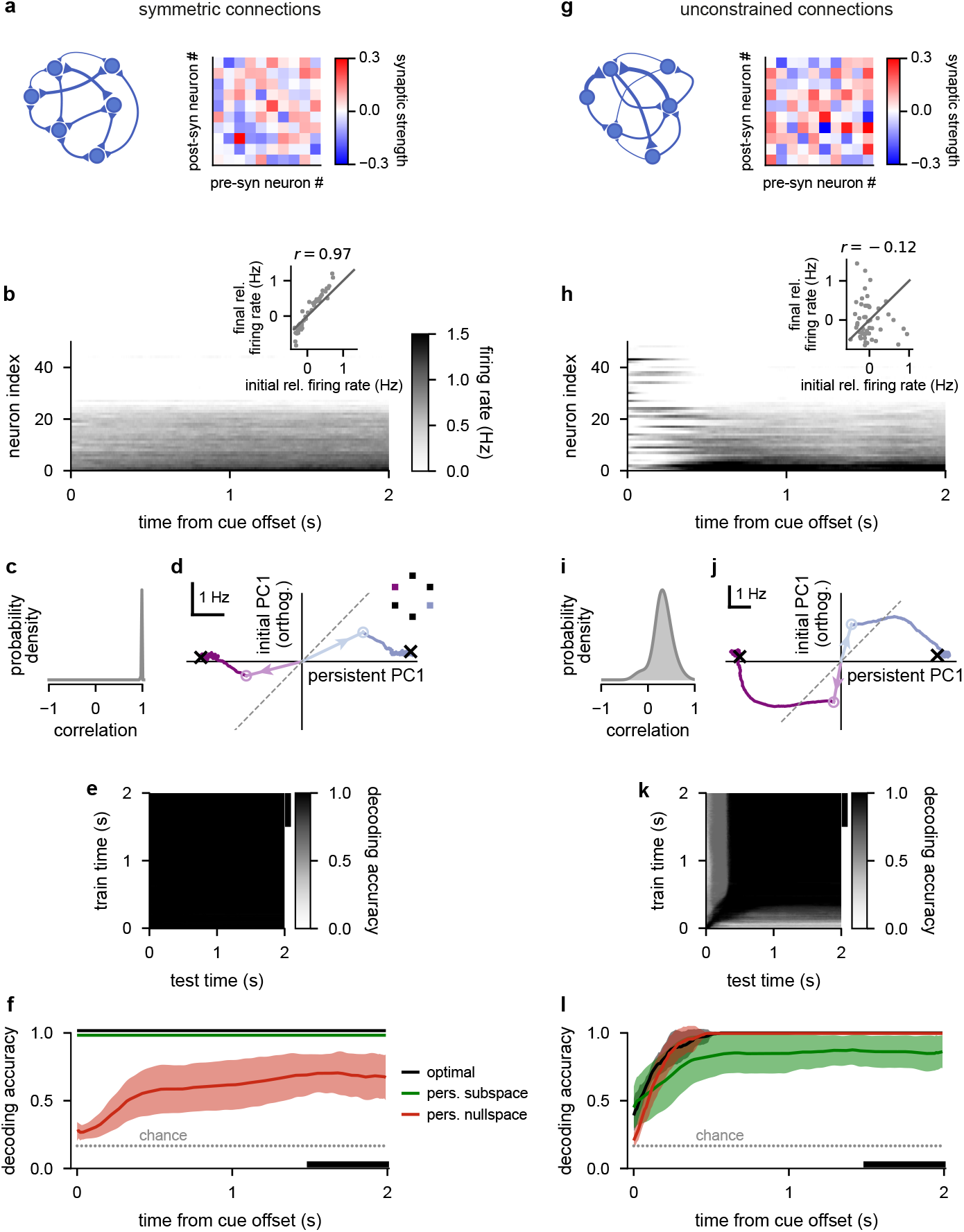
Pattern completion and optimal information loading in attractor networks. **a**, A network with symmetric connections. Left: network schematic. Right: the recurrent weight matrix for 10 of the 50 neurons. **b**–**f**, Analysis of neural responses in symmetric attractor networks (such as shown in **a**) with optimized initial conditions. **b**, Firing rates in a representative trial. Neurons are ordered according to their rates at the end of the trial. Inset shows initial vs. final firing rates (mean-centered, i.e. relative to the time-dependent but condition-independent mean) across neurons in this trial (gray dots) and their Pearson correlation (*r*; *p <* 0.001). Gray line is the identity line. **c**, Distribution of Pearson correlations between initial and final mean-centered neural firing rates across all 6 cue conditions and 10 networks. **d**, Sub-threshold activity for 2 cue conditions in an example network. Horizontal axis (persistent PC1) shows network activity projected on to the 1st principal component (PC1) of activities at the end of the delay period (across the 2 conditions shown in the inset), vertical axis (initial PC1, orthogonalized) shows projection to PC1 of initial activities orthogonalized to persistent PC1. Pale open circles (with arrows pointing to them from the origin) show the optimized initial conditions, dark traces show activity trajectories, black crosses show stable fixed points, dashed gray line is the identity line. **e**, Cross-temporal decoding of neural firing rate activity (cf. Fig. 1b). The black vertical bar on the right indicates the delay-trained decoder training time period from **f. f**, Performance of a delay-trained decoder (black bar indicates decoding training time period) on neural firing rate activity over time starting from optimized initial conditions with full optimization (black), or restricted to the 5-dimensional subspace spanning the 6 cue-specific attractors (persistent subspace, green), or the subspace orthogonal to that (persistent nullspace, red). Solid lines and shading indicate mean 1 s.d. across all 6 cue conditions and 10 networks. Gray dotted line shows chance level decoding. Green and black lines are slightly offset vertically to aid visualization. **g–l**, Same as **a**–**f**, for attractor networks with unconstrained connections. The Pearson correlation in **h** (inset) is not significant (*p >* 0.4).

Optimal initial activities gave rise to classical pattern completion dynamics in symmetric networks. First, initial activities were noisy versions of (and in fact highly similar to) the desired persistent patterns (Fig. 2b inset, and Fig. 2c). Second, the ensuing dynamics were driven directly into the corresponding persistent state, resulting in only small and gradual changes in activities over the delay period (Fig. 2b). Further analysis of these dynamics showed that the optimal initial activities aligned well with directions in neural state space that best distinguished between the desired persistent activities (Fig. 2d, ‘persistent PC1’ component of pale arrows and circles; Extended Data Fig. 2b), with only a comparably small component in orthogonal directions specific to these initial activities (Fig. 2d, ‘initial PC1, orthogonalized’) which subsequently changed little over time (Fig. 2d, dark trajectories). As a result, cross-temporal decoding performance was high for all pairs of times (Fig. 2e), and—as a special case—a decoder based on templates of neural activity during the late delay period (i.e. during the steady state of the network), generalized well to all times and was able to decode the cue identity from neural activities with high accuracy throughout the delay period (Fig. 2f, black line).

The similarity between initial and persistent activities was critical for these networks. When constrained to use initial activities that were orthogonal in neural state space to persistent activities (i.e. lying in the ‘persistent nullspace’), these networks performed substantially more poorly (Fig. 2f, red line) and activity often did not settle into the correct attractor state (Extended Data Fig. 2d). In contrast, explicitly enforcing these networks to use initial activities that were similar to persistent activities (i.e. lying in the ‘persistent subspace’) did not compromise their performance (Fig. 2f, green line; Extended Data Fig. 2c). Thus, when connectivities were constrained to be symmetric, our approach using explicitly optimized inputs and connectivities recapitulated earlier results obtained with classical attractor networks using hand-crafted inputs and connectivities^5,7,12,28,30,33,42^.

In contrast, attractor networks optimized without a symmetry constraint exhibited dynamics distinctly unlike simple pattern completion (Fig. 2g–l). First, initial activities resembled persistent activity much less than in symmetric networks (Fig. 2i), such that their correlation could even be negative (Fig. 2h inset). Second, neural activities often underwent substantial and non-monotonic changes before ultimately settling into an attractor state (Fig. 2h). This was also reflected in optimal initial activities (Fig. 2j, pale arrows and open circles) being strongly orthogonal to persistent activities (Fig. 2j, black crosses; Extended Data Fig. 2f), with this orthogonality decaying over the delay period (Fig. 2j, dark trajectories). Such dynamics are consistent with PFC recordings from primates performing a variety of working memory tasks^8,17,22–24,32,47–49^. Decoding analyses revealed further similarities with experimental data: a decoder trained on neural activity from the late delay period generalized poorly to early times (Fig. 2k, and Fig. 2l, black line) and vice versa (Fig. 2k), thus exhibiting a fundamental signature of ‘dynamic coding’^8,10,14–16^ (cf. Fig. 1b). Importantly, we found that the orthogonality of initial conditions in these networks was instrumental for high performance: in a double dissociation from symmetrically constrained networks, restricting initial conditions to be in the persistent subspace (Fig. 2l, green line; Extended Data Fig. 2g), but not in the persistent nullspace (Fig. 2l, red line; Extended Data Fig. 2h), diminished decodability at the end of the delay period (cf. Fig. 2f).

The above results were obtained with networks storing a small number of discrete attractors, corresponding to the six cue conditions. Previous work found that several aspects of working memory dynamics in lPFC are better captured by networks in which instead a large number (or even a continuum) of attractor states form a ring in neural state space^5,7,44,45^. Thus, we repeated our analyses on optimized networks while explicitly encouraging such a ring attractor to form during optimization (Methods 1.3.4). We found a highly similar pattern of results in ring attractor networks as compared with discrete attractor networks (Extended Data Fig. 3).

### Dynamical analysis of optimal information loading

To understand why optimal information loading in classical symmetrically constrained versus unconstrained attractor networks is so different, and in particular why inputs orthogonal to attractor states are optimal for unconstrained networks, we reduced these networks to a canonical minimal model class consisting of only two neurons^35,50,51^. For analytical tractability, we considered networks with linear dynamics (i.e. in which neurons had linear activation functions). Critically, with the appropriate set of synaptic connections, even linear networks can exhibit persistent activity^6,26,34,35,46,52^—the key feature of working memory maintenance in attractor networks.

For our analyses, we again distinguished between models with symmetric connectivity between neurons (Fig. 3a; top)^34,35,51^, and models without this constraint (Fig. 3a; bottom)^6,26^. In either case, the specific connection strengths were chosen to create illustrative examples providing intuitions that—as we show below— also generalize to large networks with randomly sampled connection strengths (Fig. 3d–e, Fig. 4). The dynamics of these networks are fully described in a two-dimensional neural state space spanned by the activities of the two neurons (Fig. 3b) and define a flow-field in this space determining how neural activities change over time (Fig. 3b; blue arrows). An important subspace of the full neural state space of these networks is the ‘persistent subspace’ corresponding to persistent patterns of activities. In our two-neuron linear networks, the persistent subspace simply corresponds to a line onto which the neural activities ultimately converge over time (Fig. 3b; green lines showing the persistent mode). Therefore, the persistent mode allows these networks to distinguish between two stimuli depending on which side of the origin the state of the network is. The larger the magnitude of its activity along this persistent mode at the end of the delay period, the more robustly the identity of the stimulus can be decoded (e.g. in the presence of noise, as we show below).

**Fig. 3.**
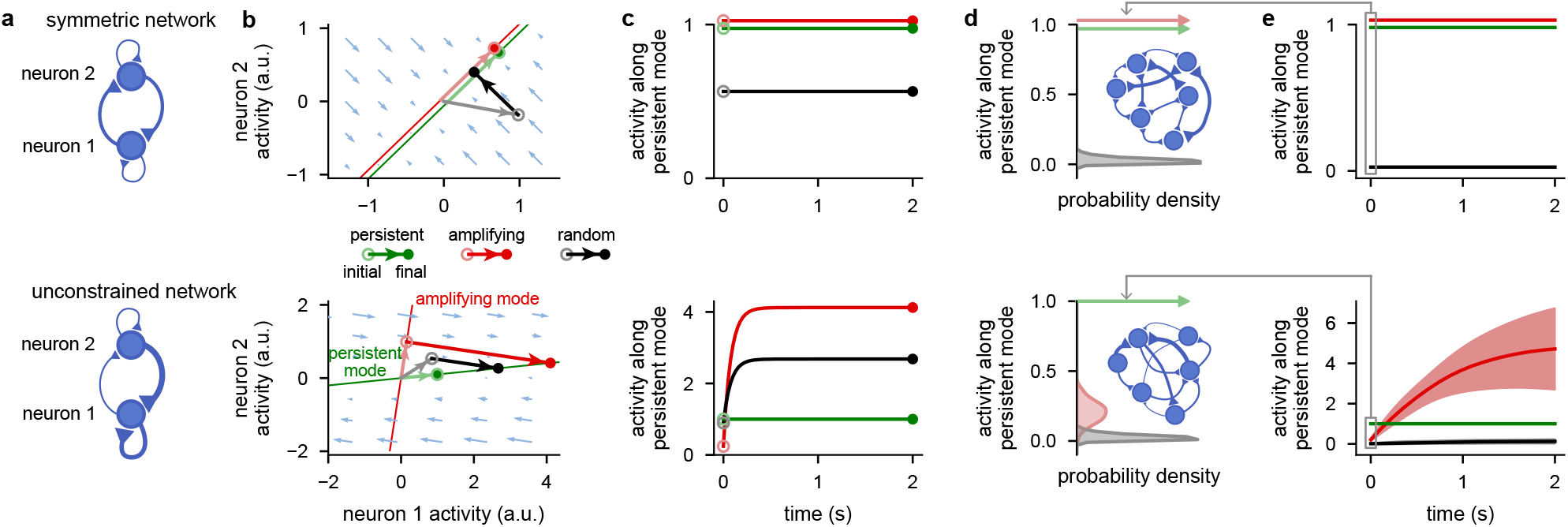
Dynamical analysis of optimal information loading. **a**, Architecture of a symmetric (top) and an unconstrained network (bottom). **b**, Neural state space of the symmetric (top) and unconstrained network (bottom). Pale blue arrows show flow field dynamics (direction and magnitude of movement in the state space as a function of the momentary state). Thin green and red lines indicate the persistent and most amplifying modes, respectively (lines are offset slightly in the top panel to aid visualisation). Pale green, red, and gray arrows with open circles at the end indicate persistent, most amplifying, and random initial conditions, respectively. Dark green, red, and black arrows show neural dynamics starting from the corresponding initial condition. (Green arrows, and the red arrow in the top panel cannot be seen, as no movement in state space happens from those initial conditions.) Filled colored circles indicate final (persistent) neural activity. **c**, Time course of network activity along the persistent mode (i.e. projection onto the green line in **b**) when started from the persistent (green), most amplifying (red), or random initial conditions (black) for the symmetric (top) and the unconstrained model (bottom). **d**, Distributions of absolute overlap with the persistent mode for persistent (pale green), most amplifying (pale red), or random initial conditions (gray) across 100 randomly connected 1000-neuron symmetric (top) or unconstrained networks (bottom). The persistent (and for the symmetric models, also the equivalent most amplifying) initial conditions produce delta functions at 1 (arrows). Insets show illustration of large networks of neurons with either symmetric (top) or unconstrained (bottom) connections. **e**, Time course of absolute overlap with the persistent mode when starting network dynamics from persistent (green), most amplifying (red), or random initial conditions (black) for the symmetric (top) and the unconstrained network (bottom). Lines and shaded areas show mean±1 s.d. over the 100 randomly sampled 1000-neuron networks from **d**.

**Fig. 4.**
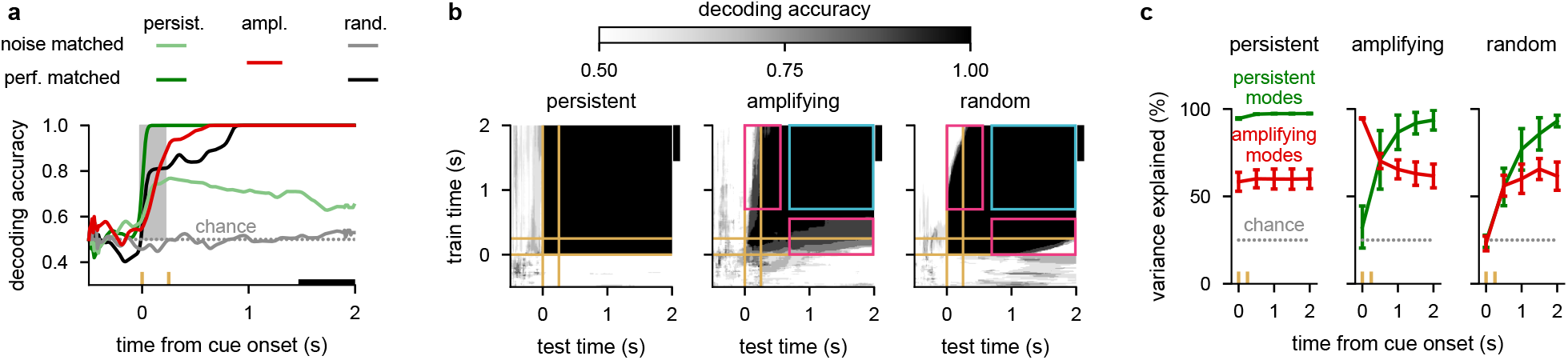
Neural signatures of optimal information loading. **a**, Performance of a delay-trained decoder (black bar indicates decoder training time period) on neural activity over time. Two cue conditions were used with inputs that were identical but had opposite signs. Lines show mean across 10 randomly connected 100-neuron linear unconstrained networks. Yellow ticks on horizontal axis indicate cue onset and offset times and the gray shading indicates the cue period. We show results for inputs aligned with the persistent mode (dark and pale green), the most amplifying mode (red), or a random direction (black and gray). Light colors (pale green and gray, ‘noise-matched’) correspond to networks with the same level of noise as in the reference network (red), while dark colors (dark green and black, ‘performance-matched’) correspond to networks with the same level of asymptotic decoding performance as that in the reference network (red). Gray dotted line shows chance level decoding. **b**, Cross-temporal decoding of neural activity for the 3 different information loading strategies (persistent, most amplifying, and random respectively in left, center, and right panels) for a representative network for the performance-matched condition from **a**. Yellow lines indicate cue onset and offset times. Pink rectangles indicate poor generalization between time points (i.e. dynamic coding) and cyan squares indicate examples of good generalization between time points (i.e. stable coding). The black vertical bars on the right of each plot indicate the delay-trained decoder training time period from **a. c**, Percent variance of responses explained by the subspace spanned by either the 25% most persistent (green) or 25% most amplifying (red) modes as a function of time in the same networks analyzed in **a**. Lines and error bars show mean ±1 s.d. across networks. We show results for inputs aligned with the persistent mode (left), most amplifying mode (center), or a random direction (right). Gray dotted line shows chance level overlap with a randomly chosen subspace occupying 25% of the full space.

To understand the mechanisms of information loading, we considered three distinct stimulus input directions. We then analysed the time course of the neural activities projected onto the persistent mode^6,26,30^ after being initialised in each of these directions. First, we considered inputs aligned with the persistent mode, the input direction studied in classical attractor networks^6,26,34,35,51^ (Fig. 3b; pale green arrows and open circles). Second, we considered the ‘most amplifying mode’, which is defined as the stimulus direction that generates the most divergent and thus best discriminable activity over time^53–57^ (Methods 1.7.1; Fig. 3b, red lines, and pale red arrows and open circles). Third, we considered a random input direction (Fig. 3b; gray lines/circles).

We were able to show mathematically that optimal information loading, in the sense of maximizing overlap with the persistent mode at sufficiently long delays, is always achieved with inputs aligned with the most amplifying mode (Supplementary Information S2). Equivalently, the most amplifying mode is the input direction that requires the smallest magnitude initial condition to achieve a desired level of persistent activity (i.e. a desired level of performance). More generally, we could also show both mathematically and in simulations (Extended Data Fig. 4) that the most amplifying mode is near optimal in achieving a desired level of performance while minimizing total neural activity over time (i.e. the total energy used by the network) for sufficiently long delay lengths.

In symmetric networks, the most amplifying mode is aligned with the most persistent mode (Fig. 3b; top)^58,59^, and thus does not generate activity transients (Fig. 3c; top)—accounting for the simple pattern completion dynamics seen in classical attractor networks with symmetric connectivity^5,7,28,30,32,34,35,42,43^ (Fig. 2a–f). However, in unconstrained networks, the most amplifying mode is typically different from the most persistent mode (Fig. 3b; bottom). Intuitively, this is because effective feedforward connections exist in unconstrained networks^21,27,50,56,60^. For example, neurons 1 and 2 in the example network shown in Fig. 3a (bottom) respectively align strongly with the persistent and amplifying modes (Fig. 3b, bottom). Thus, feeding neuron 1 indirectly through the feedforward connection from neuron 2 can increase its activity more than just feeding it directly. This means that activity evolving from the most amplifying mode exhibits a distinct transient behaviour: its overlap with the most persistent mode is initially low and then increases over time (Fig. 3c; bottom, red line), accounting for the richer transients seen in unconstrained attractor networks (Fig. 2g–l). Thus, there is a form of ‘speed–accuracy’ trade-off between whether inputs should use the most amplifying or persistent mode: if information is required immediately following stimulus offset, such as in a perceptual decision-making task^13,40,59^, inputs need to use the persistent mode. However, if there is a time delay until the information is needed, as is the case in all working memory tasks^2,61^, then the most amplifying mode becomes the optimal input direction. Indeed, an analogous trade-off was already apparent between the persistent sub- vs. nullspace inputs in the nonlinear attractor networks we analysed earlier (Fig. 2l, red vs. green).

The insights obtained in the simple two-neuron network also generalized to large randomly connected linear integrator networks, with more than two neurons (Fig. 3d,e; see Methods 1.4.1). Moreover, as network size grows, in unconstrained (but not in symmetric) networks, the most amplifying direction becomes increasingly orthogonal to the most persistent mode^62^, further accentuating the advantage of amplifying over persistent mode inputs^62^ (Fig. 3d–e, Extended Data Fig. 5a–b; red vs. green). This is because in large unconstrained networks, there are many effectively feedforward motifs embedded in the full recurrent connectivity of the circuit, which can all contribute to transient amplification^21^. Random initial conditions become fully orthogonal in both networks and result in poor overlap with the persistent mode (Fig. 3d–e, Extended Data Fig. 5a–b; black). Numerical simulations confirmed that these results also generalized to networks with noisy dynamics (Extended Data Fig. 5c). Moreover, explicitly optimizing the initial condition of such a network so as to maximize the persistent activity it generated at the end of a delay period also made this initial condition overlap strongly with the network’s most amplifying mode (Extended Data Fig. 5d).

As our mathematical analyses only applied to linear dynamics, we used numerical simulations to study how they generalized to nonlinear dynamics. We found that the same principles applied to the dynamics of a canonical 2-dimensional nonlinear attractor system (analogous to the networks in Fig. 3a–c), when the persistent and most amplifying directions were defined locally around its ground state (Methods 1.6; Extended Data Fig. 6, see also Supplementary Information S3). Importantly, we also found that large optimized nonlinear neural networks (with discrete or ring attractors) also showed a similar pattern of results (Extended Data Fig. 3e, and Extended Data Fig. 7a–c, see also Supplementary Information S4).

### Neural signatures of optimal information loading

Our dynamical analysis suggested that there should be clearly identifiable neural signatures of a network performing optimal information loading. To demonstrate this, and to allow a more direct comparison with data, we used the same large, randomly connected, unconstrained networks that we analysed earlier (Fig. 3d–e, bottom), with noisy dynamics (as in Extended Data Fig. 5c–d) and the cue period modelled using temporally extended constant inputs— mimicking typical experiments^3–5,10^ (Fig. 4). We studied the three different information loading strategies that we identified earlier: inputs aligned with either the persistent mode, the most amplifying mode, or a cue-specific random direction.

We began by conducting a decoding analysis using templates of late delay activity, as is often done for prefrontal cortical recordings^6,8,10,14,15,25^ (and also in Fig. 2f,l). We first verified that for a fixed level of neuronal noise, the most amplifying inputs were indeed optimal for achieving high decodability at the end of the delay period (Fig. 4a, compare red line to pale green and gray lines). We were also able to show mathematically that, in line with our original definition of optimal information loading, the most amplifying inputs in noisy linear networks are optimal for maximizing average decodability during the delay period (Supplementary Information S2.7). In contrast, random inputs performed considerably more poorly (Fig. 4a, gray line). Remarkably, persistent mode inputs achieved a similarly low level of decodability at late delay times (Fig. 4a, compare pale green and gray lines).

The level of noise in the networks we have studied so far was not constrained by data, which typically shows high decodability^6,8,10,14,15,25^. This is important because the sub-optimal input conditions (Fig. 4a, pale green and gray lines) could achieve high decoding performance by appropriately reducing the noise level in our simulations (Fig. 4a, asymptotic values of dark green and black lines). Thus, asymptotic decoding performance alone cannot be used to identify the information loading strategy employed by a network. To address this, in subsequent analyses, we used networks in which the level of late-delay performance was matched between the three information loading strategies by appropriately reducing the level of noise when using persistent or random inputs. Nevertheless, a critical difference emerged between the different information loading strategies even in these ‘performance-matched’ networks. For both random and most amplifying input directions, the delay-trained decoder only performed well when tested late in the delay period (Fig. 4a, black and red lines), whereas for inputs aligned with the persistent direction this decoder performed near ceiling at all times after cue onset (Fig. 4a, dark green line).

Next, in order to more fully characterise the differences between persistent versus random or most amplifying inputs, and for a comprehensive comparison with experimental data^8,10,14,15,25^, we also employed full cross-temporal decoding (Fig. 4b). This analysis showed that all information loading strategies led to dynamics in which stimulus information was present at all times after cue onset (Fig. 4b, diagonals are all black). Moreover, for the persistent mode inputs, stimulus information was maintained using a ‘stable code’^10,11,14,16^ (Fig. 4b, left, all off-diagonals are black)—similar to previous integrator models of working memory^34,35^ (Extended Data Fig. 1c). In contrast, random and most amplifying mode inputs led to poor cross-temporal decodability between early and late time points after cue onset (Fig. 4b, center and right, off-diagonals indicated by pink rectangles are white/gray). This gave rise to the phenomenon of ‘dynamic coding’^8,10,11,14–16^, and suggested sequential activities during the early-to-late delay transition^21,27,36^. These activities then stabilised during the late delay period as the network dynamics converged to a persistent pattern of activity (Fig. 4b, center and right, off-diagonals inside cyan squares are black). In sum, these decoding analyses were able to clearly distinguish between persistent mode and random or amplifying inputs, but not between the latter two.

To clearly distinguish between networks using most amplifying inputs or merely a random input direction, we constructed a targeted measure for identifying networks using most amplifying inputs. To achieve this, we exploited the fact that in large networks, random inputs typically have negligible overlap with any other direction in neural state space, including the most amplifying mode. Thus, we directly measured the time courses of the overlap of neural activities with the top 25% most amplifying modes. We quantified this overlap as the fraction of across-condition variance of neural activities that these modes collectively explained (Fig. 4c, red lines; Methods 1.7.3). For a comparison, we also measured the overlap of neural activities with the top 25% most persistent modes (Fig. 4c, green lines).

Persistent mode inputs led to constant high and moderate overlaps with the persistent and most amplifying modes, respectively (Fig. 4c, left). Random inputs started with chance overlap for both modes, which then increased to the same levels that resulted from persistent mode inputs (Fig. 4c, right). In contrast, most amplifying inputs were uniquely characterised by a cross-over between the time courses of the two overlap measures. Initially, neural activities overlapped strongly with the most amplifying mode, but showed only chance overlap with the persistent mode (Fig. 4c, middle). Over time, these overlap measures changed in opposite directions, such that by the end of the delay period overlap was high with the persistent mode and lower with the most amplifying mode (Fig. 4c, middle). Therefore, the cross-over of these overlap measures can be used as a signature of optimal information loading utilizing inputs aligned with the most amplifying modes.

To further illustrate how our overlap measures can distinguish between optimal and random input directions, we modified an earlier integrator model of working memory^6^ (Extended Data Fig. 1c, Extended Data Fig. 8a,d) so that inputs lay in a purely randomly oriented subspace. This resulted in cross-temporal decoding matrices that looked similar to that achieved by the most amplifying mode (Extended Data Fig. 8b), but the overlap measures that we developed here clearly revealed the lack of optimal information loading, even in this modified model (Extended Data Fig. 8 e). In addition, we confirmed in numerical simulations that the same signature of optimal information loading remains detectable (and distinguishable from other information loading strategies) even under the practical constraints of experimental data analysis: when the underlying network dynamics is nonlinear, and only accessible indirectly by fitting linear dynamical models to the neural responses they generate (Extended Data Fig. 7d, Methods 1.4.3 and Supplementary Information S4.4).

### Signatures of optimal information loading in monkey lPFC

To study whether the PFC shows the dynamical signatures of optimal information loading that our theoretical analyses identified, w e a nalysed a d ata set^48^ of multi-channel recordings of the lateral prefrontal cortex (lPFC) in two monkeys during a variable-delay memory-guided saccade task (Fig. 1a). These recordings yielded 438 and 625 neurons (for monkeys K and T, respectively; Extended Data Fig. 9, Methods 1.1). We analysed the population dynamics of all recorded neurons in each monkey and applied the same metrics to this dataset that we applied to our models. Population dynamics appeared to show rich transient dynamics during the cue and early delay period, followed by relatively stable dynamics during the late delay period (Fig. 5a). This was reminiscent of the dynamics we found in unconstrained attractor networks following optimal information loading (Fig. 2h).

**Fig. 5.**
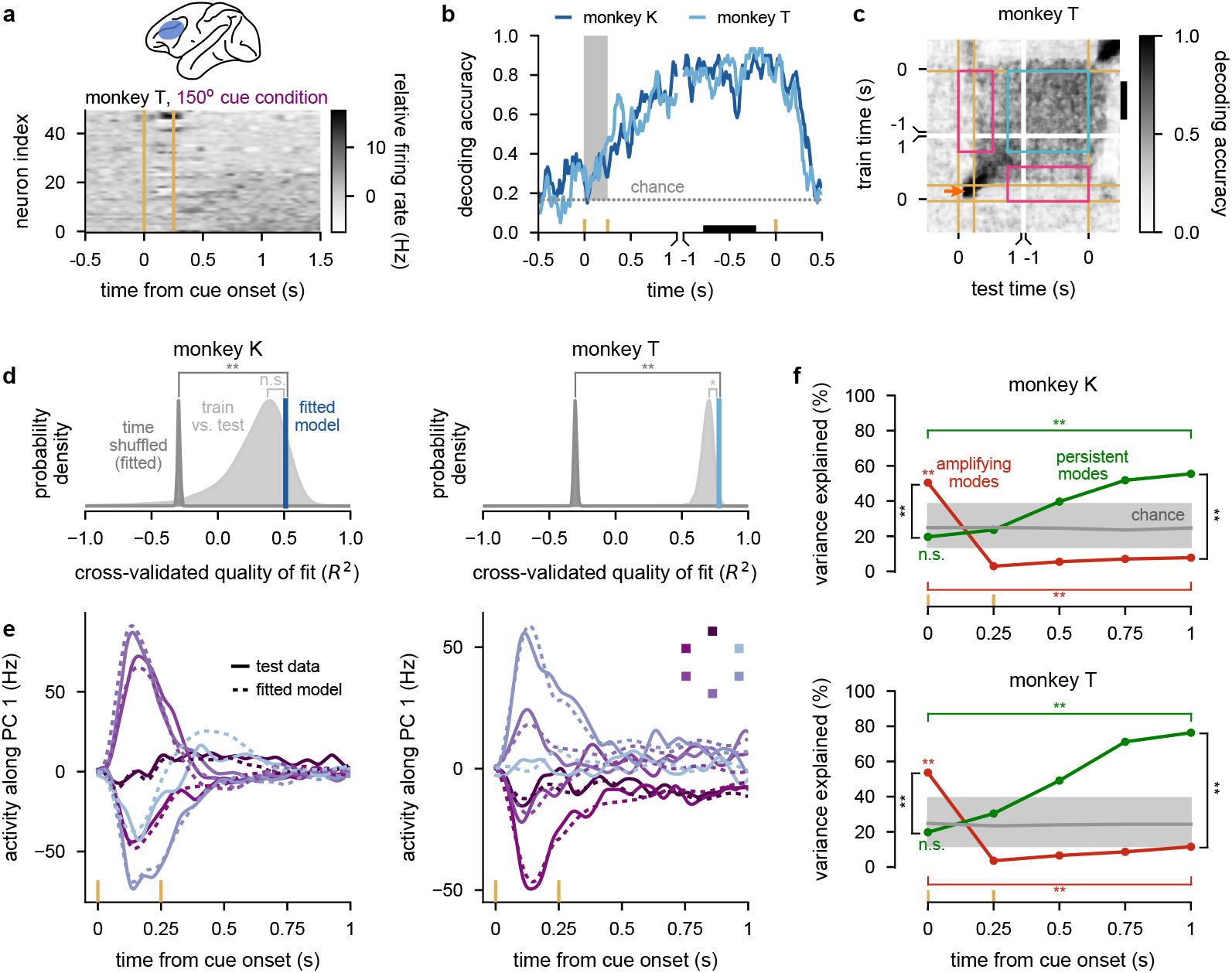
Signatures of optimal information loading in monkey lPFC. **a**, Top: lPFC recording location. Bottom: neural firing rates (relative to the time-dependent but condition-independent mean) for one stimulus cue condition for 50 example neurons. See Fig. 1a for experimental paradigm. Neurons are ordered according to their firing rate at the end of the period shown. Vertical yellow lines indicate stimulus cue onset and offset. **b**, Performance of a delay-trained decoder (black bar indicates decoder training time period) on neural activity over time. Yellow ticks on horizontal axis indicate stimulus cue onset, offset, and go cue times, and the gray shading indicates the stimulus cue period. Data is aligned to either stimulus cue onset (first 1.5 s) or to the go cue (final 1.5 s). Gray dotted lines show chance level decoding. **c**, Cross-temporal decoding of neural activity for monkey T (see Extended Data Fig. 10a for Monkey K). Yellow lines indicate stimulus cue onset, offset, and go cue times. Pink rectangles indicate poor generalization between time points (i.e. dynamic coding) and the cyan square indicates examples of good generalization between time points (i.e. stable coding). The orange arrow indicates good same-time decoding during the cue period. The black vertical bar on the right indicates the delay-trained decoder training time period from **b. d**, Cross-validated quality of fits when fitting 20-dimensional linear neural networks to neural activity (blue) and time shuffled controls (dark gray). We also show quality of fits of the data against itself (‘train vs. test’; light gray). **e**, Neural activity for each of the 6 cue conditions projected onto the top PC (solid lines) for monkey K (left) and monkey T (right). Solid lines show held-out test data, dashed lines show predictions of fitted model dynamics. The inset for monkey T shows which color corresponds to each cue condition. **f**, Percent variance of responses explained by the subspace spanned by either the 25% most persistent (green) or 25% most amplifying (red) modes as a function of time for the 20-dimensional linear neural networks fitted to data from monkey K (top) and monkey T (bottom). Gray lines show chance level overlap defined as the expected overlap with a randomly chosen subspace occupying 25% of the full space (median and 95% C.I. across 200 random subspaces). Comparisons shown in **d** and **f** use two-sided permutation tests (*, *p <* 0.05; **, *p <* 0.01; n.s., not significant).

To further quantify this behaviour, we conducted decoding analyses. First, we found that a delay-trained decoder did not generalize to times outside of the delay period (Fig. 5b). In particular, performance was near-chance level during the cue period and increased over the first 1 s of the delay period—in line with previous studies^6,10,14–16,25^. This was distinct from the pattern completion dynamics seen in classical attractor network models of working memory (Fig. 2f,l green and Fig. 4a green), but similar to that expected from random or optimal inputs in unconstrained networks (Fig. 2l black and red; Fig. 4a bottom, black and red).

Full cross-temporal decoding reinforced these results: decoders trained during the delay period did not generalize to the cue or go periods and vice versa (Fig. 5c and Extended Data Fig. 10a, pink rectangles). Thus, neural activity exhibited dynamic coding^14,15^ rather than the stable coding characteristic of simple pattern completion (Fig. 1c right; Fig. 4b left; and Extended Data Fig. 1a–c right). Importantly, same-time decoding performance was close to 1 throughout the cue and delay periods (Fig. 5c and Extended Data Fig. 10a, orange arrow). This confirmed that the poor cross-temporal generalization between early and late periods of a trial was not because the cue information had not yet reached PFC, or was maintained by activity-silent mechanisms^11,41,45^. At the same time, also in line with previous studies^8,10,14–16^, we found relatively stable coding during the late delay period (Fig. 5c and Extended Data Fig. 10a, cyan square). This ruled out purely sequential activity-based dynamics^21,27,37,38,63^ (Fig. 1d and Extended Data Fig. 1d).

Quantifying the relative alignment of the subspaces occupied by neural dynamics across time using PCA^6,64^ confirmed t he o rthogonality o f n eural activities between different task periods (Extended Data Fig. 10b–c). Further analyses showed that this orthogonality was not simply due to distinct sub-populations of neurons being active in different task periods (due to either feedforward connections between these populations, or single-neuron adaptation mechanisms), but was instead largely due to changes in population-wide activities patterns^10,49^ (Extended Data Fig. 10d–e).

These results, in line with previous findings^8,10,15,16^, clearly indicated that activities during the cue period were largely orthogonal from those during the delay period. However, these analyses alone were unable to distinguish between two fundamentally different information loading strategies PFC could employ: random input directions, or optimal input directions. Thus, in order to clearly identify the information loading strategy underlying the combination of dynamic and stable coding that we found, we applied our overlap measure (Fig. 4c) to these PFC recordings. For this, we first fitted a 20-dimensional linear dynamical system model to the cue and early delay periods of our recordings (0–1 s after cue onset, Methods 1.4.3). We confirmed that linear dynamics provided a reasonably accurate cross-validated fit to the data compared to a time shuffled control (which destroyed the lawful dynamics of the data; Fig. 5d, dark gray, see also Methods 1.4.3), and model-free train vs. test performance (which indicated that cross-validated errors were mostly due to sampling noise differences between the train and test data; Fig. 5d, light gray) and recapitulated the most important aspects of the trial-average dynamics in each condition (Fig. 5e).

We then performed the same overlap analysis on the fitted linear dynamics of the data that we used on our simulated networks with linear dynamics (Fig. 4c; Methods 1.7.3). As expected from our decoding analyses (Fig. 5b,c), the overlap of neural activities with the most persistent modes was at chance initially and gradually increased (Fig. 5f, green and Extended Data Fig. 10i). Critically however, the overlap of neural activities with the most amplifying modes was high initially and decreased with time (Fig. 5f, red and Extended Data Fig. 10i). Consistent with these results, we found that at early times, stimulus information was just as decodable within the amplifying subspace as in the full space and was more poorly decodable in the persistent subspace (Extended Data Fig. 10h, *t* = 0). Later in the delay period, stimulus information was significantly better decodable in the persistent subspace than in the amplifying subspace (Extended Data Fig. 10h, *t >* 0).

We also noted that the overlap with the most amplifying directions became significantly lower than chance over time. This suggests that PFC circuits may be more mathematically ‘non-normal’^21,27,56,57,60^ than the networks with randomly chosen weights that we used in Fig. 4. For example, Extended Data Fig. 8f shows this phenomenon in a highly non-normal (purely feedforward) network using optimal information loading (see also Discussion).

As a control, we repeated the same analyses on time-shuffled data, or on data taken from the late delay period (when the network should already be near an attractor state). Neither control analyses resulted in the same cross-over pattern that we found in our main analysis. In particular, the overlap with the most amplifying modes remained at (or below) chance at all times (Extended Data Fig. 10f,g,i).

Therefore, these analyses provide strong experimental evidence that PFC circuit dynamics utilize optimal information loading with inputs aligning with the most amplifying modes (compare to Fig. 4c; middle and Extended Data Fig. 10i, third vs. fourth row) rather than simply using random input directions (compare to Fig. 4c; right and Extended Data Fig. 10i, first vs. fourth row).

### Information loading in task-optimized nonlinear networks

The definition of most amplifying inputs relies on full access to the algebraic form of the dynamics of a network, something that the brain will not have explicitly when performing a working memory task. In turn, the formal equivalence of using the most amplifying input directions to optimal information loading could only be established for networks with linear dynamics receiving instantaneous inputs, while fixing the magnitude of those inputs. Thus, an important question is whether optimizing simple task-relevant cost functions in nonlinear networks^13,17,20,24,39–41,65^, under only a generic energy constraint^13,39–41,65^, can be sufficient for such networks to adopt an optimal information loading strategy.

We trained nonlinear recurrent networks (Fig. 6a; Methods 1.3.2) on the same memory-guided saccade task as that which our animals performed (Fig. 1a). Following previous approaches^13,39,40^, all recurrent weights in the network, as well as weights associated with the input and read-out channels, were optimized, while only penalizing the average magnitude of neural responses over the course of the whole trial (Methods 1.3.3).

**Fig. 6.**
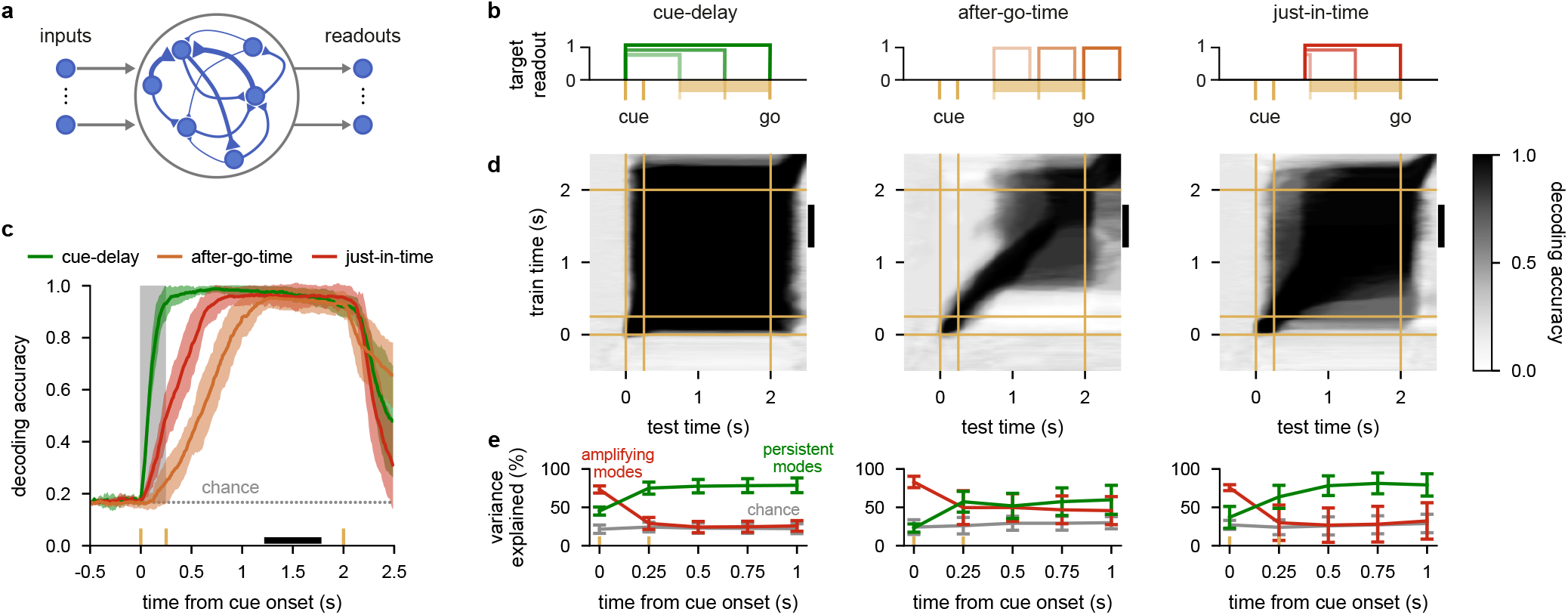
Information loading in task-optimized nonlinear networks. **a**, Illustration of a recurrent neural network model with unconstrained connectivity (middle). During the cue period, networks received input from one of six input channels on any given trial depending on the cue condition (left). Network activity was decoded into one of six possible behavioural responses via six readout channels (right). All recurrent weights in the network (50 neurons), as well as weights associated with the input and readout channels, were optimized. **b**, Illustration of cost functions used for training. Yellow ticks indicate cue onset and offset times, yellow bars indicate range of go times in the variable delay task. Boxcars show intervals over which stable decoding performance was required in three example trials with different delays for each of the cost functions considered: cue-delay (left), after-go-time (center), or just-in-time (right). **c**, Performance of a delay-trained decoder (black bar indicates decoder training time period on model neural activity over time in trials with a 1.75 s delay. Yellow ticks show stimulus cue onset, offset, and go times, and the gray shading indicates the cue period. Neural activities were generated by networks optimized for the cue-delay (green), after-go-time (orange), or just-in-time (red) costs. Solid colored lines and shading indicate mean ±1 s.d. across 10 networks. Gray dotted line shows chance level decoding. **d**, Cross-temporal decoding of model neural activity for cue-delay (left), after-go-time (center), and just-in-time (right) trained models. Yellow lines indicate stimulus cue onset, offset, and go times. The black vertical bars on the right of each plot indicate the delay-trained decoder training time period from **c. e**, Percent variance of responses explained by the subspace spanned by either the 25% most persistent (green) or 25% most amplifying (red) modes as a function of time for 20-dimensional linear neural networks fitted to the model neural activities of nonlinear networks optimized for the cue-delay (left), after-go-time (center), or just-in-time cost (right). Gray lines show chance level overlap defined as the expected overlap with a randomly chosen subspace occupying 25% of the full space. Lines and error bars show mean±1 s.d. over 10 networks.

To study the generality of optimal information loading, we first implemented two standard cost functions that have been widely used in previous work^13,17,24,39,40^. These cost functions required networks to maintain cue information either stably throughout the delay period, starting immediately after cue onset (cue-delay; Fig. 6b, left), or only at response time (after-go; Fig. 6b, center). Both networks achieved high performance, as measured by a late-delay decoder, in line with what their respective cost functions required: immediately after cue onset for the cue-delay cost (Fig. 6c and Extended Data Fig. 11a, green), or only shortly before go time for the after-go-time cost (Fig. 6c orange and Extended Data Fig. 12b).

We then further analyzed the dynamics with which these networks achieved competent performance. In particular, we evaluated whether they employed optimal information loading, and how well they reproduced critical aspects of the empirical data. The cue-delay network showed signatures of classical attractor dynamics with simple pattern completion: cross-temporal decoding was high at all times, including between the cue and delay periods (Fig. 6d, left; cf. Fig. 1c, Extended Data Fig. 1a–c), neural activity overlapped strongly between the cue and delay periods (Extended Data Fig. 11c, left), and at the time of cue offset, neural activity was already very close to its final attractor location in state space (Extended Data Fig. 11d, left). In line with our theory of optimal information loading, this was achieved by neural activities during the cue period aligning predominantly with the most amplifying modes (Fig. 6e, left, red). However, at the same time, activities were also already aligned well above chance with the most persistent modes (Fig. 6e, left, green). This was consistent with these networks being explicitly required to exhibit stable coding at all times by the cue-delay cost. These features also made this network a poor match to the experimental data, which showed a combination of dynamic and stable coding and at-chance overlap of activities with the most persistent mode during the cue period (Fig. 5b–c,f,Extended Data Fig. 10a–b). We also found similar behavior for networks optimizing a ‘full-delay’ cost, in which cue information must be stably maintained only after cue offset (Extended Data Fig. 13, Methods 1.3.3).

At the other extreme, the after-go-time network did not make particular use of attractor dynamics. Instead, it generated largely sequential activities, i.e. pure dynamic coding akin to the dynamics of a feedforward network: cross-temporal decoding was only high at the very end of the delay period (Fig. 6d, center; cf. Fig. 1d and Extended Data Fig. 1d, right), neural activity was strongly orthogonal between the cue and delay periods (Extended Data Fig. 12d, left), and these networks did not exhibit attractor states (Extended Data Fig. 12e, left). This was particularly the case for a fixed delay task, for which this cost function yielded purely sequential dynamics (Extended Data Fig. 12c–e, right). As required by optimal information loading, neural activities also had a strong initial overlap with the most amplifying modes in this network (Fig. 6e, center, green). However, as expected for sequential dynamics, the overlap with the most persistent modes never significantly exceeded that with the most amplifying modes (Fig. 6e, center). Again, the apparent lack of attractor dynamics was well explained by the cost function not requiring any stable coding during the delay period. Therefore, this network also deviated from the data in important ways, in this case by failing to exhibit stable coding and high overlap with the persistent mode during the late delay period (cf. Fig. 5b–c,f,Extended Data Fig. 10a–b). In summary, network dynamics trained for standard cost functions exhibited optimal information loading and recovered classical network models of working memory (Fig. 1c,d and Extended Data Fig. 1a–d), but were different from those seen in experimental recordings^8,10,14–16,25^ (Fig. 5b,c,f).

However, we reasoned that neither of these standard cost functions may be appropriate for understanding PFC function. The cue-delay cost is well justified when stimuli need to be decoded potentially instantaneously after cue onset, and as such it is most relevant for sensory areas^59^. Conversely, the after-go-time cost may be most directly relevant for motor areas, by only requiring stable coding during the short response period^65^. Therefore, we also considered a third cost function that required stable coding just in time before the go cue appeared, i.e. during a period that was divorced from the stimulus or response time windows, and as such was more consistent with the putative role of PFC in cognitive flexibility^2,25,61^ (just-in-time; Fig. 6b, right).

In contrast to both standard training costs, just-in-time networks showed the signatures of a combination of attractor and sequential dynamics which were consistent with its cost function. The performance of a late-delay decoder was high only after cue offset but remained so for most of the delay period (Fig. 6c and Extended Data Fig. 11a, red), cross-temporal decoding was poor between early and late periods of a trial, but high during the late delay period (Fig. 6d, right; Extended Data Fig. 11b; cf. center, Fig. 5c; see also Extended Data Fig. 11d for state-space plots), neural activity was strongly orthogonal between the cue and delay periods (Extended Data Fig. 12d, right), and at the time of cue offset, neural activity was far from its final attractor location in state space (Extended Data Fig. 11d, right). Critically, the overlap of neural activities with the most amplifying and persistent modes showed the characteristic cross-over that we found experimentally (Fig. 6e, right; cf. Fig. 5f). Thus, this network both used optimal information loading and reproduced the key features of the experimental data.

In summary, all task-optimized networks exhibited a key feature of optimal information loading: they made use of most amplifying modes early during the trial (Fig. 6e, all red lines start high at 0 s). The extent to which they showed the complete cross-over of amplifying and persistent overlaps predicted by our earlier analyses (Fig. 4c, center), and characteristic of the experimental data (Fig. 5f), was consistent with how much they were required to exhibit stable coding^8,10,11,14–16^. These results suggest that optimal information loading emerges naturally as a dynamical strategy in task-optimized networks, without explicit requirements on their inputs.

## Discussion

While attractor networks have been proposed to underlie a number of core cognitive functions^12,17,17,28–31,33–35,42,51,66–68^, prominently including working memory^5–7,26,29,31–33,69^, their operation was almost exclusively analyzed in terms of how their intrinsic connectivity supports information maintenance^5,7,12,29–31,34,35,70,71^ (but see Refs. 6,26, discussed below). We instead studied information loading by external inputs in attractor networks and showed that optimal information loading provides a normative account of the widely observed and puzzling phenomenon of dynamic coding^8,10,14–16^. We predict that these results should also generalize to more cognitively demanding working memory tasks in which, unlike in the simple memory-guided saccade task we studied here, the correct response is unknown during the delay period, thus requiring the maintenance of stimulus information before a response can be prepared^14,15,23,72,73^. Indeed, strongly dynamic population activity, similar to those that we identified here, has been observed in monkey PFC^10,14–16,23,24,73^ and in neural networks^20,24,39^ trained on such tasks. To fully test the generality of these principles, it will also be important to extend the theory and these analyses even further, to tasks with multiple delay periods^8,15^.

For understanding the dynamics of optimal information loading, we used networks whose connectivity was constrained to be symmetric as a pedagogical stepping stone. Some classical attractor and integrator networks indeed used purely symmetric connectivities^28,33–35,42^, and these continue to form the basis of our analytical understanding of the capacity, noise-tolerance, and input amplification o f m ore realistic networks^46^. The perfect symmetry of classical models has been relaxed by more recent, highly influential models of working memory that instead used quasi-symmetric connectivities, i.e. connectivities that were only weakly non-symmetric. These include models whose connectivity is not strictly symmetric at the microscopic level of cell-to-cell connections, but their macroscopic connectivity (at the level of connections between groups of similarly tuned cells) is strongly symmetric^71,74^, as well as models in which excitatory cells are connected symmetrically, but perfect symmetry is broken by the introduction of effectively a single inhibitory neuron providing a spatially uniform, global level of inhibition^5,7,12,30,32,44,45^. Indeed, previous analyses of such weakly non-symmetric networks^5–7,30,44^ and our simulations of such networks revealed largely stable coding dynamics (e.g. see Extended Data Fig. 1a–b). In contrast, we showed that dynamic coding naturally arises in networks whose connectivity is not constrained to be symmetric, and especially so under optimal information loading.

Our dynamical analysis revealed a novel, theoretically-grounded aspect of dynamic coding: not only should neural activities during the cue and early delay period be orthogonal to those during the late delay period, but they should be orthogonal in the specific directions that are aligned with the most amplifying directions. We found strong evidence for these predictions of optimal information loading in lPFC during a memory-guided saccade task. These results unify previous, seemingly conflicting m odels o f working memory maintenance that typically either use attractor dynamics^5,7,29^ or rely on sequential activities often generated by non-normal dynamics^21,27,36,37^.

We found that although both classes of models can capture select aspects of neural data (e.g. sequential models can capture early delay activity whereas attractors are better suited to capturing late delay activity), no model could capture the experimentally observed rich combination of sequential and persistent dynamics^72^ (Fig. 1; see also^39^). We showed that optimal information loading in attractor models with realistic, unconstrained connectivity, leads to the specific combination of sequential and persistent dynamics that has been observed in experiments. We found that this was true across a range of different specific network architectures: using either hand-set (Figs. 3 and 4 and Extended Data Fig. 5a,b) or optimized stimulus inputs (Extended Data Fig. 5c,d); and linear integrator (Figs. 3 and 4 and Extended Data Fig. 5), nonlinear discrete attractor (Figs. 2 and 6 and Extended Data Figs. 2, 7 and 11–13) or nonlinear ring attractor dynamics (Extended Data Fig. 3).

In contrast to our optimal information loading-based account, previous attempts to reconcile transient and persistent dynamics specifically p roposed t hat transient dynamics do not affect the delay (or ‘mnemonic’) coding of the stimulus information^6,26^. These stable delay dynamics are very different from dynamic coding as observed in experiments^3,8,10,11,14–24^, and as predicted by our theory of optimal information loading. Put simply, in previous models, the stimulus input is strongly aligned with the desired persistent state (Fig. 1d, left). In real data, and in models that exhibit optimal information loading, stimulus inputs drive network activity strongly orthogonal to the desired persistent state (and specifically in a direction that is aligned with the most amplifying mode) before activity ultimately settles into the correct state (Fig. 1f, left). Indeed, previously observed high correlations between cue and delay periods^6^, which partially motivates using inputs aligned with the persistent state, are likely due to high overall baseline firing rates, and they have been shown to disappear (and even become negative) when data is mean-centered across cue conditions^8,24^.

There are aspects of the data that were not reproduced accurately by any of the specific models we implemented. First, the overlap with the most amplifying directions became significantly lower than chance over time in the data. This suggests that PFC circuits may be more mathematically ‘non-normal’ (i.e. include stronger effective feedforward loops^21,27,57^, or excitatory–inhibitory interactions^53,56^) than the networks with randomly chosen or initialised weights we used here^60,62^. (For example, we found that networks with strong feedforward connectivity reproduced this phenomenon; Extended Data Fig. 8f.) Second, the time evolution of the overlaps with the most persistent and most amplifying modes seemed to obey different time constants, with the persistent overlap evolving substantially slower than the amplifying overlap. This may be a result of high dimensional, graded dynamical transitions between multiple amplifying and persistent modes compared to the less complex dynamical transitions that we observed in our models. More generally, analysing the data at single trial resolution, as opposed to the across-trial averages we analysed, may provide further important constraints on the underlying circuit dynamics^72,75,76^. Conversely, constraining the models to be more biologically plausible, e.g. by using spiking neurons or Hebbian forms of plasticity, may provide better fits to the data and more detailed predictions.

There have been multiple mechanisms proposed to account for some of the features of the data, most prominently dynamic coding^14,15^, that previously seemed to be at odds with basic attractor network dynamics These hypothetical mechanisms include in short-term plasticity^11,23,39,41,77^, specific changes the strength of input and recurrent connections^44,78^, and separate stimulus- and delay-responsive cells^3,10^. We showed that the core phenomenon of dynamic coding emerges naturally, without any of these additional mechanisms, from the same ultimate principle that explains persistent activities (robust memory maintenance implemented by attractor dynamics). Moreover, the high initial overlap with the most amplifying modes, which was a core prediction of our theory confirmed by the data and our optimized networks, is not specifically predicted by any of these alternative mechanisms. Nevertheless, these mechanisms are not mutually exclusive with ours. In fact, they might help explain the more nuanced aspects of the data that our specific network implementations did not capture (see above), as well as aspects of the data that lie outside the scope of our theory (e.g. activity silent information maintenance during inter-trial intervals^45^).

A number of recent studies of neural network dynamics have analysed the relationship between the direction of inputs and the magnitude of responses they evoke^53,57,59,62^. However, these studies focused on networks with transient dynamics, such as those relevant for perception^59^, or motor control^53,62^. In particular, Ref. 62 found that optimal inputs (resulting in the largest transients) are typically orthogonal to the activity patterns that the network expresses in response to them, providing a normative account for the experimentally observed orthogonality of preparation and execution subspaces in motor cortex^64,79^. Our work suggests that the use of optimal inputs to drive network dynamics, and the orthogonality of those inputs to network responses, is a more general principle of cortical circuits, extending beyond the motor cortex. In particular, our results demonstrate the importance of optimal initialization even when the transients following initialization themselves may be irrelevant, as information is ultimately maintained by stable attractor states.

In line with our results, previous studies optimizing networks on related tasks requiring persistent, rather than transient, responses also exhibited key features of dynamic coding: neural activities initially pointed strongly orthogonal to the ultimate attractor location in state space^17^; the dynamics during the stimulus period had 0 correlation with late delay activity^24^; and cross-temporal decoding of time revealed strongly sequential dynamics in a variety of tasks^20^ (see also^39,41^ for related results). In fact, these features of model activities were also shown to be reflected in the corresponding experimental data in each case^17,20,24^. Nevertheless, it remained unclear whether these features were epiphenomenal or an integral part of the functioning of these networks. Our results suggest optimal information loading as a unifying principle underlying these observations.

## Acknowledgements

This work was funded by the Wellcome Trust (Investigator Award in Science 212262/Z/18/Z to M.L. and Sir Henry Wellcome Postdoctoral Fellowship 215909/Z/19/Z to J.S.), the Human Frontiers Science Programme (Research Grant RGP0044/2018 to M.L.), the Biotechnology and Biological Sciences Research Council (award BB/M010732/1 to M.G.S.), the James S. McDonnell Foundation (award 220020405 to M.G.S.), the Japan Society for the Promotion of Science (JP18K03197 and JP21K03141 to K.W., 18H05380 to T.S.), and the Japan Science and Technology Agency (CREST JPMJCR186 to T.S.). For the purpose of open access, the authors have applied a CC-BY public copyright license to any author accepted manuscript version arising from this submission. We thank Flavia Mancini, John Duncan, Guillaume Hennequin, Yashar Ahmadian, and Kris Jensen for useful feedback and detailed comments on the manuscript.

## Author Contributions

J.P.S., M.L., and M.G.S. conceived the study. K.W. performed all experimental recordings, T.S. assembled the neural recording system, and K.W. and T.S. performed data pre-processing. J.P.S. and M.L. developed the theoretical framework, performed analytical derivations, and wrote the first draft of the manuscript. J.P.S. performed all numerical simulations, analysed the data, and produced the figures. J.P.S., M.L., and M.G.S. interpreted the results. All authors revised the final manuscript.

## Competing Interests statement

The authors declare no competing interests.

## Reporting Summary

Further information on research design is available in the Life Sciences Reporting Summary linked to this article.

## Randomization

No new experimental data was gathered for this paper. There is no group allocation in this study. Trial types were randomly determined by a computer program.

## Blinding

As data collection had been performed well before our theory of optimal information loading was developed and our corresponding analyses were performed, it was effectively blind to the purposes of our study. Data analysis was not performed blind to the conditions of the experiments.

## Data exclusion

As described below (Methods 1.1.4), 1 neuron from monkey K’s dataset was removed from all analyses because it was recorded in fewer than 10 trials for at least one stimulus cue condition.

## Data availability

All experimental data will be made available in the following repository upon peerreviewed publication: https://github.com/jakepstroud.

## Code availability

All code was custom written in Python using NumPy, SciPy, Matplotlib, Scikit-learn, and Tensorflow libraries. All code will be made available in the following repository upon peer-reviewed publication: https://github.com/jakepstroud.

## 1 Methods

### 1.1 Experimental materials and methods

Experimental methods have been described before^48^ and largely followed those used in our previous publications^10,80,81^. We briefly summarize the methods below.

#### 1.1.1 Subjects and apparatus

We used two female macaques (monkey T, *Macaca mulatta*, 5 kg; monkey K, *Macaca fuscata*, 8 kg). Both monkeys were housed individually. The light/dark cycle was 12/12 hr. (light, from 8:30 a.m. to 8:30 p.m.). The monkeys sat quietly in a primate chair in a dark, sound-attenuated shield room. During both training and neural recording sessions, we restrained the monkeys’ head movement non-invasively using a thermoplastic head cap as described in^82^. This head cap is made of a standard thermoplastic splint material (MT-APU, 3.2 mm thick, CIVCO Radiotherapy, IA., USA), and was molded out so that it conformed to the contours of the animals’ scalp, cheek bone, and occipital ridge. Visual stimuli were presented on a 17 inch TFT monitor placed 50 cm from the monkeys’ eyes. Eye movements were sampled at 120 Hz using an infrared eye tracking system (ETL-200, ISCAN, MA.). Eye fixation was controlled within a 6.5^*°*^ imaginary square window. TEMPO software (Reflective Computing, WA.) was used to control behavioral tasks. All experimental procedures were approved by the Animal Research Committee at the Graduate School of Frontier Biosciences, Osaka University, Japan and were in full compliance with the guidelines of the National BioResource Project ‘Japanese Macaques’. Experimental work performed in non-human primates that was not funded by Wellcome may not adhere to the principles outlined in the NC3Rs guidance on Non-human Primate Accommodation, Care and Use.

#### 1.1.2 Behavioral task

The monkeys were trained on a memory-guided saccade task requiring them to remember the location of a visual stimulus cue on a screen and to make a correct eye movement after a delay period (Fig. 1a). Specifically, this task required monkeys to fixate on a central ring for a period of 2.6–7.4 s followed by a stimulus cue (a white square) appearing in one of six pre-determined locations for 0.25 s. After a variable delay period of 1.4–7.5 s, the fixation ring was replaced by placeholders at all six possible stimulus cue locations (go cue). Monkeys were required to make a saccade within 0.5 s to the placeholder where the original stimulus cue was presented and maintain their gaze for 0.25 s for monkey T and either 0.25 s or 0.6 s for monkey K (these two gaze maintenance times were switched in different blocks for monkey K) to receive a juice reward. The monkeys were extensively trained, with close to perfect performance (monkey T, 96.1%; monkey K, 96.3%, mean across sessions). Fixation break errors were excluded from the calculation of percent correct rate.

#### 1.1.3 Recordings

After training was completed, we conducted an aseptic surgery under general anesthesia. We stereotypically implanted a plastic recording chamber on the lateral surface of the prefrontal cortex, under the guidance of structural MRI images (Extended Data Fig. 9). In monkey T, we implanted a cylindrical chamber (RC-T-S-P, internal diameter 12.7 mm, Gray Matter Research, MT.) in the right hemisphere (AP = 33, ML = 14.5; AP, anteriorposterior; ML, medio-lateral). A 32-channel semi-chronic microdrive system (SC-32, Gray Matter Research) was mounted inside this chamber. In monkey K, we implanted a cuboid chamber (width 12 mm, depth 16 mm, height, 15 mm, S-company ltd., Tokyo, Japan) over the principal sulcus in the left hemisphere.

We collected neural data in a total of 48 daily sessions (21 in monkey T; 27 in monkey K). In monkey T, we used the 32-ch microdrive (SC-32) that housed 32 single-contact tungsten electrodes with inter-electrode spacing of 1.5 mm. In monkey K, we used a 32-ch linear microelectrode array (Plexon U-Probe, Plexon, TX.) with an interelectrode spacing of 150 *µ*m along a single shaft. We positioned the U-Probe by using a custom-made grid (width 12 mm, depth 16 mm, height, 10 mm) which had a total of 165 holes with 1 mm spacing. We advanced the U-Probe by a custom-made hydraulic microdrive (S-company ltd.).

Raw extracellular neural signals were amplified and recorded in reference to a titanium bone screw at the vertex (in monkey T) or the shaft of the linear array (monkey K) using a neural signal amplifier RZ2 Bioamp Processor (Tucker-Davis Technologies, FL.). Behavioral data (task-event information and eye-movement information) were also sent to the RZ2 Bioamp. Neural data acquisition was performed at a sampling frequency of 24414.08 Hz, and behavioral data acquisition at 1017.25 Hz. For analysis of spiking activity, the raw neural signal was filtered (300 Hz to 6 kHz) for offline sorting (Offline Sorter, Plexon). In monkey T, approximately three hours before each recording session, we took the monkey to the testing room and advanced each electrode in the SC-32 by a minimum of 62.5 *µ*m in order to ensure recording of new neurons. We then put the monkey back in the home cage until we brought it out again for the recording session. In monkey K, we adopted the method of the U-Probe insertion reported in^83^. We first punctuated the dura using a guide tube (a shortened 23 gauge needle), and inserted the U-Probe array slowly, usually with a step of 500 *µ*m. We kept monitoring electrocardiogram (pulsatory fluctuation) on superficial electrodes to identify the point of cortical entry. We usually left 3–5 superficial channels outside the cortex. After array insertion, we waited 1–1.5 hours until the recorded single-unit and multiunit activities indicated that the electrode array was stably positioned in the cortex. While waiting, the monkey watched nature and animal video clips and received a small snack on a monkey chair.

In monkey T only, to determine location of the frontal eye field (FEF), and confirm that our recording area was outside it, intracortical microstimulations (22 biphasic pulses, 0.2 ms duration at 333 Hz, ≤ 150 *µ*A) were applied through microelectrodes. When eye movements were elicited below 50 *µ*A, the site was considered to be in the low-threshold FEF. In monkey T, our recording area did not include the low-threshold FEF.

#### 1.1.4 Pre-processing

We excluded neurons that were recorded in fewer than 10 trials for any cue condition. For each monkey, we pooled neurons from all recording sessions to create pseudopopulations of 438 neurons for Monkey K (after we removed 1 neuron from monkey K’s dataset due to an insufficient number of trials) and 625 neurons for Monkey T (no neurons were removed from monkey T’s dataset). To compute neural firing rates, we convolved spike trains with a Gaussian kernel with a standard deviation of 25 ms. Trial-averaged trajectories of time-varying mean firing rates were computed separately for each neuron and each cue condition. For analysis methods that used crossvalidation (see below), we split trials into separate train and test sets with a 1:1 train:test ratio, and computed trial-averaged trajectories for each training and test set (using 1:1 splits). For non-cross validated analyses, we either computed trial averages based on all the data, or on a subset of the data (see below). We aligned neural activity to either stimulus or go cue onset (see also below in Methods 1.7) and shifted activity by -50 ms to allow for the delay in time for information about these cues to enter PFC. For consistency with our simulations (see below), we subsampled neural firing rates at a 1-ms time resolution.

### 1.2 Neural network models: overview

All our simulated networks (Figs. 2–4 and 6 and Extended Data Figs. 2–5, 7 and 11–13) evolved according to a canonical model of stochastic recurrent neural circuit dynamics^40,84^:

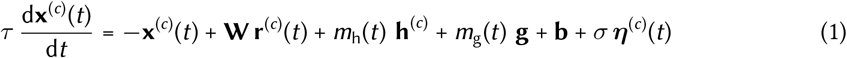

with

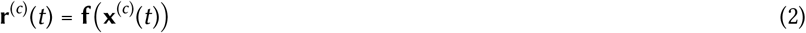

where 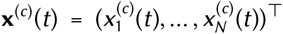 corresponds to the vector of (unitless) trial-averaged raw somatic membrane potentials of the *N* neurons of the network^84^ in cue condition *c* = 1, …, *C* (initialised at **x**^(*c*)^(*t*_0_) at the beginning of the simulation *t*_0_, which could be at or before stimulus onset at *t* = 0). **r**^(*c*)^(*t*) is their momentary firing rates, with **f**(**x**) being the activation function that converts membrane potentials to firing rates, *τ* is the effective time constant of the cell), **W** is the recurrent weight matrix (shown e.g. in Fig. 2a and g), **h**^(*c*)^ is the input given to the network depending on the stimulus cue, **g** is the stimulus-cue-independent go cue that occurs at the go time *t*_go_, *m*_h_(*t*) and *m*_g_(*t*) are box car ‘masking’ kernels such that the stimulus and go cues are only effective within a limited period at the beginning and end of the trial, respectively, **b** is a cue-independent bias, *σ* is the standard deviation of the noise process, and ***η***^(*c*)^(*t*) is a sample from a standard (mean 0 and variance 1) Gaussian white (temporally and spatially) noise process.

Networks shown in different figures corresponded to different special cases of Eqs. 1 and 2 (see Table 1). Specifically, for linear networks **f**(**x**) = **x** was the identity function. For nonlinear networks *f*_*i*_(**x**) = [*x*_*i*_]_+_ was the rectified linear (ReLU) activation function applied element-wise (except for the ring attractor networks where we used *f*_*i*_(**x**) = tanh(*x*_*i*_); Extended Data Fig. 3). Given that the focus of our study was optimal information loading, stimulus inputs were either optimized numerically (Fig. 2, Fig. 6, Extended Data Figs. 2 and 3, Extended Data Fig. 5c,d, and Extended Data Figs. 11–13), or set to analytically computed values as dictated by our mathematical analysis (Figs. 3 and 4, Extended Data Fig. 4c,d, and Extended Data Fig. 5a,b), or as a baseline, set to random values (Figs. 3 and 4, Extended Data Fig. 4a,b, and Extended Data Fig. 5a,b). For networks used to study the effects of instantaneous initial conditions (Figs. 2 and 3 and Extended Data Figs. 2–5 and 7), the stimulus masking kernel was zero and instead the initial condition was set to the stimulus input; for other networks (Fig. 4, Fig. 5e–f, Fig. 6 and Extended Data Figs. 11–13) the stimulus masking kernel was a boxcar between 0 and 0.25 s. For task-optimized networks (Fig. 6 and Extended Data Figs. 11–13), the go cue masking kernel *m*_g_(*t*) was a boxcar starting at go cue onset, *t*_go_ and lasting for 0.5 s, for all other networks it was set to 0 everywhere. The networks used to analyse the dynamics of information loading (Fig. 3, Extended Data Fig. 4, and Extended Data Fig. 5a,b) were deterministic by setting *σ* = 0, all other networks used noisy dynamics (see Table 1). We solved the dynamics of Eqs. 1 and 2 using a first-order Euler–Maruyama approximation between *t*_0_ and the simulation end time, *t*_max_, with a discretization time step of 1 ms.

**Table 1.**
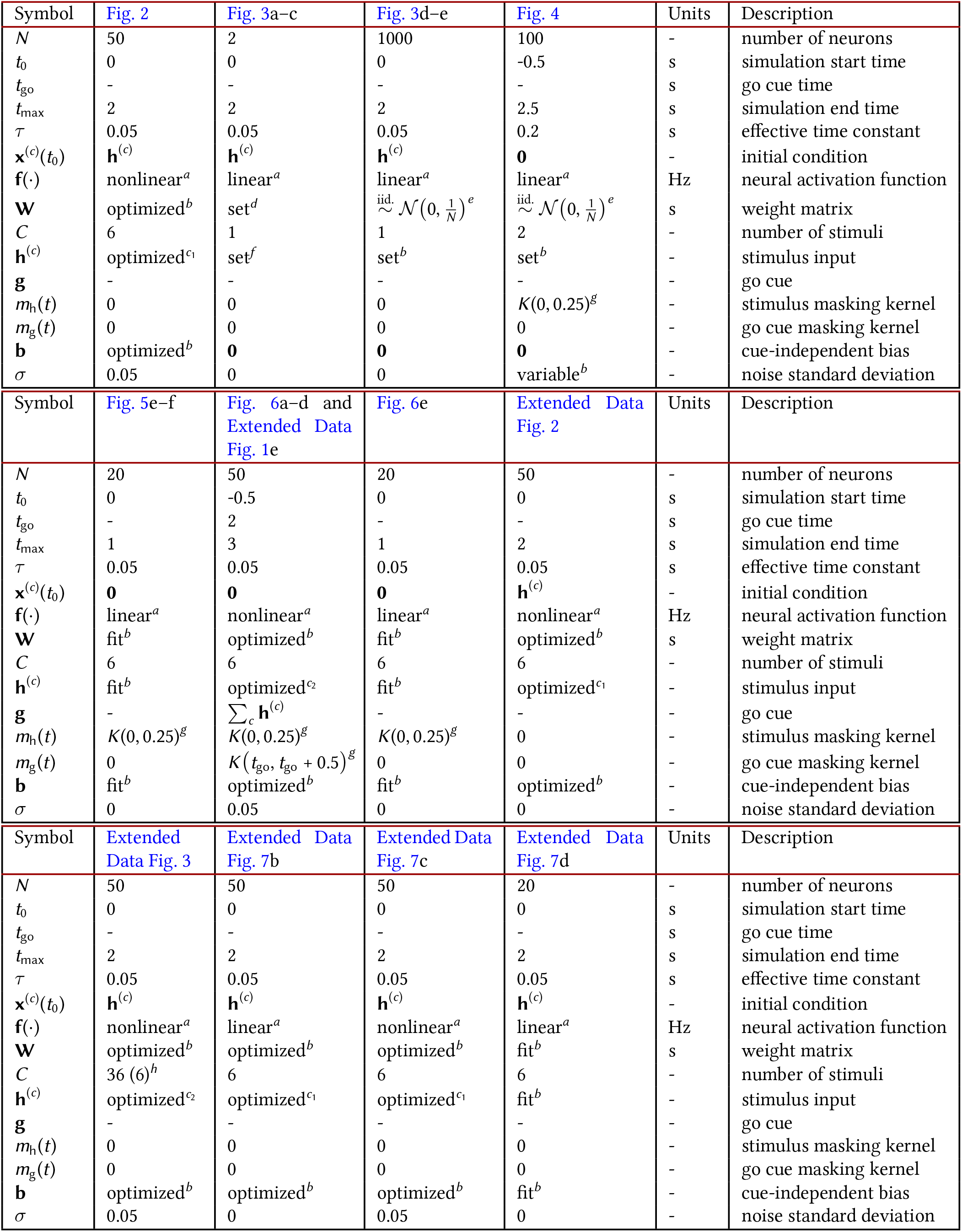
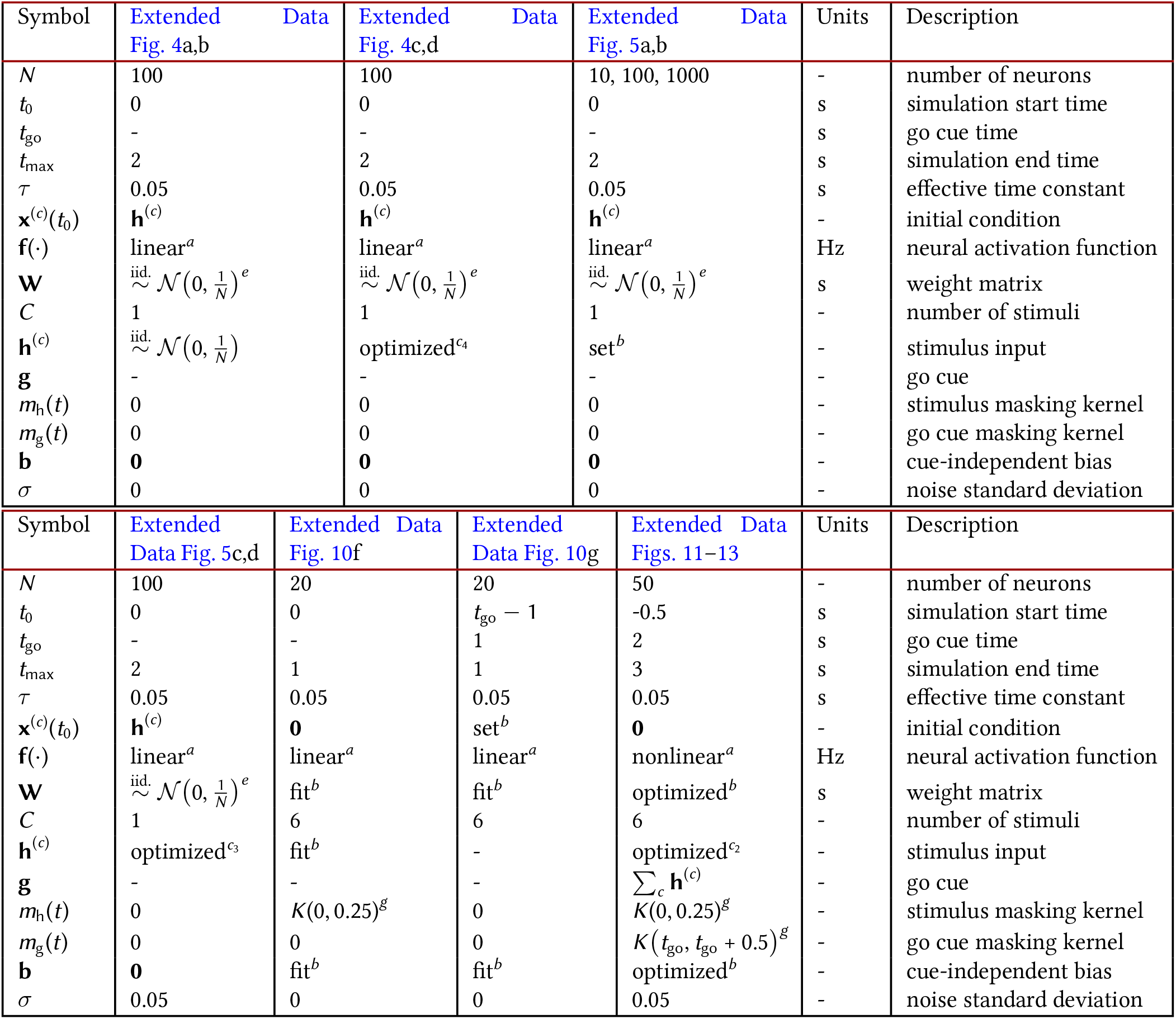
Parameters used in the simulations of our models. ^*a*^ For nonlinear networks, *f*_*i*_(**x**) = [*x*_*i*_]_+_ was the rectified linear (ReLU) activation function. For linear networks *f*_*i*_(**x**) = *x*_*i*_. The only exception to this was when we created ring attractor networks (Extended Data Fig. 3) in which we used a tanh nonlinearity *f*_*i*_(**x**) = tanh(*x*_*i*_). See also text. ^*b*^ See text for details. ^*c*^ Inputs were optimized either with both a norm constraint and an overall energy constraint (*c*_1_); only an overall energy constraint (*c*_2_); only a norm constraint (*c*_3_); or so that the dynamics produced the mathematically minimal overall energy (*c*_4_, see Supplementary Information S2). See text for more details. ^*d*^ For the symmetric network, we used 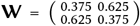; for the unconstrained network, we used 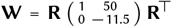, where **R** is the rotation matrix 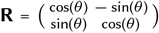 with *θ* = 0.1. ^*e*^ For the symmetric networks, we enforced 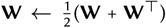. For all networks we also shifted the obtained weight matrix by the identity matrix multiplied by a constant so that the largest real part in the eigenvalues of **W** is exactly 1 (i.e., the largest eigenvalue of the associated Jacobian would therefore be 0 due to the leak term), and we rejected any **W**’s for which the top eigenvalue (the eigenvalue with largest real part) had an imaginary component. For Fig. 4, to provide a slightly better agreement between the model dynamics and the experimental recordings, we rejected any **W**’s for which the inner product between the most amplifying mode and persistent mode was greater than 0.2 (i.e. we only kept **W**’s that were relatively mathematically non-normal). ^*f*^ We used 3 possible input directions (which all had a Euclidean norm of 1): inputs either aligned with the most persistent mode (**x**^p^), the most amplifying mode (**x**^a^), or a random direction (**x**^r^). For the symmetric model, **x**^p^ = **x**^a^ = [0.707, 0.707]^⊤^ and we used **x**^r^ = [0.98, 0.18]^⊤^. For the unconstrained model, **x**^p^ = [0.995, 0.0998]^⊤^, **x**^a^ = [0.1537, 0.9881]^⊤^ and we used **x**^r^ = [0.8453, 0.5343]^⊤^. ^*g*^ 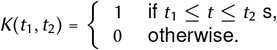 In the table, *t*_go_ refers to the timing of the go cue (see text). ^*h*^ For training, we used *C* = 36 cue conditions. For our subsequent analyses (Extended Data Fig. 3), we used *C* = 6 cue conditions to be consistent with the other models.

For analysis methods that used cross-validation (see below), for each cue condition, we simulated network dynamics twice with independent realizations of ***η***^(*c*)^(*t*), to serve as train and test data. For other analyses, we used a single set of simulated trajectories. All analyses involving networks with randomly generated (or initialized) connectivities that also did not require re-fitting their responses with other networks (Fig. 2, Fig. 4, Fig. 6c,d, Extended Data Figs. 1–3, Extended Data Fig. 7a–c, and Extended Data Figs. 11–13) were repeated a total of *n* = 100 times, consisting of 10 different networks and 10 different simulations. For those analyses that did require the re-fitting of nonlinear networks’ responses with linear deterministic networks (Extended Data Fig. 7d and Fig. 6e), we used one simulation of the original (stochastic nonlinear) network, so *n* = 10 simulations in total.

### 1.3 Nonlinear networks

For the dynamical equations of nonlinear networks, see Methods 1.2. For nonlinear networks (Figs. 2 and 6, Extended Data Figs. 2 and 3, Extended Data Fig. 7c, and Extended Data Figs. 11–13), we ensured that they performed working memory maintenance competently by optimizing their free parameters, **W, b**, and **h**^(*c*)^ for appropriate cost functions (see below, Methods 1.3.3).

#### 1.3.1 Nonlinear networks with instantaneous inputs

Following classical theoretical approaches to attractor network dynamics, we first used nonlinear neural networks in which stimulus inputs acted instantaneously to determine the initial conditions of the dynamics Fig. 2 and Extended Data Figs. 2, 3 and 7. These networks were optimized using a ‘just-in-time’ cost function (Methods 1.3.3) under one or two constraints. First, for all these networks, we constrained stimulus inputs to have a Euclidean norm of 3 so that we could compare information loading strategies fairly when inputs were constrained to lie in certain subspaces (see also below): either the persistent subspace, persistent nullspace, locally persistent subspace, locally most amplifying subspace, or a random subspace (Fig. 2f,l, Extended Data Fig. 2, and Extended Data Fig. 7).. We also obtained qualitatively very similar results without this norm constraint, with only a more general energy-based penalty^13,39,40,65^ (Fig. 6 and Extended Data Figs. 3 and 11–13, see also Methods 1.3.3). Second, for symmetric networks, we enforced 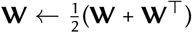.

These networks were trained in two epochs. For the first 1000 training iterations, we optimized all free parameters. After this, we confirmed that our trained networks did indeed have attractors (i.e. that they were attractor networks) and determined where these attractors were in state space by finding the stable fixed points of the networks’ dynamics (Fig. 2d,j, Extended Data Fig. 2b,f, and Extended Data Fig. 3a)—see below. We then continued for another 1000 training iterations (without any firing rate regularization, 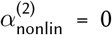 in Eq. 3) with only optimizing the initial conditions, **h**^(*c*)^, while keeping the other parameters, **W** (Fig. 2a,g) and **b**, fixed at the values obtained at the end of the first 1000 iterations. We did this so that we could fairly compare different initial conditions that are constrained to lie in different subspaces but which otherwise rely on the same underlying network dynamics. We considered three possible scenarios for introducing additional constraints on the initial conditions (beside the one on their norm, see above): they were either projected and then restricted to the persistent subspace or to the persistent nullspace (see Methods 1.7.1 for how these subspaces were computed), or there was no such constraint applied so that they could utilize any direction in the full state space of the network. In addition, to understand the link between the linearized (Methods 1.4.2) and original (above) forms of the dynamics of these networks, we also considered three more constraints on the initial conditions: constraining them to the most persistent, most amplifying, or a random subspace of the linearized dynamics (Extended Data Fig. 7a–c).

#### 1.3.2 Nonlinear networks with temporally extended inputs

To more closely follow the experimental paradigms which we modelled, we also used nonlinear networks in which stimuli provided temporally extended inputs (Fig. 6 and Extended Data Figs. 11–13). To construct these networks, stimulus inputs and the weight matrix were freely optimized (Methods 1.3.3), without any constraints, and optimization proceeded for a full 2000 iterations, without dividing training into different epochs.

#### 1.3.3 Cost functions and training for nonlinear networks

To investigate how different cost functions impact network dynamics (Fig. 6), we trained networks using one of four cost functions: a ‘cue-delay’ cost, a ‘full-delay’, a ‘just-in-time’ cost, and an ‘after-go’ cost. These costs only differed in terms of the time period in which we applied the cost function. The general form of the cost function we used was a cross entropy loss plus a regularisation term:

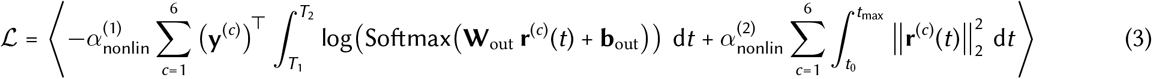

where *T*_1_ and *T*_2_ determine the time period in which we applied the cost, 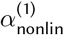 and 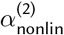 control the relative contributions of the cross-entropy loss and firing rate regularisation, **W**_out_ *∈ ℛ*^6*×N*^ and **b**_out_ *∈ ℛ*^6^ include the 6 sets of ‘readout’ weights and biases, respectively, and **y**^(*c*)^ *∈ ℛ*^6^ is a one-hot vector where 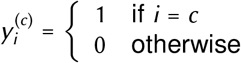 defining the ‘target’ output for each cue condition. We initialized elements of the network parameters **W, b, h**^(*c*)^, as well as the readout parameters **W**_out_ and **b**_out_ from a Gaussian distribution with mean 0 and variance 1*/N*, and then optimized using gradient descent with Adam optimization^85^, where gradients were obtained from backpropagation through time. The angle brackets,⟨ ⟩, denote averaging over batch sizes of 50 random realisations of **r**^(*c*)^. We used a learning rate of 0.0005.

See Table 2 for how we set the parameters of Eq. 3 (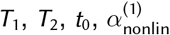 and 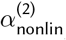) depending on the cost function and the level of regularization. Briefly, the cue-delay cost included both the cue (between stimulus cue onset and offset) and the delay period (between stimulus cue offset and go cue onset), the full-delay cost the included delay period but not the cue period, the just-in-time cost started between stimulus onset and the earliest go time and ended at the onset of the go cue, and the after-go cost started at go cue onset and lasted for the duration of the go cue (0.5 s). For simulating the random delay task (Fig. 6 and Extended Data Figs. 11–13), analogous to what animals need to solve (see below), we sampled the go time uniformly between *t*_go_ = 0.75 s and *t*_go_ = 2 s. For just-in-time (Fig. 2 and Extended Data Fig. 2) and after-go trained networks (Extended Data Fig. 12), we also used a fixed delay task with a simulation end time of *t*_max_ = 2 s or a go time of *t*_go_ = 2 s, respectively. For the other cost functions, networks trained on the fixed delay task yielded very similar dynamics to their counterparts trained on the variable delay task (not shown). We set 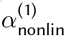 and 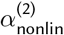 so that networks could reliably learn the task (at performance levels comparable across different settings) while also exhibiting relatively stable dynamics (i.e. if 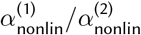 is too large, the network dynamics can explode whereas if 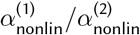 is too small, the network is not able to learn the task). Note that vanishing gradients during training also impacted the value of 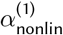 that was required for different networks to exhibit similar performance (Fig. 6c). Nevertheless, 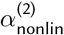 was varied by an order of magnitude between Fig. 6 and Extended Data Figs. 11–13 to specifically test the robustness of our results to this parameter.

**Table 2.**
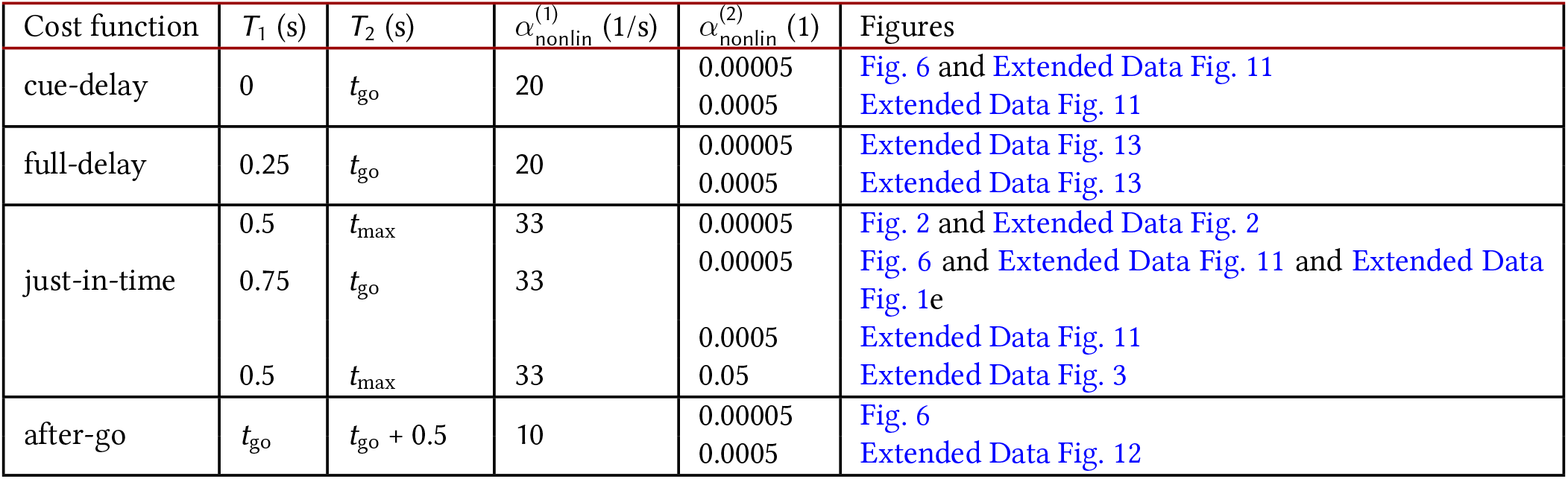
Parameters for nonlinear network optimization. Times *T*_1_ and *T*_2_ are relative to stimulus onset at *t* = 0. Units are shown in parentheses after the name of the corresponding parameter.

#### 1.3.4 Optimized ring attractor networks

When training to create ring attractor networks (Extended Data Fig. 3), we made three modifications to the nonlinear networks described above. First, in line with other approaches for optimizing recurrent neural networks^17,20,24,65^, we used a hyperbolic tangent nonlinearity because the saturation of this nonlinearity greatly encouraged a continuous attractor to form compared with a ReLu nonlinearity. Second, we trained networks with 36 cue conditions, and then subsequently restricted our analyses to 6 evenly spaced cue conditions to keep consistency with our other analyses. Third, we used a cost function that measured estimation (or fine, rather than coarse, discrimination) performance across those 36 conditions, thus encouraging a ring attractor to form:

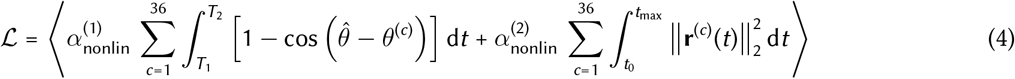

with

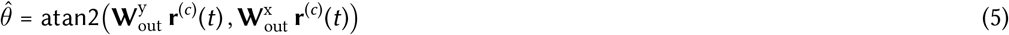

where 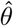 is the population vector-decoded stimulus angle, such that atan2(*y, x*) gives the angle that the vector [*x, y*] makes with the x-axis, 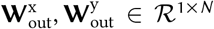 are 2 sets of ‘readout’ weights defining the plane in which decoded angles are defined, and 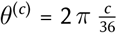 is the target angle for cue condition *c*. All other terms were the same as those defined in Methods 1.3.3. See also Supplementary Information S5 for a derivation showing how the cost function Eq. 4 relates to Fisher information.

### 1.4 Linear networks

For the dynamical equations of linear networks, see Methods 1.2. Linear networks were either constructed ‘*de novo*’ (Figs. 3 and 4 and Extended Data Figs. 4 and 5), obtained by a local linearization of canonical nonlinear dynamical systems (Extended Data Fig. 6) or of nonlinear neural network dynamics (Extended Data Fig. 3e and Extended Data Fig. 7a–c), or they were fitted to neural responses obtained from experiments (Fig. 5f, and Extended Data Fig. 10f,g) or the simulation of nonlinear networks (Fig. 6e and Extended Data Fig. 7d).

#### 1.4.1 De novo linear networks

We used *de novo* linear networks to develop an analytical understanding of the dynamics of optimal information loading. These networks included small 2-neuron networks with hand-picked parameters (see Table 1) chosen to illustrate the differences between normal (symmetric) and non-normal (unconstrained) dynamics and the effects of different initial conditions (Fig. 3a–c), as well as large networks (with 10, 100, or 1000 neurons) with randomly generated parameters (Fig. 3d,e, Fig. 4, Extended Data Fig. 4, and Extended Data Fig. 5a,b; see Table 1). We always set the largest eigenvalue of the weight matrix to be exactly 1 (thus setting the largest eigenvalue of the associated Jacobian to 0 due to the leak term) so that these networks had an integrating or ‘persistent’ mode^6,26,34,35^ (see Table 1)

Initial conditions (Fig. 3, Extended Data Fig. 4, and Extended Data Fig. 5a,b) or temporally extended inputs (Fig. 4) were determined by computing the most persistent and amplifying direction(s) based on the Jacobian of the dynamics (Figs. 3 and 4, and Extended Data Fig. 5a,b, see Methods 1.7.1; for how initial conditions were determined in Extended Data Fig. 4c,d see Supplementary Information S2.8). For the networks in Fig. 4, we also added a small amount of noise to the input to allow for some transient dynamics for all input directions (see Fig. 4c at 0 s). Alternatively, we optimized initial conditions for maximal asymptotic overlap with the most persistent mode (Extended Data Fig. 5c,d; see below). For setting the noise level, *σ*, in these networks, we considered two scenarios: noise matched (Fig. 4a, light green and gray) and performance matched (Fig. 4a, dark green and black). For noise matched simulations, we first determined the highest value of *σ* that still allowed us to obtain 100% decodability (using a delay-trained decoder) for all networks when receiving inputs aligned with the most amplifying mode (Fig. 4a, red). This resulted in *σ* = 0.1 for symmetric models, and *σ* = 0.17 for unconstrained models. We then used the same *σ* for simulations using inputs aligned with the most persistent and random directions. For performance matched simulations, we used a different value of *σ* for each possible input direction so that all models achieved 100% decodability using a delay-trained decoder. For symmetric models, this required *σ* = 0.1 for inputs aligned with either the persistent or most amplifying modes, and *σ* = 0.005 for random inputs. For unconstrained models, this required *σ* = 0.17 for inputs aligned with the most amplifying mode, *σ* = 0.02 for inputs aligned with the persistent mode, and *σ* = 0.005 for random inputs. (Note that, consistent with our theory, smaller noise levels were necessary to achieve the same desired level of performance for input directions that were predicted to be increasingly suboptimal by our analysis.)

To demonstrate that the initial conditions along the most amplifying directions, obtained by control theoretic analyses, were indeed optimal for maximising the overlap with the most persistent mode (the measure of optimality we used for these networks, Fig. 3c,e), we also used a direct numerical optimization approach, analogous to that used to optimize initial conditions in our nonlinear networks (Figs. 2 and 6, see also Methods 1.3.3). Specifically, we optimized **h**^(*c*)^ (constrained to have unit Euclidean norm) with gradient descent using Adam optimization^85^ with gradients obtained from back-propagation through time using the following cost function

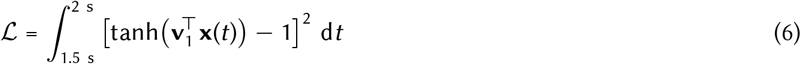

where **v**_1_ is the eigenvector associated with eigenvalue 0 of the Jacobian (i.e. the most persistent mode). We used a learning rate of 0.0001. We performed the above training procedure independently for 100 random noisy networks (either symmetric or unconstrained) and we show averaged results in Extended Data Fig. 5c,d. We also used random initial conditions as controls. These had elements that were either sampled from a standard normal distribution (re-scaled to have unit Euclidean norm) in large networks (Fig. 3d,e, Fig. 4, and Extended Data Fig. 5a,b), or in the case of 2-neuron networks, quasi-randomly chosen (with unit Euclidean norm) for illustrative purposes (Fig. 3a–c).

#### 1.4.2 Local linearization of nonlinear dynamics

To better understand how the dynamics of optimal information loading that we identified in linear networks apply to nonlinear attractor dynamics, we performed a local linearization of our simulated nonlinear networks (Extended Data Fig. 3e, Extended Data Fig. 6, and Extended Data Fig. 7a–c). This approach required access to the ‘true’ dynamical equations of the nonlinear networks—which we had by construction.

We performed local linearizations of the original nonlinear network dynamics in **x**-space (the space of variables in which the dynamics was defined, Eq. 1) around the origin (we found empirically that initial conditions were distributed close to the origin)—which served as the reference point with respect to which the norm of optimized initial conditions was constrained in the networks we linearized (Methods 1.3; analogous to our analysis of information loading in linear networks, Fig. 3a–e, and see also Methods 1.7.2). As the ReLU firing rate nonlinearity of these networks is non-differentiable at exactly the origin, we computed the ‘average’ Jacobian of the system in the immediate vicinity of the origin instead (this allowed us to use the same linearization and the same set of amplifying modes for all initial conditions; we obtained similar results by linearizing separately for each initial condition). Because the derivative of each ReLU is 0 or 1 in half of the activity space of the network, this resulted in the Jacobian 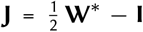, where **W**^*\**^ is the weight matrix of the original nonlinear network. Note that one obtains the same result even without averaging, by regarding the ReLu nonlinearity as the limiting case of the soft-ReLu nonlinearity: 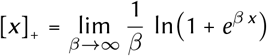, of which the derivative at *x* = 0 is 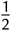 (at any value of the inverse temperature, *β*) and thus results in the same Jacobian as above. We confirmed that the resulting dynamics were always stable (largest real eigenvalue of **J** was less than 0). We then used this system to identify the locally (around the origin) most amplifying or most persistent modes (Extended Data Fig. 7a).

For simulating these linearized networks (Extended Data Fig. 7b), we then used the Jacobian we thus obtained to map the resulting linearized dynamics to a deterministic integrator with the effective weight matrix **W** = (**J** + **I**) *− λ*_max_ **I**, where *λ*_max_ is the largest real eigenvalue of **J**. Thus, the resulting dynamics were always marginally stable (largest real eigenvalue of **J** was exactly 0). (Note that for subsequent analyses involving most persistent and amplifying modes, we used the original weight matrix, see more in Methods 1.7.1. Nevertheless, the most persistent modes of the weight matrices we used for simulation and those we used in subsequent analyses were identical, as they only relied on the eigenvectors of the weight matrix, or the Jacobian, and the rank order of their associated eigenvalues, which this stabilization did not affect. We also checked numerically that making the system marginally stable only had very minor effects on the most amplifying modes, with correlations between the most amplifying modes of the original and simulated dynamics being above 0.9. Thus, in these respects, our simulations were representative of the dynamics of the original systems.) The bias parameters, **b**, were the same as in the original nonlinear networks. The initial conditions, **h**^(*c*)^, were either the ones we originally optimized for the nonlinear dynamics without any constraints (beside a constraint on their norm), or they were optimized while constraining them to the most persistent, most amplifying, or a randomly chosen subspace of these linearized dynamics (all were of the same dimensionality for a fair comparison, Extended Data Fig. 7).

For ring attractor networks (Extended Data Fig. 3), which used a tanh nonlinearity (Methods 1.3.4), the associated linearized system around the origin was given by the Jacobian **J** = **W** *−***I**, which we then used to identify the locally most amplifying and persistent modes (Extended Data Fig. 3e).

We used the same approach to linearize the dynamics of the canonical minimal nonlinear attractor dynamics that we used to gain insights into information loading in nonlinear systems (Methods 1.6, see also Supplementary Information S3 and Extended Data Fig. 6). In this case, the Jacobian was well defined at the origin, so there was no need to average it. For consistency with the notation and terminology we use in the rest of this paper, and without loss of generality (as linear dynamical systems and linear neural networks are isomorphic), we refer to the resulting linear dynamical system as a ‘linear neural network’ and define it by its ‘effective’ weight matrix (defined via the Jacobian as above). Initial conditions were magnitude-matched and chosen to align with the most persistent or the most amplifying direction extracted from the Jacobian (Methods 1.7.1), or chosen randomly, or varied systematically to cover the whole range of possible directions. There were no other parameters for these linearized ‘networks’.

#### 1.4.3 Fitting linear neural networks to neural responses

In order to be able to apply our theoretically derived measures of optimal information loading without having access to the true dynamics of the system, we also created linear neural networks whose parameters were fitted to experimental data (see below). As a control, we repeated the same fitting procedure with simulated nonlinear networks to validate that our approach provides meaningful results when 1. we do not have access to the true dynamics but only to samples of activities generated by those dynamics, and 2. we also cannot assume that the true dynamics are linear.

We fitted deterministic linear neural networks to 1 s of trial-averaged neural activity (experimentally recorded, or simulated by a nonlinear neural network model). For the main analyses (Fig. 5d–f, Fig. 6e, and Extended Data Fig. 7d), we used data starting from the onset of the stimulus cue. For the control analysis of late delay experimental recordings (Extended Data Fig. 10g), we used the final 1 s of neural activity just prior to the go cue. For the shuffle control (Fig. 5d; dark gray, and Extended Data Fig. 10f), we again used data starting from stimulus onset but randomly shuffled neural activity across time and proceeded by fitting this shuffled data instead.

For fitting high dimensional neural data, we first performed principal components analysis on neural activity (dimensions: neurons, data points: time points, indexed by *t*, and cue conditions, indexed by *c*), and projected it through the principal components 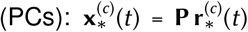, where the columns of **P** are top 20 principal components of the data, and 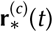 is trial averaged neural responses (mean-centered, see above) at time *t* in condition *c*. These top 20 PCs captured approximately 75% and 76% of variance for monkeys K and T, respectively during the cue and early delay period (Fig. 5d–f), 70% and 60% of variance for monkeys K and T, respectively during the late delay period (Extended Data Fig. 10g), and over 95% of the variance for all simulated neural activities (Fig. 6e). The projected neural activity time courses of the neural data, 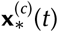, served as the targets that needed to be matched (after a suitable linear transformation with ‘read-out’ matrix **C** *∈ ℛ*^20*×*20^) by the neural activity time courses generated by the fitted neural network’s dynamics in the corresponding cue conditions, **x**^(*c*)^(*t*) (Eqs. 1 and 2). For fitting the parameters of the network (**W, h**^(*c*)^, **b**) and the readout matrix (**C**), we used the following cost function:

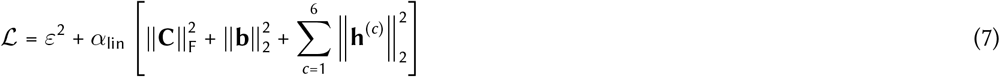

with

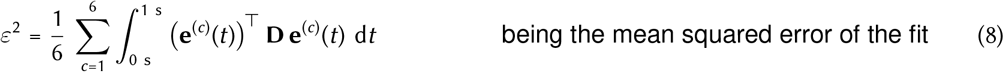

and

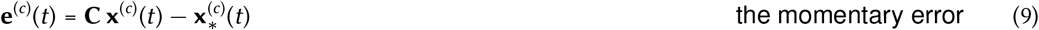

where **D** is a diagonal matrix with the variances explained by the corresponding PCs in **P** on the diagonal (encouraging the optimization procedure to prioritize fitting the top PCs),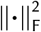 is the Frobenius norm of a matrix.

Also note that we had no constraints on **W** to define stable dynamics. Nevertheless, when fitting experimental recordings, and responses generated by nonlinear attractor networks, we found that the largest real eigenvalue of the fitted **W** was typically within the 0.95 *≤λ*_max_ *≤*1.05 range, i.e. the dynamics were near marginal stability, in line with the dynamics of our *de novo* linear neural networks (Methods 1.4.1), as well as of those that we obtained by local linearization (Methods 1.4.2). The only exception was when fitting the responses of nonlinear networks trained on an after-go-time cost (Methods 1.3.3) which resulted in dynamics without attractors and, consequently, the fitted linear dynamics typically had *λ*_max_ *>* 1.05.

We used Adam^85^ to perform gradient descent optimization of **W, h**^(*c*)^, **b**, and **C** with gradients obtained from backpropagation through time, and a learning rate of 0.0001. We initialized elements of all of these parameters from a Gaussian distribution with mean 0 and variance 1*/*20 and we set the regularisation parameter to *α*_lin_ = 1*/*12.

The stimulus-masking kernel (*m*_h_(*t*), Table 1) was matched to how the responses being fitted were obtained: with temporally extended or instantaneous inputs. Specifically, when fitting responses to temporally extended inputs (experimentally measured, Fig. 5f and Extended Data Fig. 10f, or simulated, Fig. 6e), the masking kernel of the fitted linear network matched the cue period. When fitting responses generated by networks driven by instantaneous inputs (Extended Data Fig. 7d), or when fitting the late delay period of experimental recordings (during which no stimulus is present, Extended Data Fig. 10g), the stimulus masking kernel was set to zero, and instead the initial condition of the fitted linear network was tuned to match the responses (see below).

In most cases (Fig. 5f, Fig. 6e, and Extended Data Fig. 10f), we set the initial condition **x**^(*c*)^(*t*_0_) = **0**. There were two exceptions to this. First, when fitting the late delay dynamics in the experimental recordings (Extended Data Fig. 10g), we set 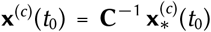 (i.e. we fixed the initial condition of the latent dynamics to the data as no stimulus is present during the late delay period; we also observed qualitatively similar results when we included **x**^(*c*)^(*t*_0_) as a separate optimizable parameter in this case). Second, when fitting simulated data from models that used instantaneous stimulus inputs (Extended Data Fig. 7d), we set **x**^(*c*)^(*t*_0_) = **h**^(*c*)^.

### 1.5 Previous working memory models

We used the following dynamics for implementing all previous neural network models of working memory:

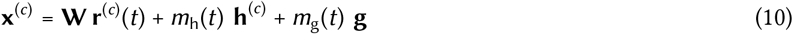

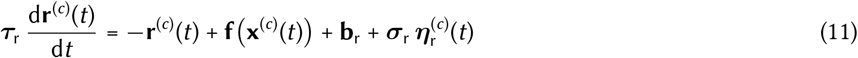

where all symbols refer to the same (or a closely analogous, see below) quantity as in Eqs. 1 and 2. Note that we use this notation to best expose the similarities with and differences from the dynamics of our networks (Eqs. 1 and 2), rather than the original notation used for describing these models^5,6,27^, but the dynamics are nevertheless identical to those previously published. Overall, these dynamics are closely analogous to those that we used earlier for our networks with the following differences. First, for us, dynamics were defined in **x**-space, with **r** being an instantaneous function of **x**. Here, the dynamics are defined instead in **r**-space (Extended Data Fig. 1a–d and Extended Data Fig. 8), with **x** being an instantaneous function of **r**. (There are slightly different assumptions underlying these rate-based formulations of neural network dynamics when deriving them as approximations of the dynamics of spiking neural networks^46^, and the two become identical in the case of linear dynamics.) As a result, time constants, ***τ***_r_, biases, **b**_r_, and the variance of noise, ***σ***_r_ (as well as noise itself, 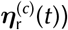, are defined for **r** rather than **x**. For nonlinear variants of these networks, there are also differences for the choice of single neuron nonlinearities, **f**(·). Furthermore, some of these networks distinguish between excitatory and inhibitory cells, with different time constants, and noise standard deviations. Thus, each of these parameters is represented as a diagonal matrix, ***τ***_r_ and ***σ***_r_, respectively, with each element on the diagonal storing one of two possible values of that parameter depending on the type (excitatory or inhibitory) of the corresponding neuron (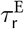 and 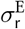, or 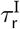 and 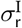 respectively). Most importantly, all of these networks used a set of parameters which were hand-crafted to produce the required type of dynamics, rather than optimized for a function (or to fit data) as in the case of our networks. In line with our analyses of experimental data and task-optimized networks (Figs. 5 and 6), simulations started at *t*_0_ = *−* 0.5 s, i.e. 0.5 s before stimulus cue onset (defined as *t* = 0), the stimulus cue lasted for 0.25 s, and the go cue appeared at *t*_go_ = 2 s and lasted for 0.5 s. (Note that for these networks we considered the fixed-delay variant of the task as that is what these networks were originally constructed to solve.) As with our networks (Methods 1.2), we solved the dynamics of Eqs. 10 and 11 using a first-order Euler–Maruyama approximation between *t*_0_ and the simulation end time with a discretization time step of 1 ms.

For analysis methods that used cross-validation (see below), we simulated network dynamics twice (for each cue condition) with independent realizations of 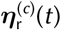, to serve as (trial-averaged) train and test data. For other analyses, we used a single set of simulated trajectories. All analyses involving these networks were repeated *n* = 10 times, using 10 different simulations (non-cross-validated) or simulation-pairs (cross-validated), each time with independent samples of 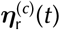.

Table 3 provides the values of most network and other parameters used for simulating each model. In the following we provide the additional details for each of these models that are not included in Table 3.

**Table 3.**
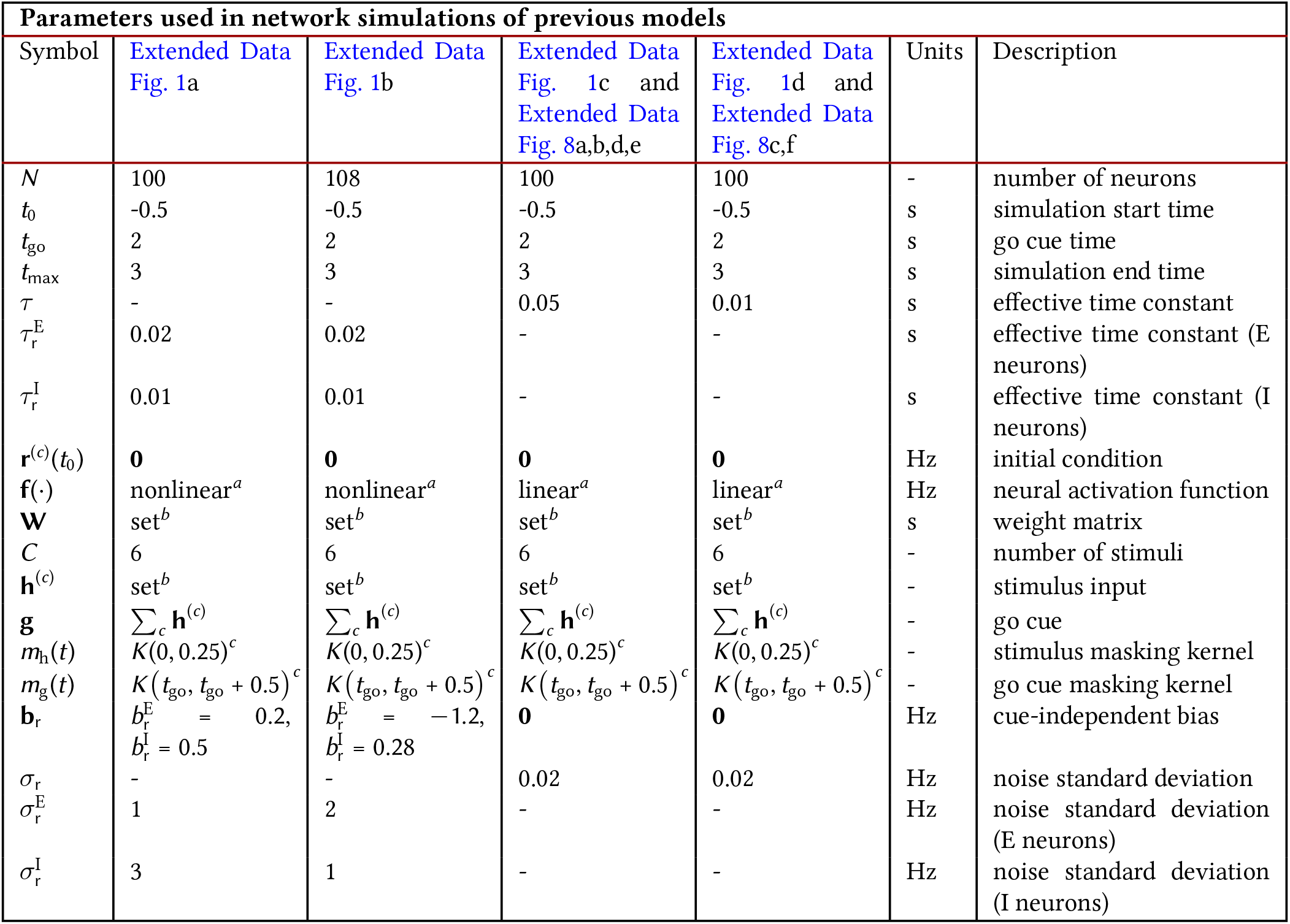
Parameters used in previous models. ^*a*^ For nonlinear networks, 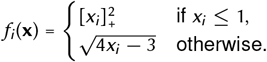. For linear networks *f* (**x**) = *x*_*i*_. ^*b*^ See text for details. ^*c*^ 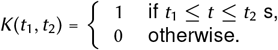

| Parameters used in network simulations of previous models |  |  |  |  |  |  |
| --- | --- | --- | --- | --- | --- | --- |
| Symbol | Extended Data Fig. 1a | Extended Data Fig. 1b | Extended Data Fig. 1c and Extended Data Fig. 8a,b,d,e | Extended Data Fig. 1d and Extended Data Fig. 8c,f | Units | Description |
| $N$ | 100 | 108 | 100 | 100 | - | number of neurons |
| $t_0$ | -0.5 | -0.5 | -0.5 | -0.5 | s | simulation start time |
| $t_{go}$ | 2 | 2 | 2 | 2 | s | go cue time |
| $t_{max}$ | 3 | 3 | 3 | 3 | s | simulation end time |
| $\tau$ | - | - | 0.05 | 0.01 | s | effective time constant |
| $\tau_r^E$ | 0.02 | 0.02 | - | - | s | effective time constant (E neurons) |
| $\tau_r^I$ | 0.01 | 0.01 | - | - | s | effective time constant (I neurons) |
| $\mathbf{r}^{(c)}(t_0)$ | $\mathbf{0}$ | $\mathbf{0}$ | $\mathbf{0}$ | $\mathbf{0}$ | Hz | initial condition |
| $\mathbf{f}(\cdot)$ | nonlinear<sup>a</sup> | nonlinear<sup>a</sup> | linear<sup>a</sup> | linear<sup>a</sup> | Hz | neural activation function |
| $\mathbf{W}$ | set<sup>b</sup> | set<sup>b</sup> | set<sup>b</sup> | set<sup>b</sup> | s | weight matrix |
| $C$ | 6 | 6 | 6 | 6 | - | number of stimuli |
| $\mathbf{h}^{(c)}$ | set<sup>b</sup> | set<sup>b</sup> | set<sup>b</sup> | set<sup>b</sup> | - | stimulus input |
| $\mathbf{g}$ | $\sum_c \mathbf{h}^{(c)}$ | $\sum_c \mathbf{h}^{(c)}$ | $\sum_c \mathbf{h}^{(c)}$ | $\sum_c \mathbf{h}^{(c)}$ | - | go cue |
| $m_h(t)$ | $K(0, 0.25)^c$ | $K(0, 0.25)^c$ | $K(0, 0.25)^c$ | $K(0, 0.25)^c$ | - | stimulus masking kernel |
| $m_g(t)$ | $K(t_{go}, t_{go} + 0.5)^c$ | $K(t_{go}, t_{go} + 0.5)^c$ | $K(t_{go}, t_{go} + 0.5)^c$ | $K(t_{go}, t_{go} + 0.5)^c$ | - | go cue masking kernel |
| $\mathbf{b}_r$ | $b_r^E = 0.2,$<br>$b_r^I = 0.5$ | $b_r^E = -1.2,$<br>$b_r^I = 0.28$ | $\mathbf{0}$ | $\mathbf{0}$ | Hz | cue-independent bias |
| $\sigma_r$ | - | - | 0.02 | 0.02 | Hz | noise standard deviation |
| $\sigma_r^E$ | 1 | 2 | - | - | Hz | noise standard deviation (E neurons) |
| $\sigma_r^I$ | 3 | 1 | - | - | Hz | noise standard deviation (I neurons) |

#### 1.5.1 Classical bump attractor model

The bump attractor model that we used (Extended Data Fig. 1a) has been described previously (see Ref. 5). The model contained separate excitatory and inhibitory populations. As in the discrete attractors model, the weight matrix was of the form

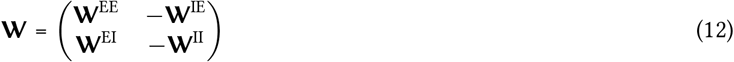

where the elements of **W**^IE^,**W**^EI^, and **W**^II^ were set to 6.8*/N*, 8*/N*, and 1.7*/N*, respectively. The excitatory sub-matrix **W**^EE^ had a circulant form:

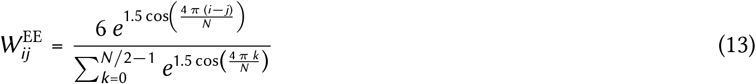

for cell-pairs *i, j* = 1, …, *N /*2.

Stimulus cue inputs were also analogous to those used in the discrete attractors models and were set to

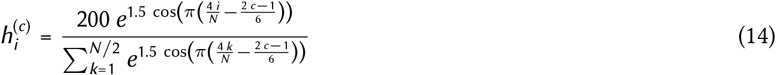

for cues *c* = 1, …, 6 and cells *i* = 1, …, *N /*2 (i.e., as above, inputs were only delivered to the excitatory neurons).

#### 1.5.2 Discrete attractors model

The discrete attractors model that we used (Extended Data Fig. 1b) has been described previously (see the methods of Ref. 5). The model contained separate excitatory and inhibitory populations.

The weight matrix was of the form

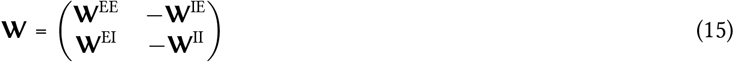

where the elements of **W**^IE^,**W**^EI^, and **W**^II^ were set to 2.4*/N*, 8*/N*, and 2.6*/N*, respectively. The excitatory sub-matrix **W**^EE^ was constructed by dividing the population of excitatory cells into six clusters (of 9 neurons each), with each cluster corresponding to one of the stimulus cue conditions. Connections within each cluster were strong, with a value of 30*/N*. Connections between neurons belonging to clusters that corresponded to adjacent stimulus cues were weaker, with a value of 2.5*/N*. All other connections were very weak, with a value of 0.02*/N*. This resulted in a block circulant structure for **W**^EE^.

Stimulus cue inputs were set to

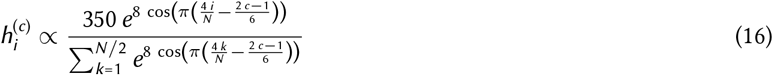

for cues *c* = 1, …, 6 and cells *i* = 1, …, *N /*2 (i.e. inputs were only delivered to the excitatory neurons).

#### 1.5.3 Linear integrator model

The linear integrator model that we used (Extended Data Fig. 1c and Extended Data Fig. 8a,d) has been described previously (see Ref. 6). There were no separate excitatory and inhibitory populations in this model, and the weight matrix was constructed such that network dynamics were non-normal, non-oscillatory, and stable with a single two-dimensional neutrally stable subspace (i.e. a plane attractor). We achieved this by defining **W** via its eigen-decomposition:

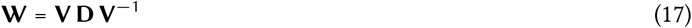

where the eigenvectors (columns of **V**, denoted as **v**_*j*_, for *j* = 1, …, *N*, with elements *v*_*ij*_, for *i, j* = 1, …, *N*) were generated by the following process:

1. Generating a random vector:

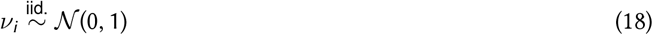

for *i* = 1, …, *N*.
2. Making the first 10% of vectors overlapping so that the resulting matrix is non-normal:

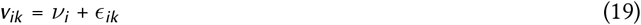

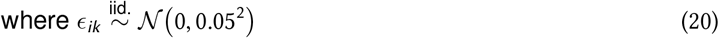

for *i* = 1, …, *N* and *k* = 1, …, *K* with *K* = 0.1 *N*.
3. Making the the other 90% of vectors orthogonal:

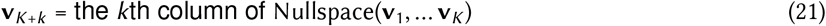

for *k* = 1, …, *N − K*
4. Unit normalizing each vector:

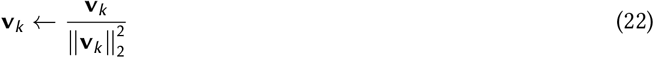

and the eigenvalues (*λ*_*i*_, for *i* = 1, …, *N*, the diagonal elements of the diagonal matrix **D**) were generated by the following process

1. Generating random (real) values:

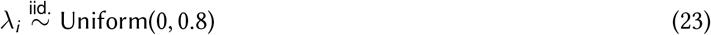

for *i* = 1, …, *N −* 2.
2. Creating a pair of neutrally stable eigenmodes:

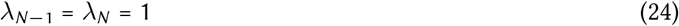

The stimulus cue inputs were set to

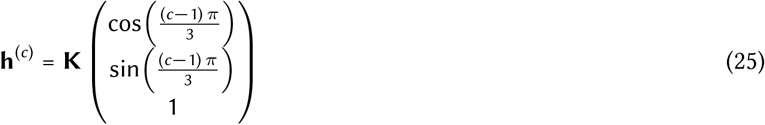

for cues *c* = 1, …, 6, and we considered two forms for **K**: either 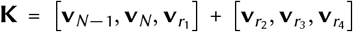 (Extended Data Fig. 1c and Extended Data Fig. 8a,d; as in the original formulation^6^) or 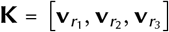 (Extended Data Fig. 8b,e), where *r*_1_, *r*_2_, *r*_3_, *r*_4_ were randomly drawn integers over the range 1 to *N* − 2. The first formulation of **K** ensured that stimulus cue inputs partially align with the persistent subspace, whereas the second formulation of **K** ensured that stimulus cue inputs align only with random directions.

#### 1.5.4 Feedforward network model

The linear feedforward network model that we used (Extended Data Fig. 1d and Extended Data Fig. 8c,f) has been described previously (see Refs. 21,27). (For pedagogical purposes, we used the simplest set up consisting of a feedforward chain of neurons, see below. However, using a more general network model that contained ‘hidden’ feedforward chains^21^ did not affect our analyses except for Extended Data Fig. 10e which, in contrast to the simple feedforward chain, could display overlap values greater than 0.5.) There were no separate excitatory and inhibitory populations in this model, and the weight matrix included a single chain running from neuron 1 to neuron *N* :

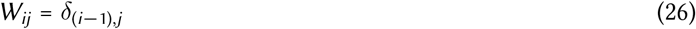

for cell-pairs *i, j* = 1, …, *N*.

The stimulus cues provided random inputs delivered to only the first 10 neurons so that each input could pass through the feedforward network:

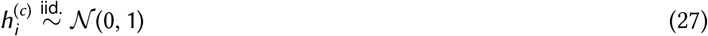

for cues *c* = 1, …, 6 and cells *i* = 1, …, 10.

### 1.6 Canonical nonlinear systems with two stable fixed points

In order to illustrate the applicability of our analysis of optimal information loading in linear dynamical systems to the behaviour of nonlinear dynamical systems, we first studied two variants (either symmetric or non-symmetric) of a canonical nonlinear system that can exhibit two stable fixed points. (These systems are closely related to the damped, unforced Duffing oscillator which is a classic example of a [non-symmetric] system that can exhibit two stable fixed points. Additionally, the analysis of these systems also holds for the Duffing oscillator.)

The dynamics of the first system (which has a symmetric Jacobian matrix) are governed by

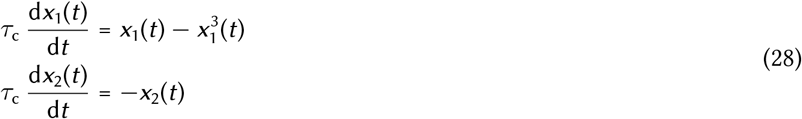

and the dynamics of the second system (which has a non-symmetric Jacobian matrix) are governed by:

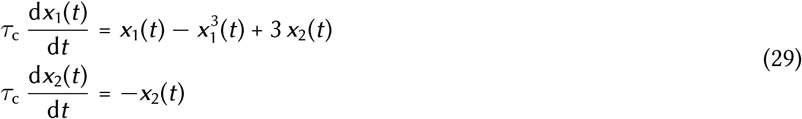

We used a cubic polynomial in Eqs. 28 and 29 because it is the lowest order polynomial that allows a system to exhibit 2 stable fixed points. Both systems exhibit 3 fixed points: both have a saddle point at the origin and both have 2 asymptotically stable fixed points at (± 1, 0) (see Extended Data Fig. 6 for the state space dynamics of these two systems).

We solved the dynamics of Eqs. 28 and 29 using a first-order Euler approximation starting from *t* = 0 with a discretization time step of 0.02 and a time constant of *τ*_c_ = 0.4 (note time was unitless for this model).

### 1.7 Analysis methods

Here we describe methods that we used to analyse neural data. Whenever applicable, the same processing and analysis steps were applied to both experimentally recorded and model simulated data. As a first step in all our analyses, in line with previous work analysing neural population dynamics^86^, we removed the stimulus cue-independent time-varying mean activity from each neuron’s firing rate time series (see Fig. 5a for an example). (This was done separately for training and test data for cross-validated analyses, see below.) In most of our analyses, neural activities were aligned to stimulus cue onset defined to be at *t* = 0. However, due to the variable delay duration of the task (Fig. 1a), experimentally recorded neural activities were also aligned to go cue onset for analyses that required incorporating the late delay and go epochs (i.e. beyond the first 1.65 s after the stimulus cue onset; Fig. 5b–c, Extended Data Fig. 10a–c,g). For simulated neural activities, this was not necessary, as we always simulated our networks in a fixed-delay task for ease of analysis, even if they were optimized for a variable-delay task in accordance with how our experimental monkey subjects were trained.

#### 1.7.1 Identifying amplifying, persistent, and other subspaces in network dynamics

In order to understand the dynamics of neural networks with potentially complex and high-dimensional dynamics, and the way these dynamics depend on initial conditions, we identified specific subspaces within the full state space of these networks that were of particular relevance for our analyses. These subspaces served dual roles. First, as ‘intervention tools’, to ascertain their causal roles in high dimensional network dynamics, we used them to constrain the initial conditions of the dynamics of our networks (see also Methods 1.7.2). Second, as ‘measurement tools’, to reveal key aspects of the high-dimensional dynamics of neural networks, we used them to project high-dimensional neural trajectories into these lower dimensional subspaces (see also Methods 1.7.3).

Our main analyses relied on identifying the most persistent and most amplifying modes of a network. This required dynamics that were linear—either by construction, or by (locally) linearizing or linearly fitting dynamics that were originally nonlinear (see Table 1). We computed the most persistent mode(s) in one of two different ways. First, for networks that were either guaranteed to have stable dynamics by construction (i.e. those constructed *de novo*; Figs. 3 and 4 and Extended Data Figs. 4, 5 and 8), or were confirmed to be always stable in practice (i.e. those constructed by local linearization; Extended Data Fig. 3e, Extended Data Fig. 6, and Extended Data Fig. 7), we simply used the eigenvector(s) of the weight matrix **W** associated with the eigenvalue(s) that had the largest real part(s). Second, for networks that were fitted to nonlinear dynamics or recorded data, and whose dynamics could thus not be guaranteed to be stable (Fig. 5f, Fig. 6e, Extended Data Fig. 7d, and Extended Data Fig. 10f–h), we used the eigenvectors of **W** associated with the largest real eigenvalues that were less than or equal to 1 + *δ* (with *δ* = 0.05) (i.e. we find the slowest, or most persistent, modes of the network—the *δ* was mostly relevant only for the after-go-time networks of Fig. 6 and Extended Data Fig. 12 which exhibited eigenvalues substantially greater than 1 and setting *δ* less than 0.05 did not substantially change our results). (Note that an eigenvalue of 1 for **W** corresponds to an eigenvalue of 0 for the associated Jacobian of the dynamics due to the leak term.)

For computing the most amplifying modes, we performed an eigen-decomposition of the associated Observability Gramian **Q**^53,62^. Specifically, we obtained **Q** by solving the following Lyapunov equation:

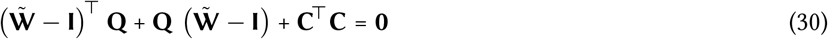

where 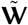 is the ‘stabilized’ weight matrix of the dynamics (and the *−***I** terms represent the effect of the leak on the Jacobian of the dynamics, Eq. 1) and **C** is the read-out matrix of the network. The most amplifying mode(s) of the network are given as the eigenvector(s) of **Q** associated with the largest eigenvalue(s). Again, for networks that were guaranteed to have stable dynamics by construction (Figs. 3 and 4, Extended Data Fig. 7a–c, Extended Data Fig. 5, and Extended Data Fig. 8), 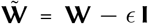, where **W** is the original weight matrix of the dynamics and *ϵ* = 0.01 (to ensure dynamical stability). For other networks, i.e. either linear networks fitted to experimental data (Fig. 5f and Extended Data Fig. 10f–h), linear networks fitted to simulated nonlinear dynamics (Fig. 6e and Extended Data Fig. 7d), or local linearizations of nonlinear dynamics (Extended Data Fig. 6, Extended Data Fig. 3e, and Extended Data Fig. 7a–c), we used 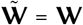 unless the largest eigenvalue *λ*_max_ of **W** was greater than or equal 1, in which case we used 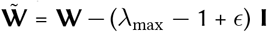, to ensure that the linear dynamics with 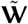 were stable (which is required for calculating **Q**). For networks obtained by fitting neural responses (experimentally recorded or simulated; Fig. 5f, Fig. 6e, Extended Data Fig. 7d, and Extended Data Fig. 10f–h), **C** was obtained by fitting those responses (Methods 1.4.3), as we wanted to understand how the fitted dynamics taking place in a latent space can generate the most discriminable fluctuations in (the principal components of) the neural responses to which they are related by this read-out matrix (although using **C** = **I** did not change our results substantially). For all other networks (Figs. 3 and 4, Extended Data Fig. 7a–c, Extended Data Fig. 3e, Extended Data Fig. 5, and Extended Data Fig. 8), we simply used **C** = **I**, as the activity of these networks was supposed to be read out in the same space within which their dynamics took place.

We also applied methods which did not rely on the linearization (or linear fitting) of network dynamics. Our goal was to develop basic intuitions for how much the dynamics of the different simulated nonlinear networks of Fig. 2 and Extended Data Fig. 2 used the persistent subspace of their dynamics. For this, we determined the ‘persistent subspace’ as the subspace spanned by the 5 principal components of the final 500 ms of neural activities (**x**) across all 6 cue conditions, corresponding to 6 distinct attractors, and the ‘persistent nullspace’ of the network as the 45-dimensional subspace orthogonal to the persistent subspace. For plots showing the projection of network activities within the persistent subspace (Extended Data Fig. 2b,f and Extended Data Fig. 2c–d and g–h, bottom) we used the first two principal components of the full, five-dimensional persistent subspace of the network, as determined above. For plots showing the projection of network activities to persistent vs. cue-aligned directions (Fig. 2d,j, and Extended Data Fig. 2c–d and g–h, top right), ‘persistent PC1’ was determined as the direction spanning the two persistent states corresponding to the two cue conditions being illustrated (i.e. as above, spanning the final 500 ms of neural activities across the two cue conditions), and ‘initial PC1 (orthogonalized)’ was determined as the the direction spanning the two initial conditions corresponding to the two cue conditions being illustrated, orthogonalized with respect to the corresponding persistent PC1.

#### 1.7.2 Subspace-constrained initial conditions

When using the subspaces identified above as ‘intervention tools’, to constrain the initial conditions of our networks, we either used the single top most persistent or amplifying mode for linear networks with low-dimensional coding spaces (including the linearized canonical nonlinear attractor dynamical system; Figs. 3 and 4 and Extended Data Figs. 5 and 6), or numerically optimized initial conditions within the corresponding higher-dimensional subspaces (Fig. 2f,l, Extended Data Fig. 2c,d,g,h, Extended Data Fig. 7; see also Methods 1.3 and Methods 1.4). When the persistent subspace was extracted from neural responses (rather than from the dynamical equations of the network, Methods 1.7.1; Fig. 2f,l, Extended Data Fig. 2c,d,g,h, Extended Data Fig. 7a) we used different sets of simulations to generate data from which we could estimate the persistent subspace (as explained above), and to analyse network dynamics when initialized within these subspaces. In all cases, for a fair comparison, the magnitude of initial conditions was fixed (Methods 1.3.1, Methods 1.4.1), and only their direction was affected by constraining them to one of these subspaces.

#### 1.7.3 Measures of subspace overlap

In order to measure the overlap of high dimensional neural dynamics with the subspaces we identified, we used one of two methods. First, for analysing network dynamics across two conditions chosen to correspond to ‘opposite’ stimulus cues (Fig. 2d,j, Fig. 3c,d,e, Extended Data Fig. 2c,d,g,h, Extended Data Fig. 6c,d, and Extended Data Fig. 5), such that the coding part of the persistent subspace was one-dimensional, we simply measured the projection of neural dynamics onto the first eigenvector (i.e. the eigenvector associated with the largest real eigenvalue) of the corresponding subspace using a dot product:

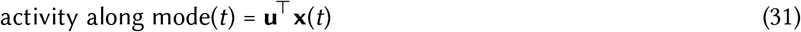

where **u** may correspond to the most persistent, or the most amplifying mode, or the first PC of the persistent-orthogonalized cue subspace (as defined above). We also used the same measure for visualising the quality of fit of linear neural network dynamics to experimental data (Methods 1.4.3) with **u** being the first PC of the full state space of neural firings rates (Fig. 5e). In those cases, when **u** had to be estimated from neural responses (Fig. 2d,j, Fig. 5e, Extended Data Fig. 2c,d,g,h), we used a cross-validated approach, using different subsets of the data to determine **u** and **x**(*t*) (from a single split of the data). In other cases, **u** was determined from the truly deterministic dynamics of the system and thus there was no need for cross-validation.

Second, to measure subspace overlaps for *d*-dimensional neural activities across multiple conditions and time points within coarser time bins (Fig. 4c, Fig. 5f, Fig. 6e, Extended Data Fig. 7d, Extended Data Fig. 8d–e, Extended Data Fig. 11c, Extended Data Fig. 12d, Extended Data Fig. 13d, and Extended Data Fig. 10b,c,f,g), thus corresponding to high-dimensional coding sub-spaces, we used the following properly normalized measure:

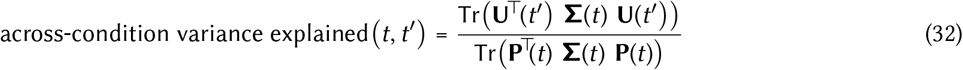

where **Σ**(*t*) is the covariance matrix of neural activities across conditions and raw (1-ms) time points within time bin *t*, the columns of **P**(*t*) are the first principal components of neural activities within time bin *t* (i.e. the eigenvectors of **Σ**(*t*) associated with the largest eigenvalues), and **U**(*t′*) is the subspace of interest with respect to which overlaps are computed (which itself may or may not depend on time, see below). The time resolution of *t* and *t′* (i.e. the duration of time bins within which data was used to compute the corresponding terms at a given *t* or *t*^*′*^), the choice of **U**(*t′*), and the number of vectors used for constructing **U**(*t′*) and **P**(*t*) depended on the analysis (see below).

Specifically, for measuring subspace overlap between neural activity and persistent vs. amplifying modes (Fig. 4c, Fig. 5f, Fig. 6e, Extended Data Fig. 7d, Extended Data Fig. 8d–e, and Extended Data Fig. 10f,g), we set **U**(*t′)* = **U** where the columns of **U** are the first *d/*4 eigenvectors of the most persistent or amplifying subspace (orthogonalized using a QR decomposition for the most persistent modes—this was not necessary for most amplifying modes which are orthogonal by construction), or *d/*4 randomly chosen orthonormal vectors as a control (shown as ‘chance’; computed analytically as 1*/*4 for ‘de novo’ linear networks (Fig. 4c and Extended Data Fig. 8d–e), and numerically for fitted linear networks, also yielding values of approximately 1*/*4, Fig. 5f, Fig. 6e, Extended Data Fig. 7d, and Extended Data Fig. 10f). **P**(*t*) contained the first *d/*4 principal components. In this case, a value of 1 for this metric implies that the *d/*4 directions of greatest variability in neural activity overlap exactly with the *d/*4-dimensional subspace spanned by **U**. The time resolution of *t* was 20 ms (for clarity, bins to be plotted were subsampled in the corresponding figures). Note that when this analysis was performed on linear networks fitted to neural data (experimentally recorded or simulated), **U, P**(*t*), and **Σ**(*t*) were all obtained from the same fitted linear network (i.e. no cross-validation). Specifically the parameters of the network were used to determine **U** (see Methods 1.4.3), and the neural responses these fitted linear dynamics generated (rather than the original neural responses that were fit by the linear model) were used to determine **Σ**(*t*) and thus **P**(*t*). See Methods 1.8 for computing the significance of these overlaps (and their differences). When analysing optimized ring attractor networks (Extended Data Fig. 3e), we used 2-dimensional subspaces (rather than *d/*4-dimensional subspaces) because we found empirically that the obtained ring attractors lay in a 2-dimensional subspace.

For analyzing subspace sharing between different task epochs (Extended Data Fig. 11c, Extended Data Fig. 12d, Extended Data Fig. 13d, and Extended Data Fig. 10b), **U**(*t*′) contained the top *k* principal components (PCs) of neural activity within the time bin indexed by *t*′ (we used *k* = 10 for the monkey data and *k* = 4 for our models because the models typically exhibited lower dimensional dynamics), while **P**(*t*) included all PCs within the time bin indexed by *t*. For these, we performed principal components analysis with dimensions corresponding to neurons and data points corresponding to time points and cue conditions. The time resolution of both *t* and *t*′ was 250 ms, such that the time periods (relative to cue onset) that we used were *−*500 to *−*250 ms (spontaneous epoch), 0 to 250 (cue epoch), 1250 to 1500 ms (delay epoch), and the first 250 ms after the go cue, i.e. *t*_go_ to *t*_go_ + 250 ms (go epoch). In this case, **U**(*t*′), **P**(*t*) and **Σ**(*t*) were obtained by fitting all the available neural data (i.e. no cross-validation). See also Ref. 64 for an ‘alignment index’ metric that is closely analogous to this use of this metric.

For showing how much variance the top 2 delay epoch PCs capture over time (Extended Data Fig. 10c), in line with Ref. 6, we set **U**(*t*′) = **U** where the columns of **U** are the first 2 principal components of neural activities over the time period 750 to 250 ms before the go cue, i.e. *t*_go_ *−* 0.75 to *t*_go_ *−* 0.25 s, and **P**(*t*) also includes the top 2 principal components. The resolution for *t* was 10 ms (for clarity, bins to be plotted were subsampled in the corresponding figure). In this case, we estimated **U** and **P**(*t*) in a cross-validated way (as in Ref. 6)—we estimated **U** using training data and **P**(*t*) and **Σ**(*t*) using test data, and we show results averaged over 10 random 1:1 train:test splits of the data. See also Ref. 6 for a measure that is closely related to this use of this metric, but uses the number of neurons in the denominator instead of the total variance.

#### 1.7.4 Linear decoding

We fitted decoders using linear discriminant analysis to decode the stimulus cue identity from neural firing rates (Fig. 2e,f,k,l, Fig. 4a,b, Fig. 5b,c, Fig. 6c,d, Extended Data Fig. 7c, Extended Data Fig. 3d, Extended Data Fig. 8a–c,Extended Data Fig. 10a,h, Extended Data Fig. 11a,b, Extended Data Fig. 12b,c, and Extended Data Fig. 13b,c). We constrained the decoders to be 2-dimensional (in line with previous studies^6^) because this was a sufficient dimensionality to decode responses. (We also trained decoders using logistic regression in the full activity space and obtained qualitatively similar results; not shown.) We primarily considered two types of decoding analyses: we either trained decoders on late delay activity and tested on all time points (‘delay-trained decoder’, e.g. Fig. 4a), or we trained decoders separately at every time point and tested on all times (‘full cross-temporal decoding’, e.g. Fig. 4b). In all cases, we measured decoding performance in a cross-validated way, using separate sets of neural trajectories to train and test the decoder, and we show results averaged over 10 random 1:1 train:test splits of the data. For delay-trained decoders, training data consisted of pooling neural activity over a 500 ms time interval (the time interval is shown by a horizontal black bar in all relevant figures), and tested the thus-trained decoder with data in each 1 ms time bins across the trial (for clarity, test bins to be plotted were subsampled every 10 ms in the corresponding figures). For full cross-temporal decoding, we binned neural responses into 10 ms time bins and trained and tested on all pairs of time bins (specifically, we plotted mean decoding performance across the 10 1-ms raw time bins corresponding to each 10-ms testing bin). We used a shrinkage (inverse regularisation parameter on the Euclidean norm of decoding coefficients) of 0.5 (we also tested various other values and found qualitatively similar results; not shown). Chance level decoding was defined as 1*/C*, where *C* = 2 or 6 is the number of cue conditions that need to be decoded (Tables 1 and 3).

#### 1.7.5 Quality of fit for linear models fitted to neural responses

When fitting linear models to neural data (experimentally recorded or simulated; Methods 1.4.3) we used a crossvalidated approach for measuring the quality of our fits, with a random 1:1 train:test split of the data (Fig. 5d). For this, we first fitted the model on training data (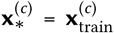 in Eq. 9). The quality of fit was then computed on the test data, 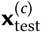, as the fraction of variance of 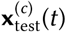 explained by the simulated responses (after the appropriate projection, i.e. **Cx**^(*c*)^(*t*)), across all 20 dimensions weighted by **D** (all parameters, including **P, C** and **D**, were set to their values obtained by fitting the training data). In other words, we computed the Pearson *ℛ*^2^ with respect to the identity line using the mean squared error, *ε*^2^ in Eq. 8, with the momentary error in Eq. 9 computed using 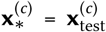. Once the quality of fit for this split was thus established, we conducted all further analysis involving fitted linear models with the model that was fit to the training half of this split.

As a meaningful lower bound on our quality of fit measure, we also computed the same measure (i.e. fitting a linear neural networks to training data and calculating the quality of fit using test data) for 100 different time-shuffled controls of the original train:test split of the data (Methods 1.4.3), such that we shuffled time bins coherently between the training and the test data, across neurons and conditions (Fig. 5d, dark gray).

To calibrate how much our fits were limited by the noisiness of the data, we also computed the quality of fit directly between 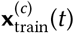 and 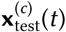 (i.e. using the mean squared error, *ε*^2^ in Eq. 8, with the momentary error redefined as 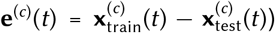 for 100 random 1:1 train:test splits of the data (Fig. 5d, light gray). The extent to which the *ℛ*^2^ computed with this control was below 1 reflected the inherent (sampling) noise of the experimental data that limited the quality of fit obtainable with any parametric model, including ours that was based on linear dynamics. Moreover, a cross-validated *ℛ*^2^ computed with our fits that was higher than the *ℛ*^2^ obtained with this control (Fig. 5d dark and light blue vs. light gray) meant that the inherent assumption of linear dynamics in our model acted as a useful regularizer to prevent the overfitting that this overly flexible control inevitably suffered from. See more in Methods 1.8 on statistical testing for our quality of fit measure.

When fitting to simulated neural data, we obtained high quality of fits using the same measure (*ℛ*^2^ *>* 0.95, not shown).

#### 1.7.6 Overlap between the coding populations during the cue and delay epochs

To test whether separate neural populations encode stimulus information during the cue and delay epochs (Extended Data Fig. 10e), we trained (non-cross validated) decoders to decode cue identity using logistic regression on either cue-epoch activity (‘cue-trained’; the first 250 ms of activity after cue onset) or delay-epoch activity (‘delay-trained’; 1250–1500 ms after cue onset). We used an L2 regularisation penalty of 0.5 (we also tested other regularisation strengths and observed no substantial changes in our results). We took the absolute value of decoder weights as a measure of how strongly neurons contributed to decodability (either positively or negatively). We then binarized the absolute ‘cue-trained’ and ‘delay-trained’ weights using their respective median values as the binarization threshold. (This binarization reduces a potential bias effect from large or small weight values in our analysis.) Our measure of overlap between the coding populations during the cue and delay epochs, was then simply the inverse normalized Hamming distance between these two sets of binarized weights:

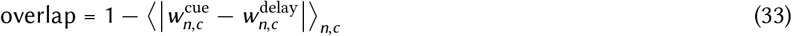

where 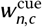 (‘cue trained’) and 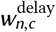 (‘delay trained’) is the binarized weight of neuron *n* in cue condition *c* during the cue and delay epochs, respectively, and ⟨·⟩_*n,c*_ denotes taking the mean across neurons and cue conditions. For completely overlapping populations, this measure takes a values of 1, for completely non-overlapping populations, it takes a values of 0, and for random overlap (shown as ‘chance’) it takes a values of 0.5.

For the shuffle controls, we randomly permuted the neuron indices of the delay-trained weights (such that using the median as a threshold thus resulted in values close to 0.5, i.e. chance level; Extended Data Fig. 10e). We show results (for both the original analysis and shuffle control) for 10 random halves of the data (equivalent to the training halves of 10 different 1:1 train:test splits). We also tested a variety of percentile values other than the median and our results did not change substantially (choosing a threshold other than the median causes both the data and shuffle controls to have overlap values lower than those that we obtained with the median as the threshold, but it does not substantially affect the difference between them). As an additional control, we also removed neurons that did not contribute to decodability: we removed neurons that had a thresholded weight of 0 for all 6 cue conditions in both the cue and delay epochs. This resulted in removing 13.3 neurons on average for monkey K and 33.5 neurons for Monkey T (when using the median as the threshold) and our results did not change substantially (not shown).

#### 1.7.7 Finding fixed / slow points

For finding the fixed / slow points of nonlinear network dynamics (Fig. 2d,j, Extended Data Fig. 2b,f, Extended Data Fig. 3a, and Extended Data Fig. 11d), we used a slow-point analysis method^17^ that searches for an **x** for which the L2 norm of the gradient determined by the autonomous dynamics of the network is below a threshold. Note that this was only possible in model neural networks as the method requires access to the equations (and parameters) defining the true (nonlinear) dynamics of a system.

Specifically, for network dynamics governed by (cf. Eqs. 1 and 2)

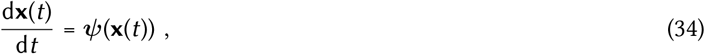

for some function ***ψ***, we sought to find points **x**^*\**^ such that ||***ψ***(**x**^*\**^) ||_2_ is small. To achieve this, we drew 1000 **x**’s from a spherical Gaussian distribution with mean 0 and variance 10 (the large variance helps to ensure that we cover a large part of state space) and we optimized each **x** to minimize ||***ψ***(**x**) ||_2_ using gradient descent with gradients obtained by back-propagation with an Adam optimizer^85^. We used an adaptive learning rate (which we found worked substantially better than a fixed learning rate in this scenario) that started at 0.1 and halved every 1000 training iterations (we used 5000 training iterations in total). Finally, we identified the **x**’s obtained at the end of optimization as asymptotically stable fixed points, **x**^*\**^, if ||***ψ***(**x**)||_2_ *<* 0.001 and if the largest real part in the eigenvalues of the linearization of ***ψ***(**x**) around **x**^*\**^ was less than 0.

#### 1.7.8 Correlations between initial and final neural firing rates

To measure correlations between initial and final simulated activities, we used the Pearson correlation coefficient (with respect to the identity line) between initial and final mean-centered firing rates across neurons within the same simulation (i.e. no cross-validation; Fig. 2b,h; insets). Histograms show the distribution of this correlation across 6 cue conditions (and the 10 different networks, each simulated 10 times, see above) using a kerneldensity estimate (Fig. 2c,i, Extended Data Fig. 2c,d,g,h, and Extended Data Fig. 3c).

### 1.8 Statistics

We performed statistical hypothesis testing in two cases.

First, we tested whether the quality of fit of linear models to experimental data was sufficiently high using permutation tests. To construct the distribution of our test statistic (cross-validated *ℛ*^2^, see also Methods 1.7.5) under the null hypothesis, we used *n* = 200 different random time shuffles of the data (Fig. 5d, dark gray), such that we shuffled time bins coherently between the training and the test data, across neurons and conditions, and for each shuffle used the same random 1:1 train:test split as for the original (unshuffled) data. For additional calibration, we also constructed the distribution of our test statistic under the alternative hypothesis that all cross-validated errors were due to sampling noise differences between the train and test data. For this, we used *n* = 200 random 1:1 train:test splits of the (original, unshuffled) data, and measured the quality of fit directly between the test data and the training data (rather than a model fitted to the training data, see also Methods 1.7.5; Fig. 5d, light gray). In both cases, we computed the two-tailed p-value of the test statistic as computed on the real data (Fig. 5d, blue lines) with respect to the corresponding reference distribution.

Second, we also used a permutation test-based approach to test whether the experimentally observed overlaps with persistent and amplifying modes (or their differences) were significantly different from those expected by chance. For testing the significance of overlaps in a given time step, we constructed the distribution of our test statistics (the overlap measures; Methods 1.7.3) under the null hypothesis by generating *n* = 200 random subspaces within the space spanned by the 20 PCs we extracted from the data (Methods 1.4.3), dimensionality matched to the persistent and amplifying subspaces (i.e. 5 orthogonal dimensions), and computed the same subspace overlap measures for the data in the given time step with respect to these random subspaces (Fig. 5f and Extended Data Fig. 10f–g; gray line and shading). For testing the significance of differences between overlaps (amplifying vs. persistent at a given time step, or amplifying or persistent between two different time steps), our test statistic was this difference (i.e. a paired test), and our null distribution was constructed by measuring it for *n* = 200 pairs of random subspace overlaps at the appropriate time step(s). Once again, in all these cases we computed the two-tailed p-value of the test statistic as computed on the real data (Fig. 5f and Extended Data Fig. 10f–g, green and red lines) with respect to the corresponding reference distribution.

Note that we did not compute p-values across multiple splits of the data because this led to p-value inflation as we increased the number of splits. Instead, we repeated all relevant analyses on 10 different random 1:1 train:test splits to see if our results were robust to the choice of data split. Indeed, we obtained qualitatively and quantitatively (in terms of p-values for quality of fits, and overlaps) similar results for all these splits.

Permutation tests do not assume that the data follows any pre-defined distribution. No statistical methods were used to predetermine experimental sample sizes. Sample sizes for permutation tests (*n* above) were chosen so as to be able to determine p-values to a precision of 0.01 (quality of fits) or 0.01 (subspace overlaps).

## Extended data figures

**Extended Data Fig. 1.**
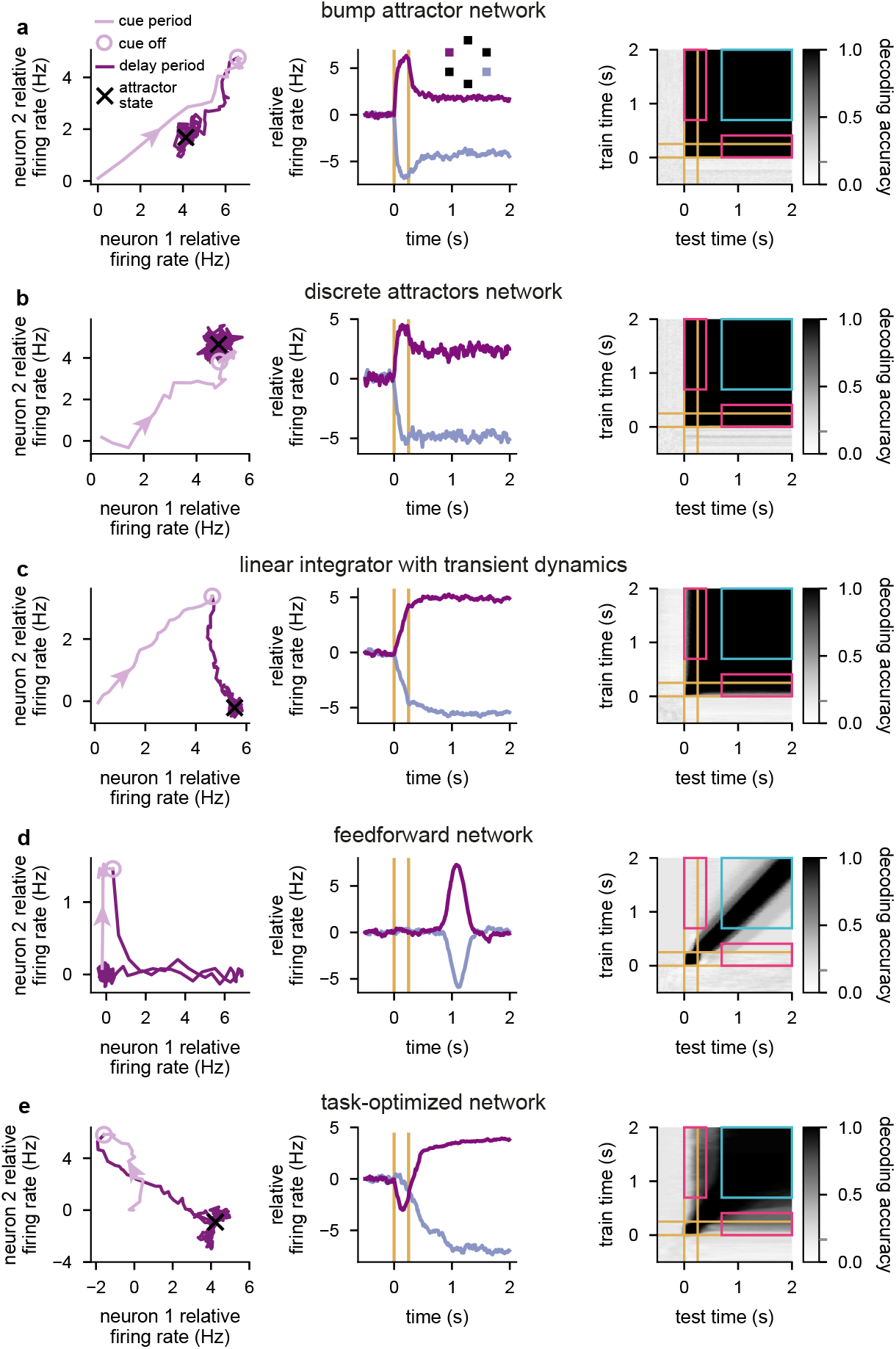
Dynamics of network models of working memory. **a**, Neural network dynamics in a bump attractor network^5^ performing the task shown in Fig. 1a. Left: trajectory in neural state space in a single cue condition during the cue period (pale purple line, ending in pale purple circle) and delay period (dark purple line). Purple arrow heads indicate direction of travel along the trajectory, black cross shows attractor state. Center: time course of relative (i.e. mean-centered) firing rates of one neuron for two cue conditions (purple vs. blue, see also inset). Yellow lines indicate cue onset and offset times. Right: cross-temporal decoding of neural activity produced by the network across all 6 cue conditions. Pink rectangles indicate generalized decoding between the cue/early delay period and the late delay period and cyan square indicates generalized decoding between time points in the late delay period. The gray tick on the color bar indicates chance-level decoding. **b**, Same as **a** but for a discrete attractors model^5,31,69^. **c**, Same as **a** but for a linear integrator model with transient dynamics that are orthogonal to the attractor subspace^6^. **d**, Same as **a** but for feedforward network model^21,27^. **e**, Same as **a** but for a network whose parameters (including recurrent, input, and readout weights) were optimized to perform the task shown in Fig. 1a (cf. Fig. 6d, right).

**Extended Data Fig. 2.**
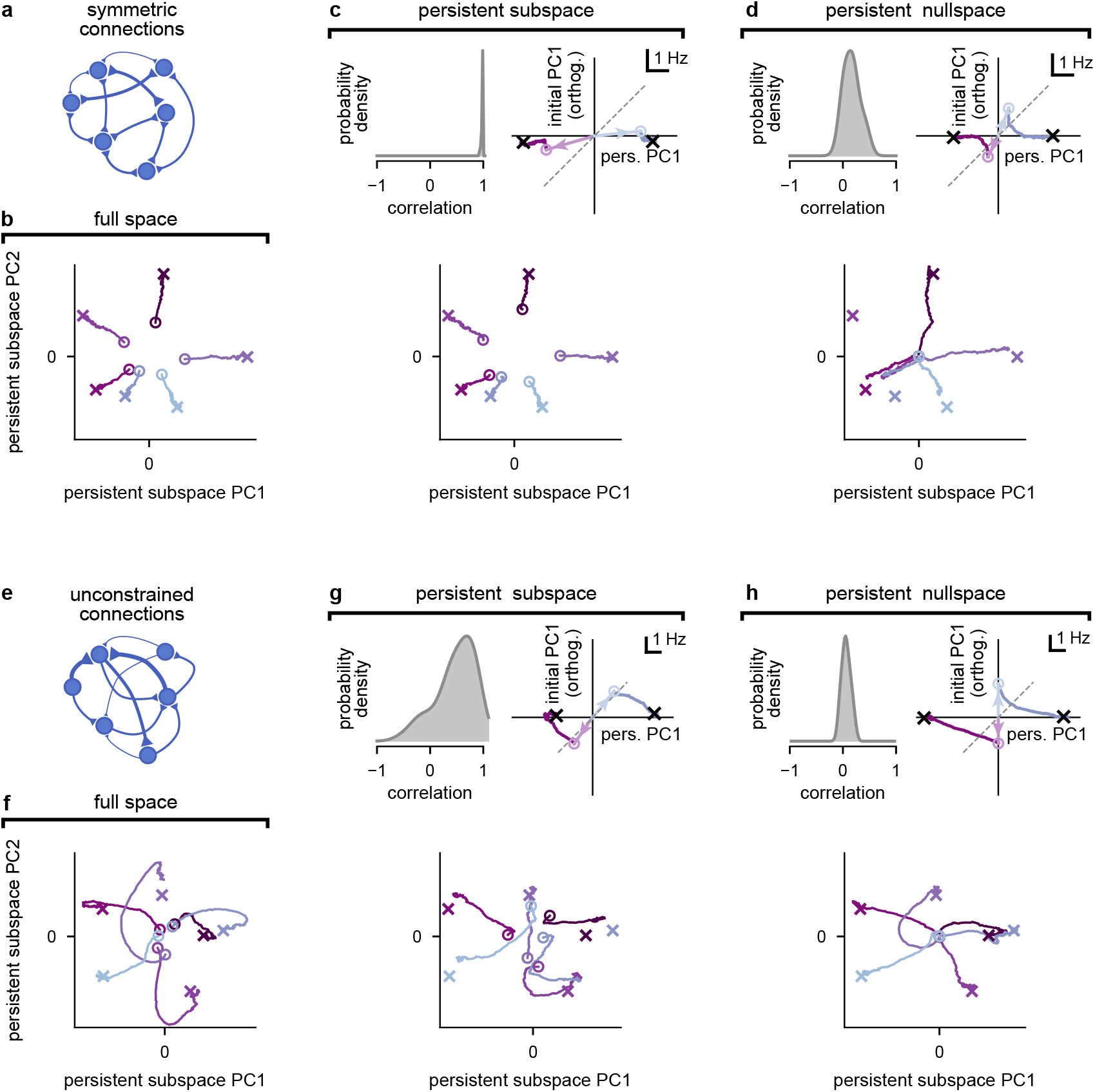
Attractor network dynamics with or without constraints on the initial condition of the dynamics. **a**, Illustration of an attractor network with symmetric connections. **b**–**d**, Analysis of neural responses in symmetric attractor networks (such as shown in **a**). **b**, Sub-threshold activity (colored trajectories) for all 6 cue conditions (color coded as in Fig. 5e) with initial conditions optimized within the full state space (Methods 1.3.1). Open circles show the optimized initial conditions and crosses show stable fixed points. We show neural activity projected onto the top two principal components of the persistent subspace. **c**, Analysis of neural responses when initial conditions are constrained to lie within the 5-dimensional persistent subspace. Top left: distribution of Pearson correlations between initial and final mean-centered neural firing rates across all 6 cue conditions and 10 networks (same as Fig. 2c, but for persistent subspace-constrained inputs, corresponding to green line in Fig. 2f). Top right: sub-threshold activity for 2 cue conditions in an example network (colored trajectories; same as Fig. 2d, but for persistent subspace-constrained inputs, corresponding to green line in Fig. 2f). Open circles (with arrows pointing to them from the origin) show the optimized initial conditions, black crosses show stable fixed points, dashed gray line is the identity line. Horizontal axis (persistent PC1) shows neural activity projected on to the 1st principal component (PC1) of network activities at the end of the delay period (across the 2 conditions shown), vertical axis (initial PC1 (orthogonalized)) shows projection to PC1 of initial neural activities orthogonalized to persistent PC1. Bottom: same as **b**, but for persistent subspace-constrained inputs, corresponding to green line in Fig. 2f. **d**, Same as **c**, but for persistent nullspace-constrained inputs. Note that the distribution of Pearson correlations of neural firing rates (top left) is distinct from a delta function at 0 because we constrained the initial conditions in the space of sub-threshold activities (rather than firing rates). In the bottom panel, which shows sub-threshold activity, we see that indeed all the colored circles overlap at the origin, indicating orthogonality of the initial conditions to the persistent subspace. **e**–**h**, Same as **a**–**d** but for attractor networks without a symmetric connection constraint (i.e. panels **f, g**, and **h**, respectively correspond to the networks shown by the black, green, and red lines in Fig. 2). Note initial conditions being near the origin in **f** mean that they are strongly orthogonal to the persistent subspace (as in **d**, but without constraining them explicitly to be in the persistent nullspace).

**Extended Data Fig. 3.**
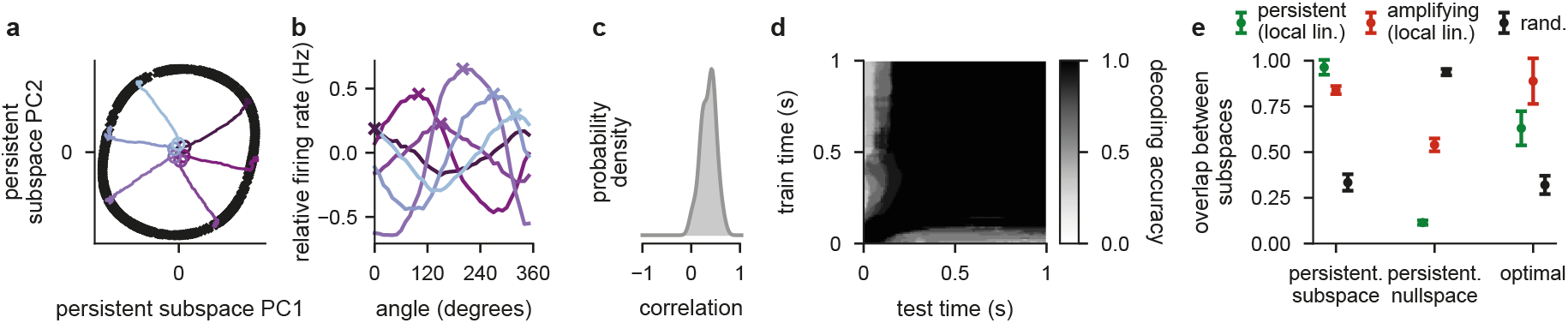
Dynamics of optimized ring attractor networks. **a**, Neural activity (colored trajectories) in a ring attractor network with unconstrained connectivity and optimized initial conditions (see Methods 1.3.1 and 1.3.4) for 6 cue conditions (color coded as in Fig. 5e). Open circles show the optimized initial conditions and black crosses show fixed points. We show neural activity projected onto the top two principal components of the persistent subspace. Thus, all circles being near the origin means that initial conditions are strongly orthogonal to this subspace (cf. Extended Data Fig. 2f). **b**, Tuning curves at *t* = 1 s for 6 example neurons (colored curves) whose preferred angles (colored crosses) correspond to the 6 cue conditions shown in **a. c**, Distribution of Pearson correlations between initial and final mean-centered neural firing rates across the 6 cue conditions and 10 networks (cf. Fig. 2i). **d**, Cross-temporal decoding of neural firing rate activity (cf. Fig. 2k). Note that only the first second of the delay period is shown on both axes because the dynamics of these networks, using a tanh nonlinearity, are faster than those shown in other figures (e.g. Fig. 2), using a ReLu nonlinearity (but the same time constant; Methods 1.2, and Table 1). **e**, Overlap (mean ± 1 s.d. across 10 networks) of the 2 locally most persistent (green), most amplifying (red), or random directions (black), obtained using a local linearization around the origin, with the ‘persistent subspace’ and ‘persistent nullspace’ of the original nonlinear dynamics, obtained without linearization, and the subspace spanned by the ‘optimal’ initial conditions of the original nonlinear dynamics (cf. Extended Data Fig. 7a, bottom; see Methods 1.4.2 and 1.7.3). We used 2-dimensional subspaces from the local linearization because we found empirically that the ring attractor lay in a 2-dimensional subspace (see also **a**).

**Extended Data Fig. 4.**
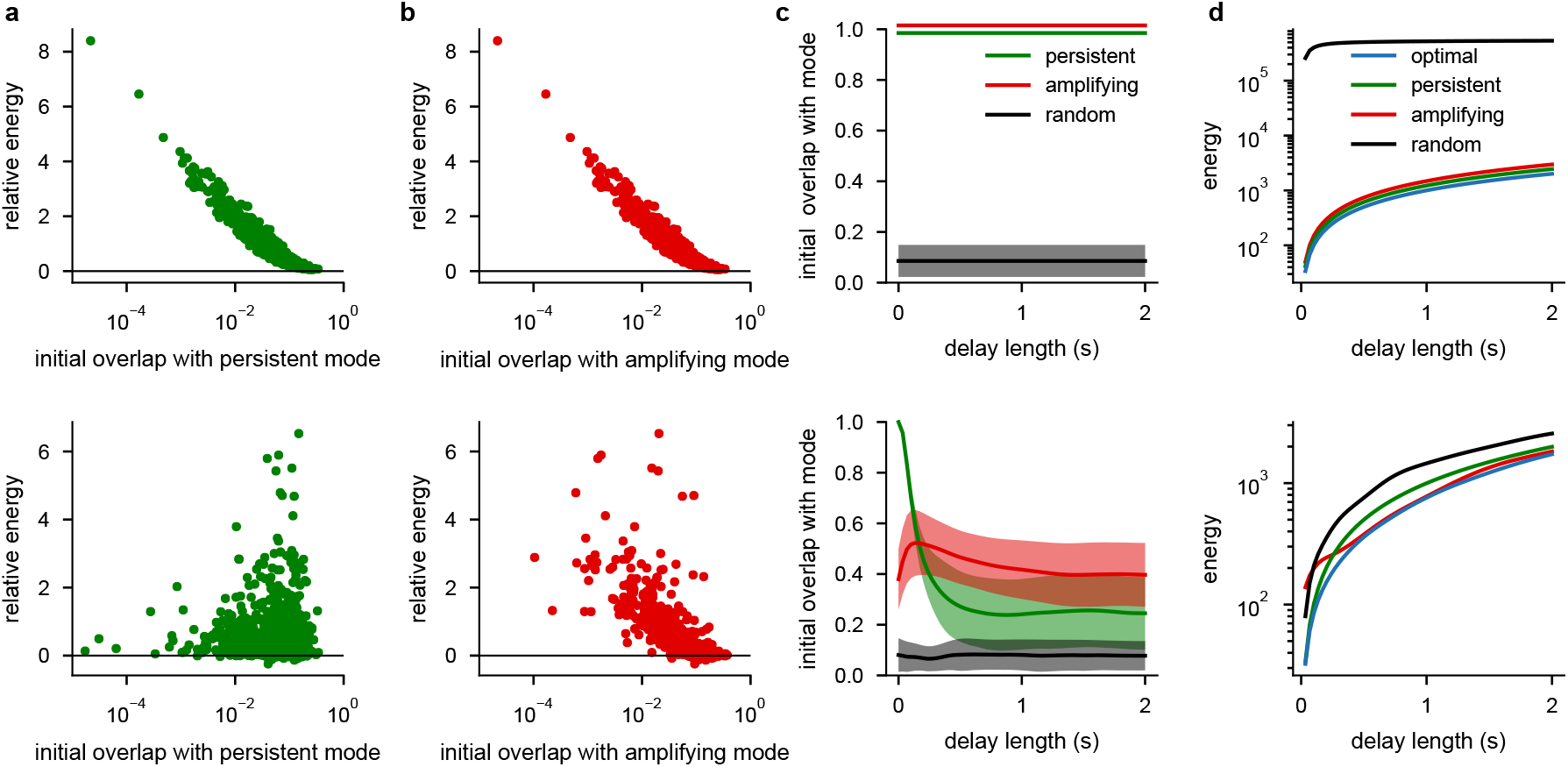
Analysis of the total energy produced by different initial conditions in linear networks. **a**, The norm of neural activity integrated over time (i.e. a measure of total energy used by the network) for each of 1000 random initial conditions (10 initial conditions for each of 100, 100-neuron networks) relative to the energy produced by the most amplifying initial condition, plotted as a function of their overlap with the persistent mode for symmetric (top) and unconstrained (bottom) linear integrator networks. A positive value on the y-axis means that the total energy produced by the given random initial condition is greater than that produced by the most amplifying initial condition. Initial conditions are scaled so that they all produce the same level of persistent activity (i.e. the same level of performance) after 2 s of simulation. **b**, Same as **a**, but initial conditions are plotted as a function of their overlap with the most amplifying mode. Note that overlap with the most amplifying mode (but not in general with the most persistent mode) is strongly predictive of total energy (with an inverse relationship between the two). **c**, Overlap (mean ± 1 s.d. across the 100 networks from **a** and **b**) of optimal initial conditions (Eq. S40), producing an overlap of 1 with the persistent mode after a given delay length (x-axis) while using the minimal total energy over time (Eq. S39), with either persistent (green), most amplifying (red), or random (black) directions, for symmetric (top) and unconstrained (bottom) networks. In unconstrained networks, for very short delay lengths, initial conditions must align exactly with the persistent mode, by necessity (green lines at 0 s). For longer delay lengths, initial conditions make greater use of the most amplifying direction (red lines). **d**, Total energy (mean across the 100 networks from **a** and **b**; we do not show error bars for visual clarity) for dynamics starting from initial conditions that produce an overlap of 1 with the persistent mode after a given delay length (x-axis) in symmetric (top) and unconstrained (bottom) networks. Initial conditions were chosen to be optimal (blue; i.e. using the least energy, cf. panel c), or aligned with the most persistent (green), most amplifying (red), or a random direction (black). In unconstrained networks, for very short delay lengths, initialising along the most persistent mode achieves near-optimal energy-efficiency (green is close to blue), but for longer delay lengths, initialising along the most amplifying mode becomes more energy efficient (red is closer to blue). (Note that for symmetric networks, top, we have offset the curves for the most amplifying, persistent, and optimal directions because these 3 directions are the same and therefore produce the same total energy.)

**Extended Data Fig. 5.**
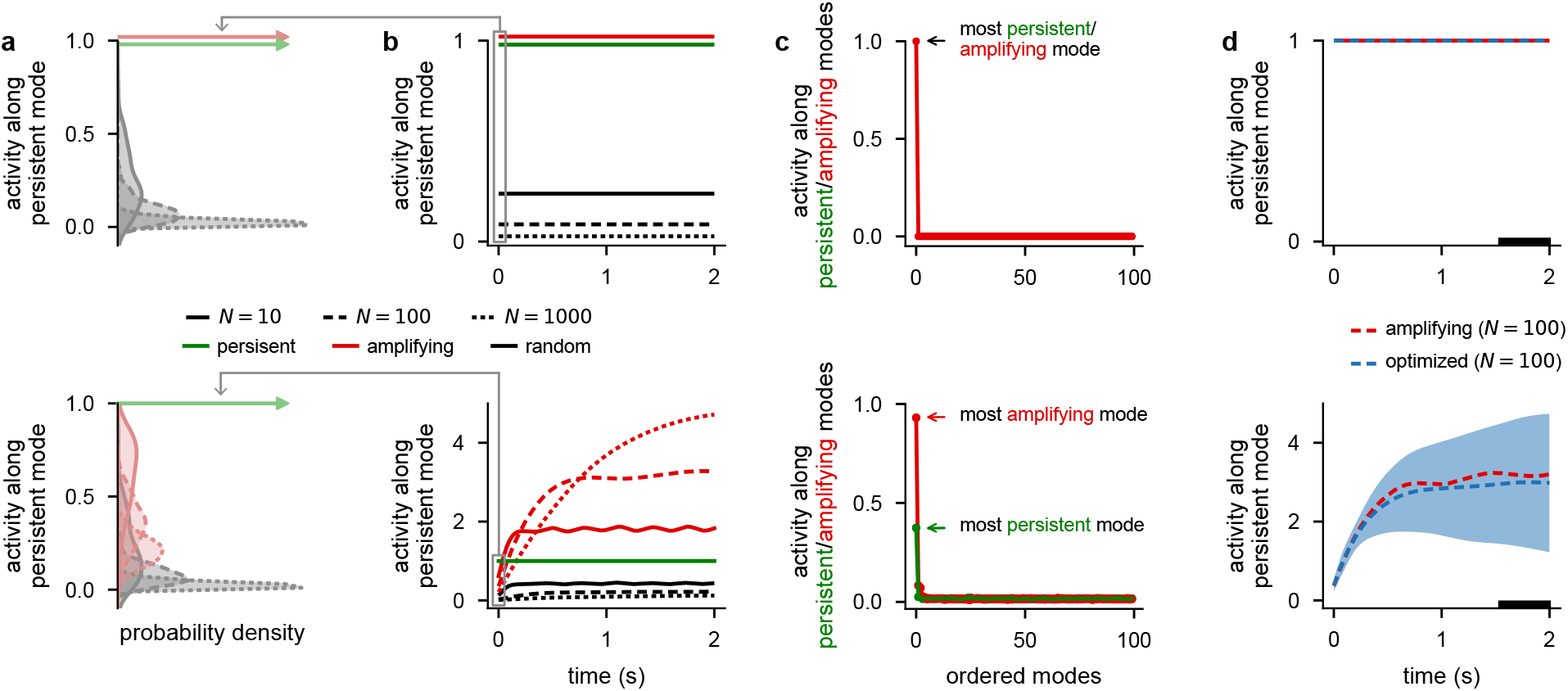
Analysis of linear networks of different sizes. **a**, Distributions of absolute overlap with the persistent mode for persistent (pale green), most amplifying (pale red), or random initial conditions (gray) across 100 randomly sampled linear symmetric (top) and unconstrained networks (bottom) consisting of either 10 (solid), 100 (dashed), or 1000 (dotted) neurons (cf. Fig. 3d). The persistent initial conditions produced delta functions at 1 (arrows). Results for persistent and most amplifying initial conditions are identical in symmetric networks (top). **b**, Time course of mean (across the 100 networks from **a**) absolute overlap with the persistent mode when starting network dynamics from persistent (green), most amplifying (red), or random initial conditions (black) in symmetric (top) and unconstrained networks (bottom) consisting of either 10 (solid), 100 (dashed), or 1000 (dotted) neurons (cf. Fig. 3e). Results for persistent and most amplifying initial conditions are identical in symmetric networks (top). **c**, Mean (across 100 networks) overlap of initial conditions that were optimized so as to generate maximal persistent activity in 100-neuron noisy symmetric (top) and unconstrained (bottom) networks with 100 orthogonal modes ordered by their persistence (green) or amplification (red) (i.e. corresponding to the rank ordered eigenvectors of the weight matrix, green, or of the observability Gramian of the dynamics, red; Methods 1.7.1). In symmetric networks (top), the optimized initial conditions overlap only with the most amplifying mode and no other mode (note that the most persistent mode is identical to the most amplifying mode in this case). In unconstrained networks (bottom), optimized initial conditions overlap strongly with the most amplifying mode and only weakly with other modes. (The non-zero overlap with the most persistent mode is simply due to the fact that there is a non-zero overlap between the most persistent and amplifying mode in random networks, and it is at the level that would be expected based on this overlap.) **d**, Time course of mean (across 100 networks) absolute overlap with the persistent mode for the same 100-neuron noisy symmetric (top) and unconstrained networks (bottom) as those shown in **c** when the network is started from optimized initial conditions (blue), and for comparison for the most amplifying (red dashed) initial conditions (cf. Fig. 3e). Note the close agreement between the two indicating that the most amplifying mode is indeed optimal in these networks. Horizontal black bar on x-axis shows the time period in which we applied the cost function to optimize the initial conditions (Methods 1.7.3).

**Extended Data Fig. 6.**
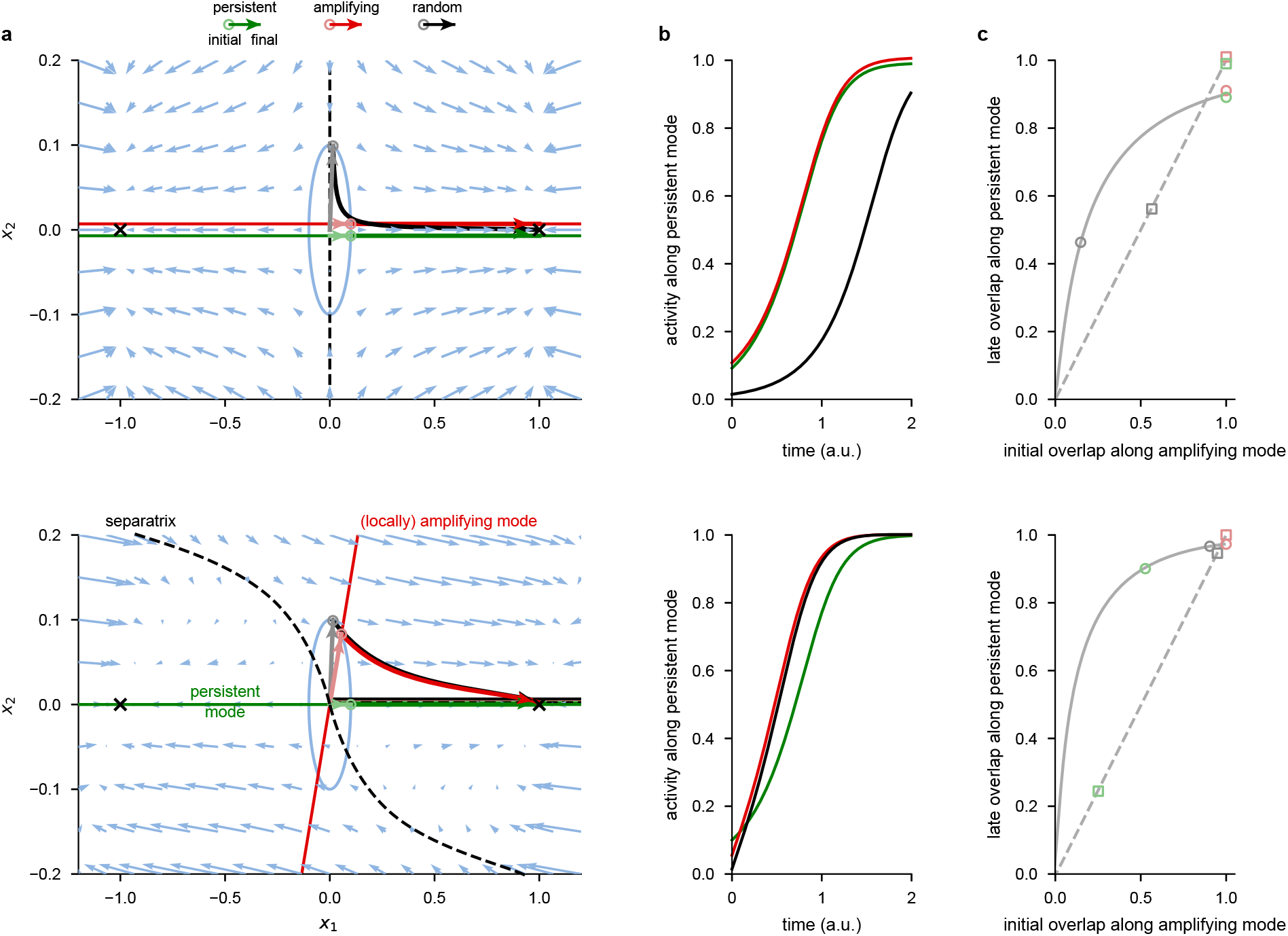
Analysis of canonical nonlinear attractor systems. **a**, State space of a canonical nonlinear system with two attractors and a symmetric (top) and non-symmetric Jacobian (bottom, see also Methods 1.6, Supplementary Information S3; cf. Fig. 3b). Pale blue arrows show flow field dynamics (direction and magnitude of movement in the state space as a function of the momentary state). Black crosses indicate asymptotically stable fixed points (i.e. attractor states), dashed black line shows the separatrix (the manifold separating the basins of attraction of the two attractors). Thin green and red lines indicate the locally most persistent and amplifying modes around the origin, respectively (lines are offset slightly in the top panel to aid visualisation). Pale green, red, and gray arrows with open circles at the end indicate most persistent, amplifying, and random initial conditions, respectively. Blue ellipses show the fixed initial condition norm around the origin to highlight the different axis scales. Dark green, red, and black arrows show neural dynamics starting from the corresponding initial condition. **b**, Time course of dynamics of the system along the persistent mode (i.e. the projection onto the green line in **a**) when started from the persistent (green), most amplifying (red), or random (black) initial conditions for the symmetric (top) and the unconstrained system (bottom). **c**, Late overlap with the locally persistent mode as a function of initial overlap with the locally most amplifying mode in the canonical nonlinear systems shown in panels **a**–**b** (solid gray line) and, for comparison, in the linear networks of Fig. 3a–c (dashed gray line) for symmetric (top) and unconstrained systems (bottom). Late overlap is measured as the mean overlap of activity along the persistent mode (panel **b**, from *t* = 0.8 to *t* = 2 for the canonical nonlinear system; Fig. 3c, from *t* = 0.8 s to *t* = 2 s for the linear networks). Open circles and squares indicate the random (gray), persistent (pale green), and most amplifying (pale red) initial conditions used respectively in panels **a** and **b** for the canonical nonlinear system, and in Fig. 3b–c for the linear networks.

**Extended Data Fig. 7.**
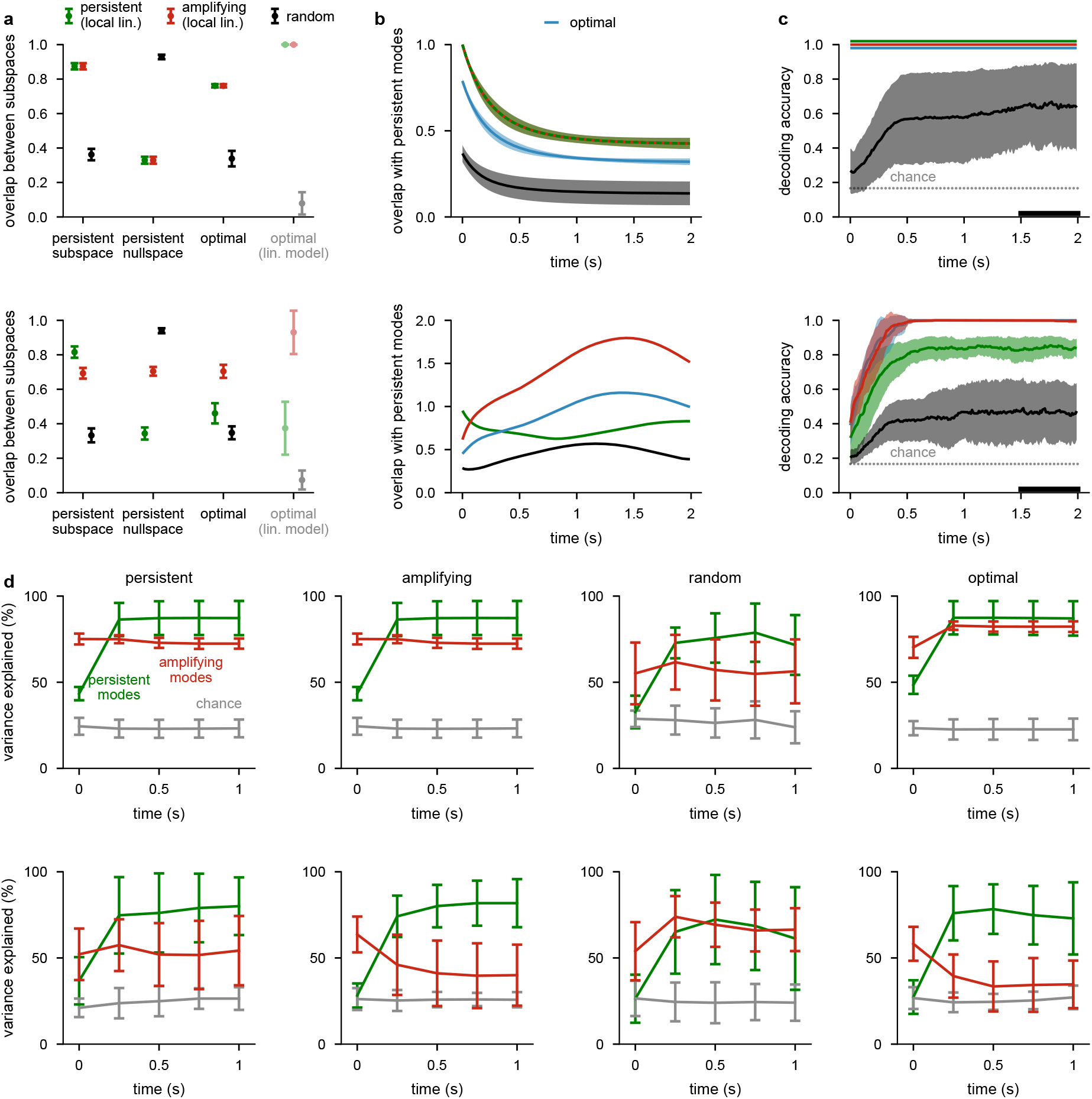
Linear analyses of the nonlinear attractor networks of Fig. 2. **a**, Overlap (mean ± 1 s.d. across 10 networks) of the 5 locally most persistent (green), most amplifying (red), or random directions (black) of the symmetric (top) and unconstrained (bottom) networks from Fig. 2, obtained using a local linearization around the origin, with the ‘persistent subspace’ and ‘persistent nullspace’ of the original nonlinear dynamics, obtained without linearization (as used in Fig. 2f and l, red and green), and the 5-dimensional subspace spanned by the 6 ‘optimal’ initial conditions of the original nonlinear dynamics (used in Fig. 2b–e, h–k, and f and l, black). For comparison, we also show the overlap (mean ± 1 s.d. across 100 networks) of the single most persistent (pale green), most amplifying (pale red), and random (gray) direction with the optimal initial condition of the linear networks from Extended Data Fig. 5c,d (‘optimal (lin. model)’). **b**, Time course of the overlap (mean ± 1 s.d. across 10 networks, s.d. not shown in bottom for visual clarity) of the linearized dynamics of symmetric (top) and unconstrained networks (bottom) with the subspace spanned by their most persistent modes when started from initial conditions that were optimized for the decoding accuracy of the nonlinear dynamics while constrained to be within the locally most persistent (green), most amplifying (red), or a random subspace (black). The linear dynamics, the persistent subspace wrt. which overlap is measured, and the subspaces within which initial conditions were constrained while being optimized, were all based on a local linearization of the nonlinear dynamics around the origin. Compare with Fig. 3e for the analogous plots for linear networks. For reference, blue line shows overlap of the same linearized dynamics when started from the initial conditions directly optimized for the decoding accuracy of the nonlinear dynamics without subspace constraints (used in Fig. 2b–e, h–k, and f and l, black). For consistency with Fig. 3b–e (where initial conditions were constrained to have unit norm), we scaled activity by the norm of the initial condition (which was constrained to be 3 here; Methods 1.4.2). **c**, Performance (mean±1 s.d. across 10 networks) of a delay-trained decoder (black bar indicates decoder training time period; Methods 1.7.4) on neural activity in stochastic nonlinear symmetric (top) and unconstrained networks (bottom) over time. Colors indicate initial conditions as in **b**. (Blue line shows same data as black line in Fig. 2f and l). Gray dotted line shows chance level decoding. Green, red, and blue lines are vertically offset slightly in the top panel to aid visualization. Compare with Fig. 4a (noise matched) for the analogous plots for linear networks (though with non-instantaneous inputs). **d**, Percent variance of responses explained (mean ± 1 s.d. across 10 networks) by the subspace spanned by either the 25% (i.e. 5) most persistent (green) or 25% (i.e. 5) most amplifying (red) modes as a function of time for 20-dimensional linear neural networks fitted to the neural responses generated by the symmetric (top) and unconstrained (bottom) nonlinear networks when started from the same (optimized) initial conditions analyzed in **b**–**c**: constrained to be within the locally most persistent (far left), most amplifying (center left), or a random subspace (center right), as determined by the local linearization of the dynamics, or without subspace constraints (far right). Gray lines show chance level overlap defined as the expected overlap with a randomly chosen subspace occupying 25% of the full space (i.e. 5 dimensions). Compare with Fig. 4c for the analogous plots for linear networks (though with non-instantaneous inputs, and performance-matched levels of noise, see also Supplementary Information S4) and with Fig. 5f and Extended Data Fig. 10f,g for analogous plots of linear neural networks fitted to experimental data.

**Extended Data Fig. 8.**
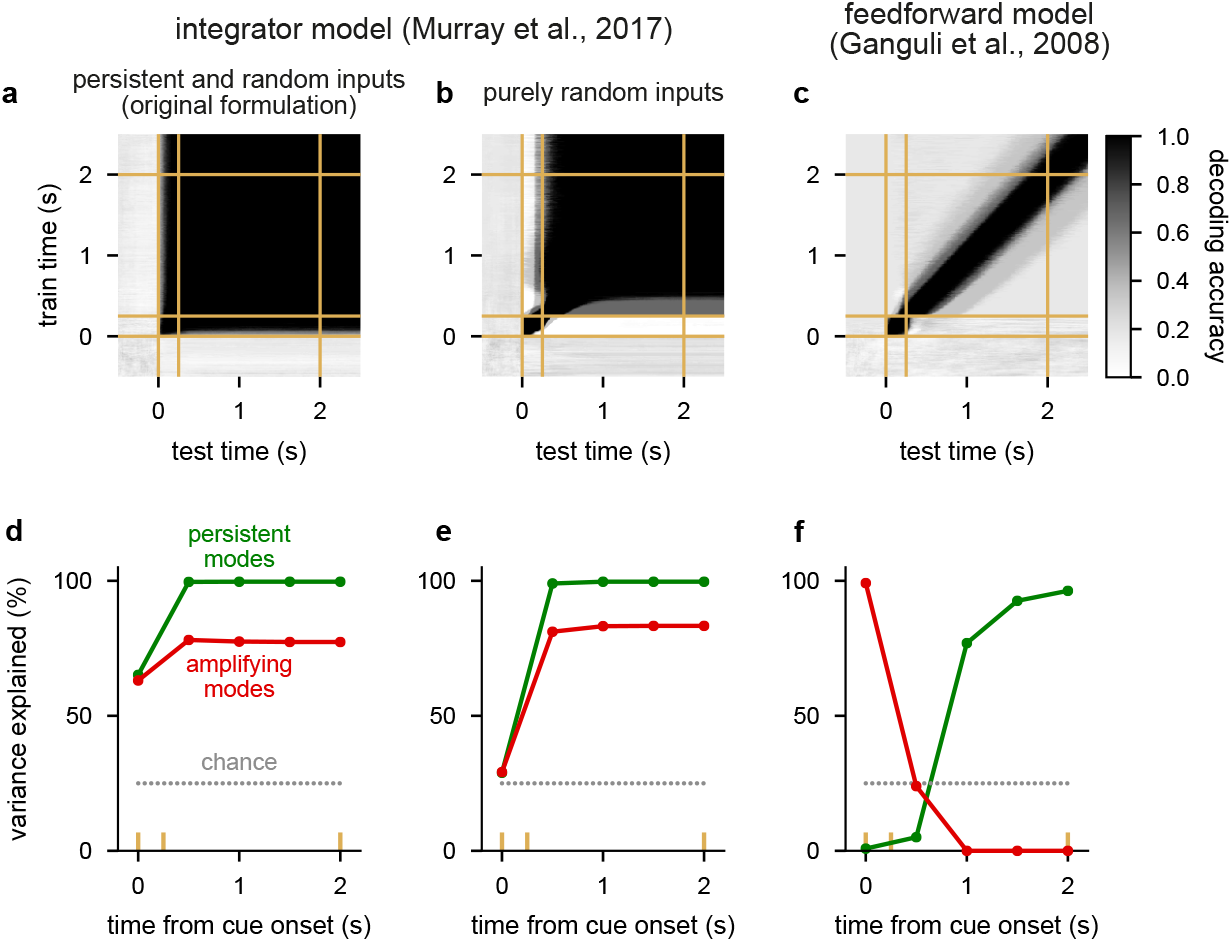
Analysis of two variants of an integrator model and feedforward model. **a**, Cross-temporal decoding of model neural activity (cf. Fig. 2e,k, Fig. 4b, and Fig. 5c) for a linear integrator model^6^ (cf. Extended Data Fig. 1c and Methods 1.5). Yellow lines indicate cue onset, offset, and go times. **b**, Same as **a** for the same model but for inputs aligned with purely random directions (as opposed to inputs aligned with both persistent and random directions as in the original formation of Ref.^6^). **c**, Same as **a** but for a linear feedforward network model^21,27^ (cf. Extended Data Fig. 1d). **d**, Percent variance of responses explained by the subspace spanned by either the 25% most persistent (green) or 25% most amplifying (red) modes as a function of time for the linear integrator model from **a** (cf. Fig. 4c,b, Fig. 5f, and Fig. 6e). Yellow lines indicate cue onset, offset, and go times. Gray dotted line shows chance level overlap with a subspace spanned by 25 random orthogonal directions. **e**, Same as **d** for the same model but for inputs aligned with purely random directions. **f**, Same as **d** but for a linear feedforward network model^21,27^.

**Extended Data Fig. 9.**
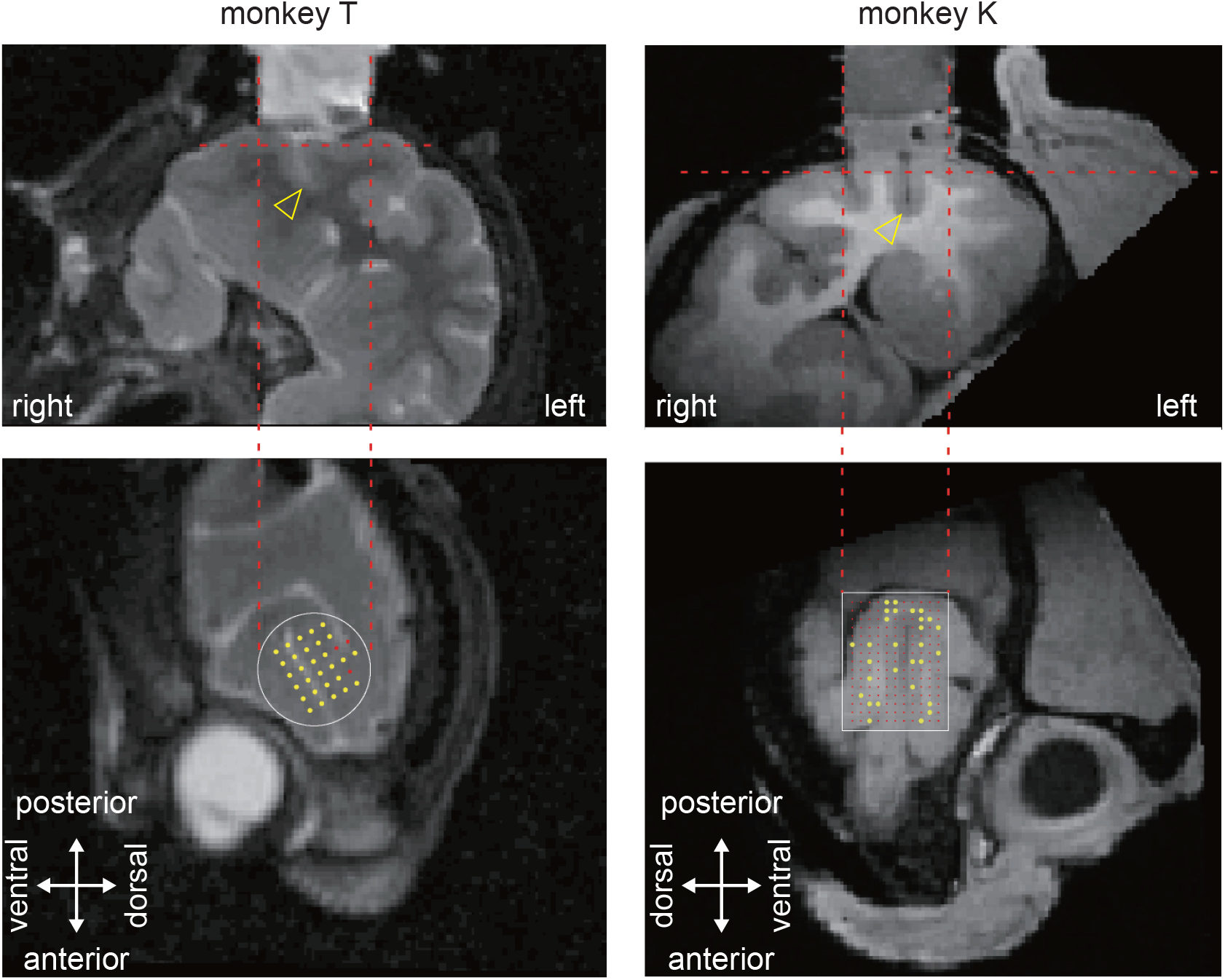
Recording locations for the two monkeys. Left: recording locations in monkey K (T1-weighted image). In order to image the interior of the chamber, we filled the chamber with cut cottons soaked in iodine. In the upper picture, the yellow arrow indicates the principal sulcus. In the bottom picture, locations of the 11 by 15 grid holes were superimposed over the MR picture. Right: recording locations in monkey T (T2-weighted image). The bottom picture shows the location for the grid of the 32 semi-chronic electrodes. Yellow dots indicate electrode penetrations and recording sites, red dots indicate non-visited sites.

**Extended Data Fig. 10.**
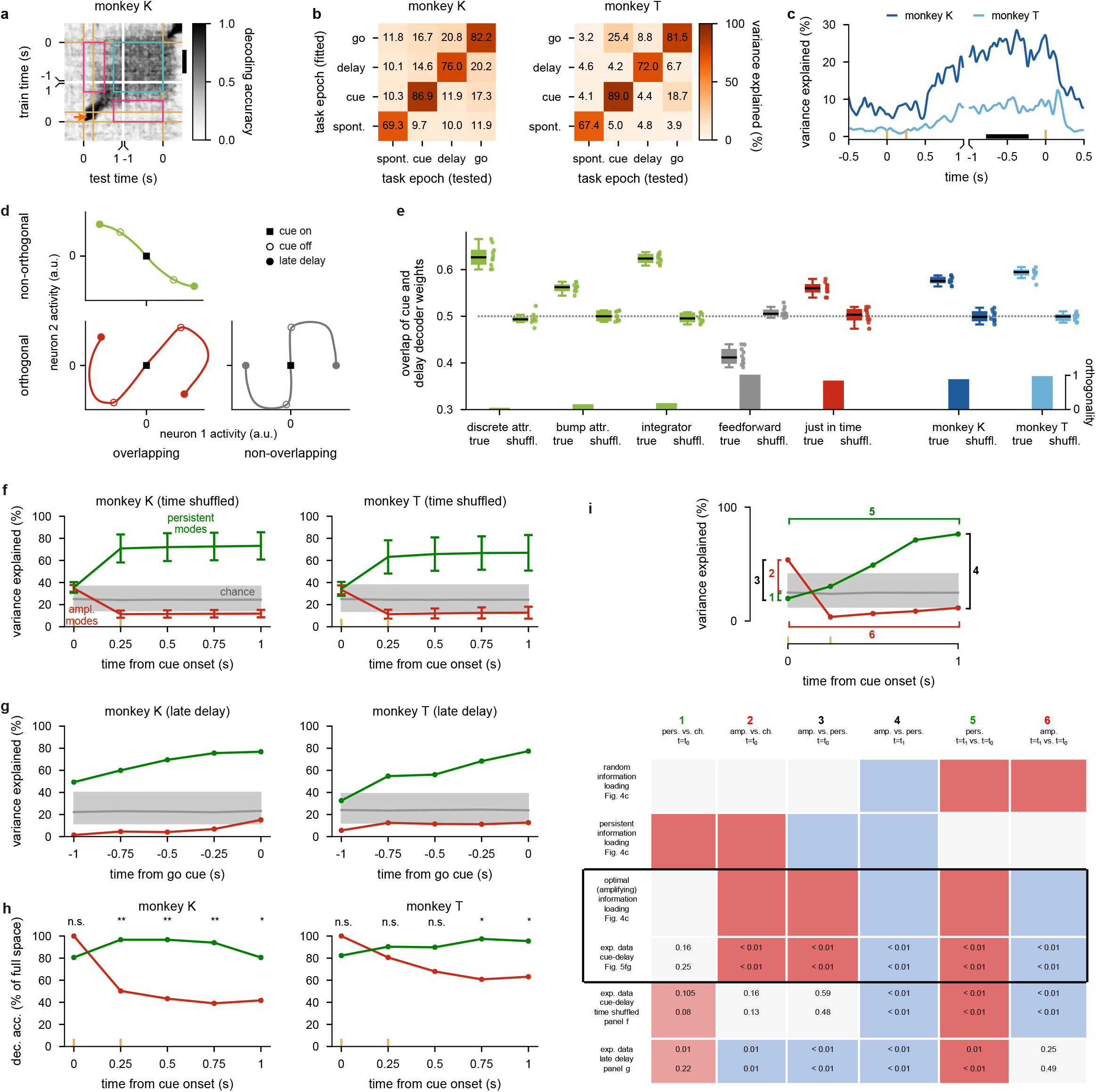
Supplemental analysis of experimental data and comparison to models. **a**, Cross-temporal decoding analysis for monkey K (cf. Fig. 5c for the same analysis for monkey T and for explanation of plotting scheme and annotations). **b**, Subspace overlap between different task epochs, measured as the percent variance explained (PVE) by projecting neural activity from one task epoch (tested) through the top 10 PCs of another task epoch (fitted). Diagonal elements show the PVE within each task epoch. We show results for monkey K (left) and monkey T (right). **c**, Time course of overlap with delay epoch subspace, measured as the percent variance explained by the top 2 PCs obtained from delay period activity (black bar shows time period of activity from which these PCs were obtained) on held-out test data taken in different time bins. This metric is called the alignment index^64^ and is very similar to that used in Ref. 6 (Methods 1.7.3). We show mean (over 10 different data splits) results for both monkeys. Yellow ticks on horizontal axis indicate cue onset, cue offset, and go times. **d**, Schematic of 3 different hypothetical scenarios for the relationship between cue and late delay activities (panels), illustrated in neural dynamics for 2 neurons and 2 cue conditions. Colored traces show neural trajectories, black squares indicate cue onset, open circles indicate cue offset, and filled circles show late delay activity. Left vs. right: populations encoding the cue during cue and late delay periods are overlapping vs. non-overlapping, respectively. Top vs. bottom: cue and delay activities are non-orthogonal vs. orthogonal, respectively. (Note that we are not showing dynamics for non-overlapping, non-orthogonal dynamics because no overlap necessarily implies orthogonality.) **e**, Relationship between cue and late delay activities in various different models and our experimental recordings (x-axis). Top: population overlap measured as the mean difference between cue and delay epoch decoder weights (left for each model and data) and, as a control, when randomly shuffling decoder weights across neurons (right for each model and data) (Methods 1.7.6). Box plots show medians (black lines), quartiles (boxes), and 1.5 times the inter-quartile range (whiskers). Dotted gray line shows chance level overlap. Bottom: orthogonality measured as 1 minus the mean overlap between cue and delay epochs (given by the corresponding elements of the subspace overlap matrices shown in panel **b** and Extended Data Fig. 11c, center right). The discrete attractors, bump attractor, and integrator models show high overlap but low orthogonality. The simple feedforward network shows high orthogonality but low overlap (note that recurrent networks with embedded feed-forward connectivity^21^ may show high overlap). The just-in-time network shows high overlap and orthogonality, similar to the experimental data in both monkeys. **f–g**, Same analysis as in Fig. 5f, but either after randomly shuffling data across time (but consistently across conditions and neurons, and applied to the same time period as in the main analysis; **f**, see also Methods 1.4.3), or applied to the late delay time period (without across-time shuffling) in which we do not expect information loading dynamics (**g**). **h**, Decoding of stimulus information within the subspace spanned by either the 25% most persistent modes (green), or the 25% most amplifying modes (red) in the linear neural networks shown in Fig. 5f relative to decoding accuracy using the full space. Comparisons use two-sided permutation tests (*, *p <* 0.05; **, *p <* 0.01; n.s., not significant; see Methods 1.8) **i**, Top inset: original data analysis of overlaps repeated from Fig. 5f to indicate the comparisons (colored numbers) we show in the table below (numbered columns). Bottom: table showing p-values (in each cell for experimental data, top: monkey K, bottom: monkey T) from two-sided permutation tests for each comparison of the main analysis (row 4, repeated from the main text associated with Fig. 5f) and the control analyses shown in panels **f** and **g** of this figure (rows 5–6). Top 3 rows show predictions for the sign of each comparison under different information loading strategies in unconstrained linear networks (Fig. 4c): using inputs aligned with random directions (1st row), persistent directions (2nd row), or the most amplifying directions (3rd row). In the column headings, pers., amp., and ch. respectively refer to overlap with most persistent, most amplifying and random subspaces (chance), *t*_0_ refers to the beginning of the analysis time window, i.e. cue onset (rows 1–5) or 1 s before the timing of the go cue (row 6), and *t*_1_ = *t*_0_ + 1 s refers to the end of the analysis time window. The colored numbers above each column correspond to the comparisons shown in the inset above the table. Gray indicates no significant difference between data points, red and blue indicate a significant difference for both monkeys where the first data point is respectively greater or smaller than the second data point, and pale red indicates a significant difference for one of the two monkeys (see Methods 1.8).

**Extended Data Fig. 11.**
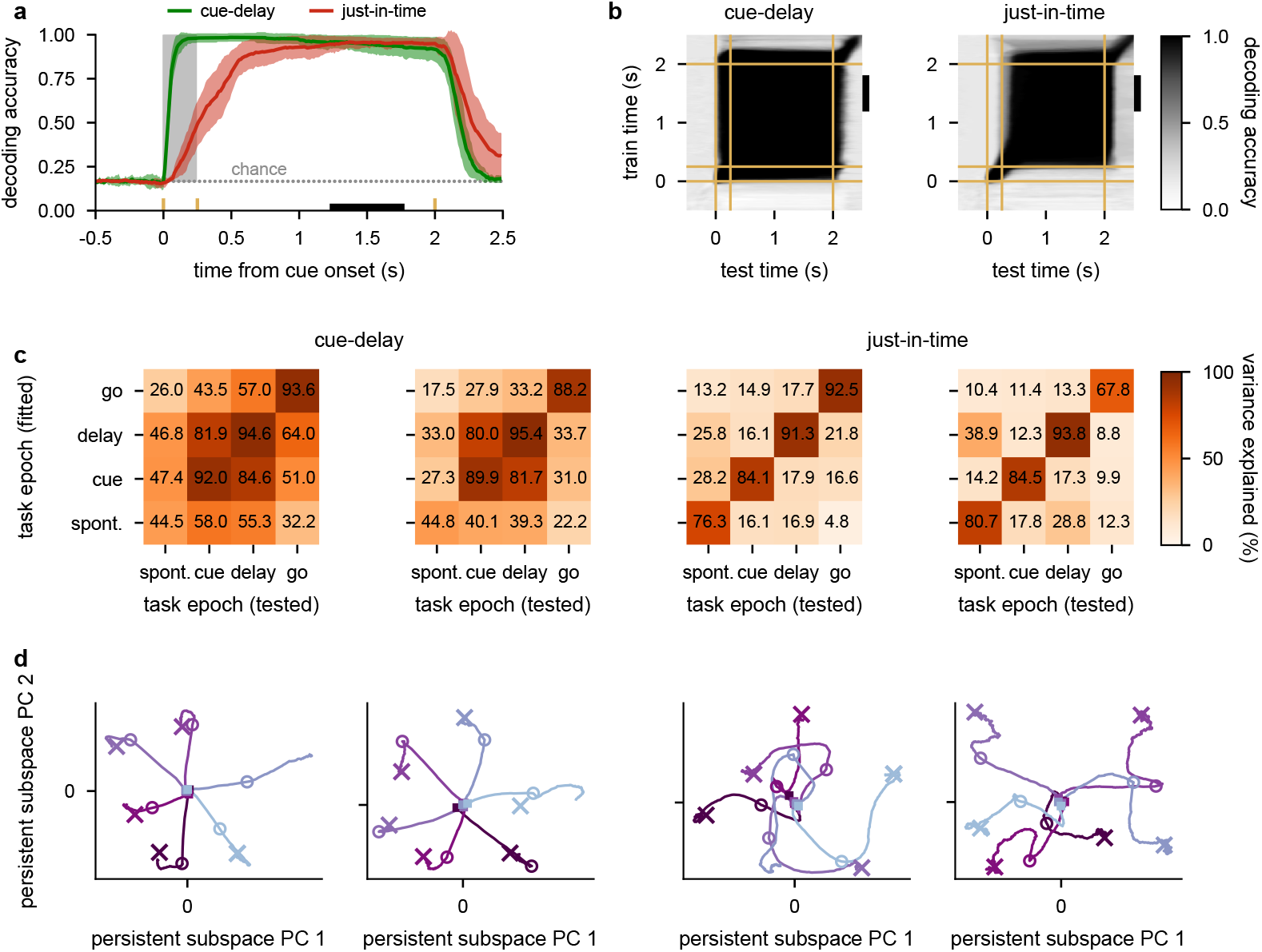
Cue-delay and just-in-time trained networks. **a–b**, Same as Fig. 6c green and red, and Fig. 6d left and right, but with a regularisation strength of 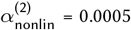 used during training (Methods 1.3.2). **c**, Subspace overlap between different task epochs, measured as the percent variance explained (PVE) by projecting neural activity from one task epoch (tested) through the top 4 PCs of another task epoch (fitted; cf. Extended Data Fig. 12d, Extended Data Fig. 13d, and Extended Data Fig. 10b). Diagonal elements show the PVE within each task epoch. We show results for cue-delay (left two panels) and just-in-time trained networks (right two panels) trained with either a regularisation strength of 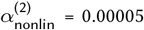 (left panel for each model, as in Fig. 6) or 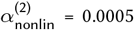 (right panel for each model, as in panels **a**–**b**). **d**, Neural activity plotted in the top two PCs of delay-epoch activity for all 6 initial conditions for cue-delay and just-in-time trained networks for each of the network-regularization combinations shown in **c** (cf. Extended Data Fig. 2b–d and f–h.) Purple traces show state-space trajectories, squares indicate cue onset, open circles indicate cue offset, and crosses indicate asymptotically stable fixed points, colors indicate cue condition as in Fig. 5e.

**Extended Data Fig. 12.**
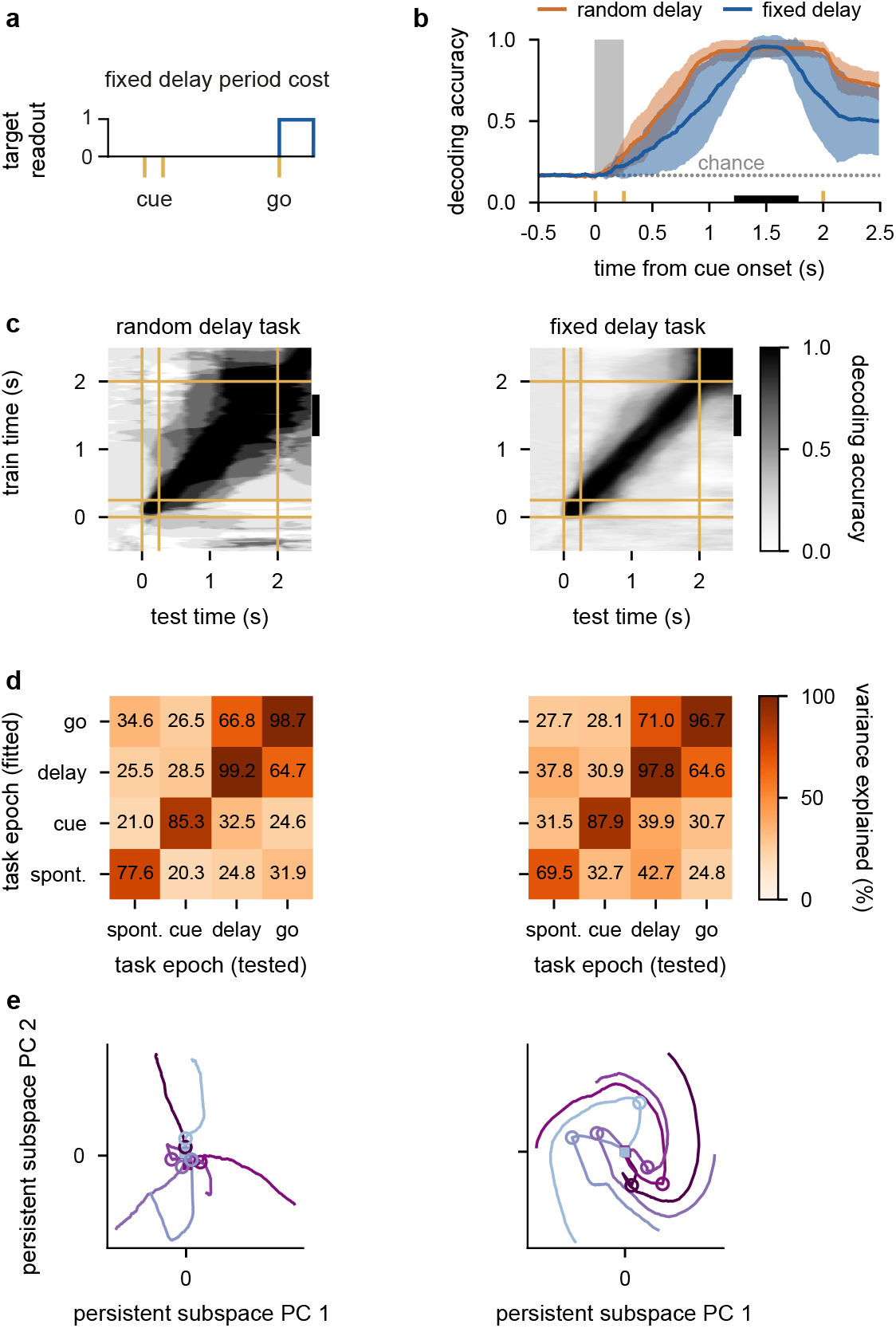
After-go-time trained networks. **a**, Cost function for after-go-time training on the fixed delay task (Methods 1.3.3). Cue onset, cue offset, and go cue times are indicated by the yellow vertical lines. The boxcar shows the interval over which stable decoding performance was required (i.e. the cost was only applied after the go cue). **b–c**, Same as Fig. 6c orange and Fig. 6d center, but with a regularisation strength of 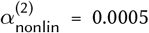 used during training and when either a random (**b** orange, **c** left) or a fixed delay task is used (**b** blue, **c** right, Methods 1.7.4). **d**, Subspace overlap between different task epochs, measured as the percent variance explained (PVE) by projecting neural activity from one task epoch (tested; cf. Extended Data Fig. 11c, Extended Data Fig. 13d, and Extended Data Fig. 10b) through the top 4 PCs of another task epoch (fitted) for the networks shown in **b**–**c**. Diagonal elements show the PVE within each task epoch. **e**, Neural activity plotted in the top two PCs of delay-epoch activity for all 6 initial conditions for random delay (left) and fixed delay (right) trained networks (cf. Extended Data Fig. 2b–d and f–h; and Extended Data Fig. 11d.) Purple traces show state-space trajectories, squares indicate cue onset, open circles indicate cue offset, and colors indicate cue conditions as in Fig. 5e. (Note that the absence of crosses indicates the absence of asymptotically stable fixed points.)

**Extended Data Fig. 13.**
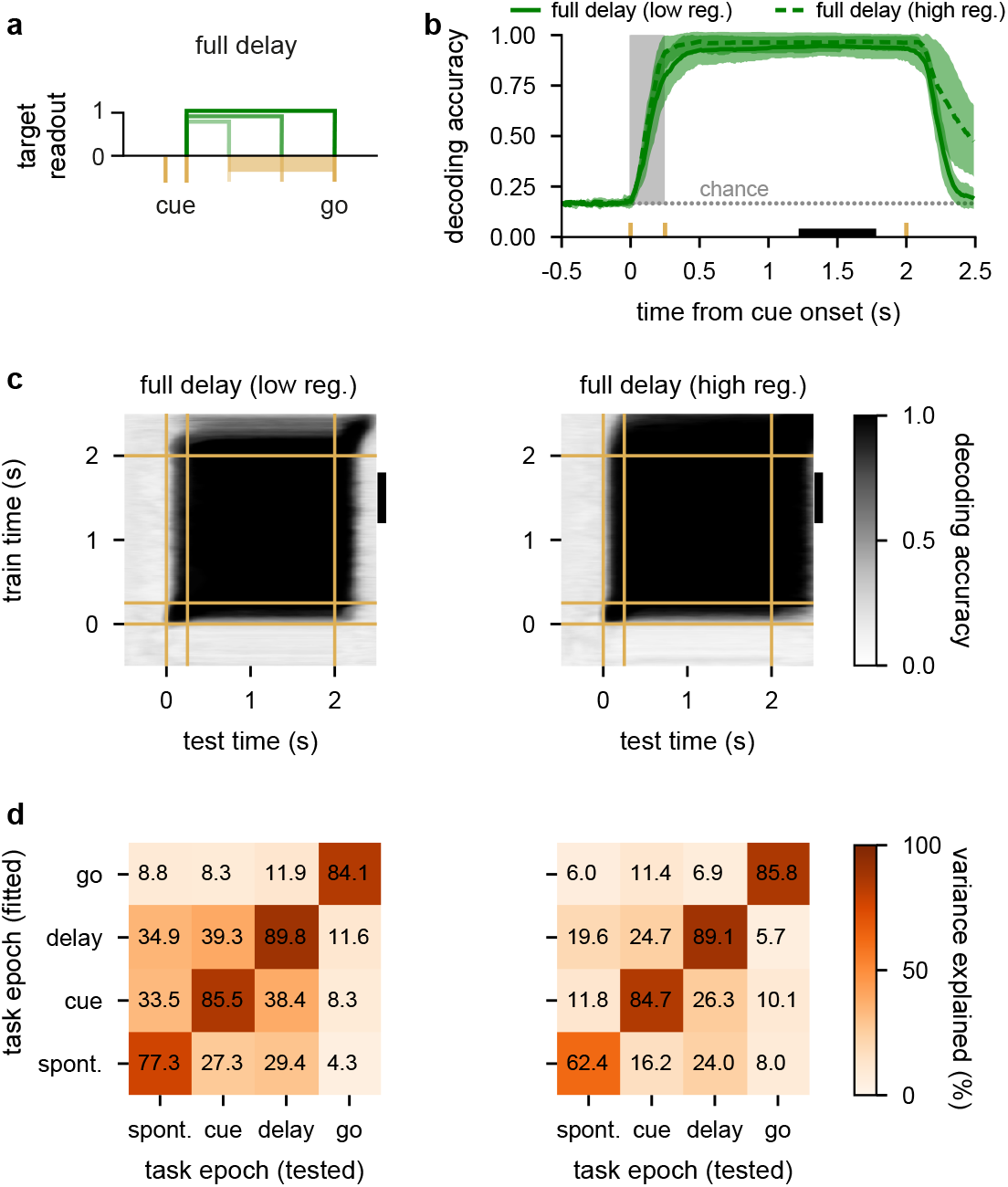
Full-delay trained networks. **a**, Cost function for full-delay training on the random delay task (Methods 1.3.3). Yellow ticks indicate cue onset and offset times, the yellow bar indicates range of go times in the variable delay task. Boxcars show intervals over which stable decoding performance was required in three example trials with different delays (Methods 1.3.3). **b–c**, Same as Fig. 6c–d, but when training with the full-delay cost with a regularisation strength of 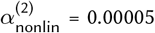 (**b** solid, **c** left) or 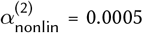 (**b** dashed, **c** right, Methods 1.7.4). **d**, Subspace overlap between different task epochs, measured as the percent variance explained (PVE) by projecting neural activity from one task epoch (tested; cf. Extended Data Fig. 11c, Extended Data Fig. 12d, and Extended Data Fig. 10b) through the top 4 PCs of another task epoch (fitted) for the networks shown in **b**–**c**. Diagonal elements show the PVE within each task epoch.

## Notes

### Competing Interest Statement

The authors have declared no competing interest.

### Summary of Updates

Updated figures 1 and 4

