## Supplementary information for "Optimal information loading into working memory in prefrontal cortex explains dynamic coding"

**S1 Comparison of previous models of working memory with task-optimized networks . . . 2**

**S2 Analysis of linear integrator networks . . . 3**
S2.1 Linear integrator networks 3
S2.2 The persistent mode 4
S2.3 The energy of the system 5
S2.4 Optimal information loading: maximising asymptotic persistent overlap with fixed initial norm 5
S2.5 Optimal information loading: achieving fixed asymptotic persistent overlap with a minimal initial norm 6
S2.6 Optimal information loading: maximising asymptotic energy with fixed initial norm 6
S2.7 Optimal information loading: noise-robust decoding 7
S2.8 Optimal information loading: minimising energy with fixed persistent overlap 8

**S3 Analysis of canonical nonlinear systems with two stable fixed points . . . 9**
S3.1 A local linearization approach based on linearizing around the ground state instead of an attractor 9
S3.2 Most persistent and amplifying directions for nonlinear dynamics 10
S3.3 Locally most amplifying modes predict optimal inputs, analogous to linear networks 10

**S4 Analysis of the nonlinear attractor networks of Fig. 2 . . . 11**
S4.1 The relationship between subspaces extracted from nonlinear dynamics and by local linearization 11
S4.2 Locally most amplifying modes predict optimal inputs, analogous to linear networks 11
S4.3 Overlap with most persistent mode in linear networks predicts decoding accuracy in nonlinear networks 11
S4.4 Fitting linear dynamics reveals information loading strategies in nonlinear networks 12

**S5 Fisher information in task-optimized ring attractor networks . . . 13**

**References . . . 14**

The main insight of our work is that optimal information loading relies on inputs aligned with the most amplifying mode of network dynamics, which in turn accounts for the combination of stable and dynamic coding seen in experiments. In Section S1, we first show that this combination does not arise in previously proposed models of working memory with hand-crafted connectivities, but does in task-optimized neural networks. To understand why network optimization leads to this behavior, in Section S2, we provide mathematically rigorous derivations of optimal information loading for linear network dynamics. However, these derivations—and indeed, the very concept of the most amplifying mode—do not readily apply to nonlinear networks, which are likely to be more relevant for understanding cortical circuits. Therefore, in Section S3 we generalize the concept of the most amplifying mode to nonlinear dynamical systems (through a local linearization around the ground state) and show that in a simple canonical nonlinear attractor system it offers the same qualitative advantage for information loading as in linear networks. In Section S4, we demonstrate the same advantage in high-dimensional multi-attractor nonlinear networks, show that the abstract notion of information loading performance we used in simple systems predicts a practically more relevant measure of performance in these networks, and that the dynamical signatures we established for identifying optimal information loading in linear networks can still be used even when the real dynamics of a circuit are nonlinear, and only indirectly accessible to us by the neural responses they generate. As there are no readily available analytical techniques that could be directly applied to the problem of information loading in nonlinear dynamical systems, our analyses in Sections S3 and S4 are mostly based on numerical simulations of specific nonlinear dynamical systems where we have ground truth knowledge of the dynamics and the information loading strategy that is optimal for them. Finally, in Section S5, we show with analytical derivations that optimizing a ring attractor with a negative cosine loss on the difference between true and decoded angles (as we do in the main text) is approximately equivalent to optimizing the population Fisher information about angle in the network.

### S1 Comparison of previous models of working memory with task-optimized networks

We simulated four, previously proposed representative models of working memory, as well as one of the task-optimized networks developed here for comparison (the just-in-time network in Fig. 6d, right).

First, we simulated the 'bump' attractor network of Ref. 1: a recurrent network in which neurons are indexed by their preferred directions and are connected recurrently by a circulant weight matrix, such that the connection strength between two excitatory neurons only depended (as a circular Gaussian) on the difference of their preferred directions, and there was a single inhibitory neuron providing global inhibition (Methods 1.5). (Note that this network thus represents a paradigmatic case of quasi-symmetric connectivity.) In the absence of inputs, this network is able to maintain the memory of a saccade direction via a 'bump' of activities: its intrinsic dynamics (mediated by the recurrent interactions between its neurons) converge to a steady state pattern in which the response of neurons is described by a Gaussian-like ('bump') function of the difference between their preferred direction and the maintained direction. To understand information loading and static vs. dynamic coding in this network, it is useful to adopt a state space view of its dynamics (cf. Fig. 3). Information about the to-be-memorised direction (the stimulus cue) is loaded into this network by a transient external input. During the cue period, this transient input drives the network (Extended Data Fig. 1a, left, pale purple line and arrow) into a suitable state (Extended Data Fig. 1a, left, pale purple circle). Then, during the delay period, in the absence of the cue, its intrinsic dynamics (Extended Data Fig. 1a, left, dark purple line), are 'attracted' into a cue-specific steady state (corresponding to a bump of activities; Extended Data Fig. 1a, left, black cross). This naturally accounts for selective persistent activity (Extended Data Fig. 1a, center). However, the state into which the external input drives network activities, i.e. the initial state during the delay period, already has large overlap with the steady state to which the dynamics eventually converge (Extended Data Fig. 1a, left, pale purple line points in the direction of the black cross). In other words, the network performs simple 'pattern completion'[2], whereby this overlap is only slightly improved until it becomes perfect (Extended Data Fig. 1a, left), with the response of each neuron quickly reaching its steady-state and maintaining it throughout most of the delay period (Extended Data Fig. 1a, center). As a result, neurons show limited transient activity during the delay period (Extended Data Fig. 1a, center), and cross-temporal decoding reveals stable coding throughout the whole trial, lacking the characteristic dynamic coding seen in experimental data (compare Fig. 1b to Extended Data Fig. 1a, right).

We also simulated the closely related 'discrete' attractor network of Ref. 1. In this network, neurons are assigned to a finite number of discrete clusters (one cluster for each cue condition), with strong recurrent connections between neurons belonging to the same cluster, and weaker connection between neurons belonging to different clusters (Methods 1.5). Thus, instead of maintaining bumps of activity, the steady states of this network correspond to one of the clusters being active while all others exhibiting weak background activity. Other than that, the same principles of simple pattern completion apply to information loading that we saw in the bump attractor network (Extended Data Fig. 1a). Therefore, this network is also characterized by the absence of dynamic coding (Extended Data Fig. 1b,c).

Recent work has suggested that linear integrator networks, that are based on similar principles as nonlinear attractor networks but use linear dynamics, can exhibit activity transients similar to those seen in experiments[3] (see also[4]). In line with earlier work, when simulating such models, we also found activity transients to emerge when cue-dependent inputs included a random component in addition to the 'usual' steady state-aligned component that ensured pattern completion

(Methods 1.5). Because the random component in these networks is designed to be orthogonal to the steady state-aligned component, the dynamics of the network take time to relax to the subspace spanned by the cue-dependent steady states (the coding subspace; Extended Data Fig. 1c, left), and neurons exhibit transient activities during this relaxation (Extended Data Fig. 1c, middle). However, precisely because this relaxation thus takes place in a subspace orthogonal to the persistent subspace, it does not affect stimulus coding during the delay. Thus a delay-trained decoder will always generalize perfectly to the cue period. Indeed, we found that cross-temporal coding demonstrated strongly stable coding (Extended Data Fig. 1c, right, also see Extended Data Fig. 8a)—just as in the classical attractor networks seen above. We were able to achieve more dynamic coding in a modified version of this model, in which the random component of the inputs completely dominated (Methods 1.5.3, Extended Data Fig. 8b). In other words, this modified model consisted of unconstrained (non-normal) connectivity and purely random inputs (similar to Fig. 4, 'random'). Nevertheless, our neural measure of optimal information loading, based on the time course of the overlap of activities with the most amplifying and most persistent modes (Extended Data Fig. 8d,e) still revealed a fundamental difference from experimental data (cf. Fig. 5f) and task-optimized neural networks (cf. Fig. 6e).

A major class of alternative models of working memory use essentially feedforward dynamics[5,6]. We thus simulated a simple, purely feedforward network that captured the essence of such dynamics (Methods 1.5). As expected, this network did not exhibit any steady state and inputs that stimulated the beginning of the feedforward chain resulted in long transients that rapidly transitioned into activity patterns that were orthogonal to these inputs (because they involved the firing of an altogether different group of neurons at the end of the chain)—after which, ultimately, all activities decayed to zero (Extended Data Fig. 1d, left and middle). (Note that the orthogonality of successively expressed activity patterns, and the eventual decay of activities, would remain the same even in the more general case when the nodes of the feedforward chain are distributed patterns of neural activities, rather than individual neurons.) As a consequence, this network showed strong dynamic coding with cross-temporal decoding being high only between neighbouring time-points, but no stable coding during the delay period (Extended Data Fig. 1d, right).

Finally, for comparison, we show the same plots for a representative example of one of our task-optimized networks (Extended Data Fig. 1e; see also Fig. 6, 'just-in-time'). In this network, inputs are also largely orthogonal to the late delay activities, as in the feed-forward networks, but eventually converge to an attractor maintaining information during the late delay period, as in attractor networks (Extended Data Fig. 1e, left). As a consequence, neurons show rich transient dynamics (Extended Data Fig. 1e, middle), and the network exhibits the experimentally described combination of dynamic and stable coding (Extended Data Fig. 1e, right).

### S2 Analysis of linear integrator networks

In this section we provide analytical derivations showing that optimal information loading is achieved by the most amplifying mode in linear integrator networks. We first define linear integrator networks and introduce the basic mathematical concepts for understanding their dynamics (Section S2.1). We then define the persistent mode, a direction in the state space of linear integrator networks that is critical for characterising their performance (Section S2.2). We also define the energy of the system as a quantity useful for both constraining optimisation and defining optimal information loading (Section S2.3). We then define optimal information loading in five seemingly different ways (via five different constrained optimization problems), but show that all definitions lead to the same solution (at least approximately):

1. The initial condition with a fixed norm that achieves the highest asymptotic overlap with the persistent mode (Section S2.4).
2. The initial condition with minimal norm that achieves a fixed asymptotic overlap with the persistent mode (Section S2.5).
3. The most amplifying mode, defined as the initial condition with a fixed norm that evokes the largest asymptotic energy (Section S2.6).
4. The initial condition with a fixed norm that provides maximal 'total' decodability in a *noisy* linear integrator over an asymptotically long time horizon (Section S2.7).
5. The initial condition that achieves a fixed asymptotic overlap with the persistent mode while using the least amount of (asymptotic) energy (Section S2.8).

#### S2.1 Linear integrator networks

We define a linear integrator network as a linear dynamical system

$$\dot{\mathbf{x}}(t) = \mathbf{J}\,\mathbf{x}(t) \tag{S1}$$

with a Jacobian $\mathbf{J}$, which we define using its eigen-decomposition:

$$\mathbf{J} = \mathbf{V}\,\mathbf{\Lambda}\,\mathbf{U} \tag{S2}$$

where $\mathbf{\Lambda}$ is a diagonal matrix containing the eigenvalues of $\mathbf{J}$, $\lambda_1, \ldots \lambda_N$ (where $N$ is the number of neurons in the network, and the eigenvalues are sorted in descending order of their real values, without loss of generality), and $\mathbf{V}$ is the matrix of the associated (right) eigenvectors of $\mathbf{J}$, $\mathbf{v}_{(1)}, \ldots \mathbf{v}_{(N)}$, as columns (unit-norm, without loss of generality). Conversely, $\mathbf{U} = \mathbf{V}^{-1}$ is the matrix of the corresponding left eigenvectors, $\mathbf{u}_{(1)}^\top, \ldots \mathbf{u}_{(N)}^\top$, as rows. Note that typical dynamics for a linear neural network (cf. Eq. 1) include a leak term and a time constant, $\tau$, such that $\mathbf{J} = \frac{1}{\tau}(\mathbf{W} - \mathbf{I})$, where $\mathbf{W}$ is the recurrent weight matrix of the network (and we ignored noise and temporally extended external inputs to the network in order to focus on the intrinsic deterministic component of its dynamics).

Note that the eigenvectors in $\mathbf{V}$ need not be linearly independent (orthogonal): they are only orthogonal in the special case when $\mathbf{J}$ is a normal matrix (e.g. symmetric), in which case $\mathbf{U} = \mathbf{V}^\top$, i.e. the left and right eigenvectors are identical. Otherwise, when $\mathbf{J}$ is non-normal, its eigenvectors are not orthogonal, and $\mathbf{U} \neq \mathbf{V}^\top$, i.e. the left and right eigenvectors are different. This will be important below for understanding the geometry of optimal information loading in the network.

In order for a linear dynamical system to act as an integrator network, we require that its top eigenvalue (eigenvalue with the largest real part), $\lambda_1$, is real, unique, and marginally stable

$$\lambda_1 = -\delta \text{ (with } \delta \to 0) \tag{S3}$$

and all of its other eigenvalues are properly stable with real parts that are

$$\mathrm{Re}(\lambda_i) \ll 0 \text{ (for } i > 1) \tag{S4}$$

(This is consistent with our procedure for constructing the linear integrator networks analysed in Figs. 3 and 4 and Extended Data Figs. 4 and 5, see Methods 1.4.1, and Table 1.)

Following standard textbook treatment[7], we often make use of the eigen-decomposition in Eq. S2 to write the state of the system in terms of its eigen-coefficients as

$$\mathbf{y}(t) = \mathbf{U}\,\mathbf{x}(t) \quad \text{, or, equivalently, } \mathbf{x}(t) = \mathbf{V}\,\mathbf{y}(t) \tag{S5}$$

which in turn allows us to rewrite the dynamics of the system in Eq. S1 as

$$\dot{\mathbf{y}}(t) = \mathbf{\Lambda}\,\mathbf{y}(t) \tag{S6}$$

Indeed, the solution to the linear dynamics of the system (Eq. S1) in the original space of neural activities is

$$\mathbf{x}(t) = e^{\mathbf{J}\,t}\,\mathbf{x}(0) \tag{S7}$$

which expresses complex interactions between neurons (in the first term). However, in the space of the eigen-coefficients, the solution of the dynamics (Eq. S6) can be conveniently written as a set of *independently* evolving components, each decaying exponentially to zero with a rate given by the corresponding eigenvalue:

$$y_i(t) = y_i(0)\; e^{\lambda_i\,t} \tag{S8}$$

In the following, we will be particularly interested in the asymptotic behavior of the system, $t \to \infty$. Given the marginally stable nature of the system (Eqs. S3 and S4), in this joint limit all but the first eigencoefficient decay to zero:

$$e^{\delta\,t}\,y_i(t) \to \begin{cases} y_1(0) & \text{for } i = 1 \\ 0 & \text{otherwise} \end{cases} \tag{S9}$$

### S2.2 The persistent mode

The persistent mode of the system is the direction in state space with which the system's state, $\mathbf{x}(t)$ is aligned in the asymptotic limit, $t \to \infty$. As such, a natural measure of the system's performance will be its position along this direction (Section S2.4).

We define the momentary direction of the system's state as

$$\bar{\mathbf{x}}(t) = \frac{\mathbf{x}(t)}{\sqrt{\mathbf{x}^\top(t)\;\mathbf{x}(t)}} \tag{S10}$$

which we can rewrite using the eigen-coefficients (Eq. S5) as

$$\bar{\mathbf{x}}(t) = \frac{\mathbf{V}\,\mathbf{y}(t)}{\sqrt{\mathbf{y}^\top(t)\ \boldsymbol{\Xi}\,\mathbf{y}(t)}} \tag{S11}$$

$$\text{where } \boldsymbol{\Xi} = \mathbf{V}^\top \mathbf{V} \tag{S12}$$

measures the overlap between the eigenvectors. Taking the asymptotic limit as defined above (Eq. S9), leads us to

$$\bar{\mathbf{x}}(t) \to \mathbf{v}_{(1)} \tag{S13}$$

In other words, asymptotically, the network expresses an activity pattern that is aligned with $\mathbf{v}_{(1)}$, the first *right* eigenvector of the Jacobian, which we thus call its *persistent mode*. In control theoretic terms, this shows (the unsurprising result) that the persistent mode of a linear integrator network is its most 'controllable' mode[8].

### S2.3 The energy of the system

As we will see below, an important quantity for constraining and understanding network dynamics is the 'energy' of the system, which is the total norm of neural responses integrated over some (potentially infinite) time window. We define the total 'energy' that the system has used up to time $t$ as:

$$\varepsilon(t) = \int_0^t \mathbf{x}^\top(t')\ \mathbf{x}(t')\ \mathrm{d}t' \tag{S14}$$

which we can re-write using the solution to the dynamics in Eq. S7 as

$$\varepsilon(t) = \mathbf{x}^\top(0) \int_0^t e^{\mathbf{J}^\top t'}\ e^{\mathbf{J}\,t'}\ \mathrm{d}t'\ \mathbf{x}(0) \tag{S15}$$

Using the eigen-coefficients (Eq. S5), and the overlap of eigenvectors as defined in Eq. S12, we can further rewrite this as

$$\varepsilon(t) = \int_0^t \mathbf{y}^\top(t')\ \boldsymbol{\Xi}\,\mathbf{y}(t')\ \mathrm{d}t' \tag{S16}$$

Using Eq. S8 to solve the dynamics in the space of eigen-coefficients analytically allows us to write the energy in closed form as an explicit function of the initial condition:

$$\varepsilon(t) = \mathbf{y}^\top(0)\ \boldsymbol{\Omega}(t)\ \mathbf{y}(0) \tag{S17}$$

$$\text{where } \Omega_{ij}(t) = \frac{\Xi_{ij}}{\lambda_i + \lambda_j}\ \left[e^{(\lambda_i+\lambda_j)\,t} - 1\right] \tag{S18}$$

Taking the asymptotic limit (Eq. S9) of Eq. S15, we obtain in the original space of neural responses

$$\varepsilon(t) \to \varepsilon_\infty = \mathbf{x}^\top(0)\ \mathbf{Q}\,\mathbf{x}(0) \tag{S19}$$

$$\text{where } \mathbf{Q} = \int_0^\infty e^{\mathbf{J}^\top t}\, e^{\mathbf{J}\,t}\,\mathrm{d}t \tag{S20}$$

is the so-called 'observability' Gramian of the dynamics[8]. We can once again rewrite this by substituting the asymptotic eigen-coefficients of marginally stable dynamics (Eq. S9) into Eq. S17 as

$$\varepsilon_\infty = \frac{1}{2\,\delta}\ y_1^2(0) \propto \mathbf{x}^\top(0)\ \mathbf{u}_{(1)}\,\mathbf{u}_{(1)}^\top\,\mathbf{x}(0) \tag{S21}$$

where $\mathbf{u}_{(1)}$ is the first column of $\mathbf{U}$, i.e. the first *left* eigenvector of $\mathbf{J}$.

### S2.4 Optimal information loading: maximising asymptotic persistent overlap with fixed initial norm

We now study the effects of using different inputs on the asymptotic behaviour of the system. For the purposes of the analyses presented here, by 'input', we mean the initial condition of the system, $\mathbf{x}(0)$. (For numerical results with temporally extended inputs, see Fig. 4, and also Fig. 6 and Extended Data Figs. 11–13 for nonlinear networks using temporally extended inputs). In the previous subsection we saw that asymptotically the system's state is always aligned with the persistent mode, independent of its initial condition. However, the precise point along that mode, i.e. the system's overlap with the persistent mode, still depends on the initial condition.

We define the momentary overlap with the persistent mode as

$$p(t) = \mathbf{v}_{(1)}^{\top}\,\mathbf{x}(t) \tag{S22}$$

Using the eigen-coefficients (Eq. S5), we once again rewrite this in closed form as an explicit function of the initial condition:

$$p(t) = \sum_i y_i(0)\;\Xi_{i1}\,e^{\lambda_i\,t} \tag{S23}$$

and take the asymptotic limit (Eq. S9) to obtain (see also Ref. 9 for an analogous derivation):

$$p(t) \to p_\infty = e^{-\delta\,t}\,y_1(0) \propto \mathbf{u}_{(1)}^{\top}\,\mathbf{x}(0) \tag{S24}$$

In order to study optimal information loading, we want to identify the initial condition, $\mathbf{x}(0)$, that achieves the maximal asymptotic overlap with the persistent mode. As the system has linear dynamics, it is obvious that $p(t)$ simply scales linearly with the norm of $\mathbf{x}(0)$. Therefore, to make our question meaningful, we have to introduce some additional constraint that prevents the trivial solution of an input with an infinitely large norm. We first consider simply a norm-constraint directly on $\mathbf{x}(0)$. This leads to solving the following optimization problem

$$\mathbf{x}^*(0) = \underset{\mathbf{x}(0)}{\operatorname{argmax}}\,\underbrace{\mathbf{u}_{(1)}^{\top}\,\mathbf{x}(0)}_{\propto\,p_\infty},\ \text{s.t.}\ \left\|\mathbf{x}(0)\right\|_2 = 1 \tag{S25}$$

of which the solution is trivially

$$\mathbf{x}^*(0) = \pm\mathbf{u}_{(1)} \tag{S26}$$

Thus, optimal information loading is achieved by the first *left* eigenvector of $\mathbf{J}$ (which, as we saw above, will *not* be identical to the first *right* eigenvector, $\mathbf{v}_{(1)}$, i.e. the persistent mode itself, in the general case, when $\mathbf{W}$—and thus $\mathbf{J}$—is non-normal).

### S2.5 Optimal information loading: achieving fixed asymptotic persistent overlap with a minimal initial norm

Defining optimal information loading instead as the input that achieves a fixed asymptotic overlap with the minimal norm, i.e. as the solution to the optimization problem

$$\mathbf{x}^*(0) = \underset{\mathbf{x}(0)}{\operatorname{argmin}}\,\left\|\mathbf{x}(0)\right\|_2,\ \text{s.t.}\ p_\infty = \text{const.} \tag{S27}$$

trivially leads to the same solution (up to multiplicative scaling) as that which we saw in Section S2.4, as it amounts to optimising the same Lagrangian:

$$\mathbf{x}^*(0) = \pm\mathbf{u}_{(1)} \tag{S28}$$

We will also consider below a third way to constrain the optimization problem, via its 'energy' (time integrated-norm), rather than just the norm of its initial condition (Section S2.8).

### S2.6 Optimal information loading: maximising asymptotic energy with fixed initial norm

The asymptotic energy we introduced above (Eq. S21) allows us to relate optimal information loading to a well-known concept in control theory: the most amplifying mode, which is the eigenvector of the observability Gramian associated with the largest eigenvalue[8,10,11]. Crucially, the most amplifying mode can also be defined as the initial condition with a unit norm that evokes the largest asymptotic energy, i.e. as the solution to the optimization problem

$$\mathbf{x}^*(0) = \underset{\mathbf{x}(0)}{\operatorname{argmax}}\,\underbrace{\mathbf{x}^{\top}(0)\;\mathbf{u}_{(1)}\,\mathbf{u}_{(1)}^{\top}\,\mathbf{x}(0)}_{\propto\,\mathcal{E}_\infty},\ \text{s.t.}\ \left\|\mathbf{x}(0)\right\|_2 = 1 \tag{S29}$$

which is straightforwardly given as

$$\mathbf{x}^*(0) = \pm\mathbf{u}_{(1)} \tag{S30}$$

Comparing Eq. S30 with Eq. S26 reveals that, in the case of 1-dimensional linear integrators, the most amplifying mode achieves optimal information loading as it is equivalent to the top left eigenvector. The intuition behind this result is simple:

- Optimal information loading is defined (Section S2.4) as using an initial condition that (asymptotically) generates maximal activity along the persistent mode.
- The most amplifying mode is formally defined as the direction for initial conditions that results in the largest overall fluctuations (on an infinite time horizon; Eq. S29).
- In linear integrators, all fluctuations are eventually quenched, except those that are expressed along the persistent mode of the system (Eq. S9).
- Therefore, if an input direction generates large overall fluctuations, it must be because it achieves a large asymptotic overlap with the persistent mode.
- Therefore, the most amplifying mode must achieve the largest overlap with the persistent mode, i.e. it is the optimal information loading direction.

More importantly, as we show in Section S2.7, the most amplifying mode generalises to cases (multiple integration directions in non-perfect integrators) in which the left eigenvectors are not relevant, thus motivating our use of the set of most amplifying directions rather than left eigenvectors to define optimal information loading throughout our analyses.

### S2.7 Optimal information loading: noise-robust decoding

We now study the more general problem of optimal inputs for general, stable linear dynamics, which are not necessarily exactly marginally stable, and may in fact have multiple dimensions along which they (approximately) integrate. To be closer to the actual (linear) networks we study in the main text (Fig. 4, Extended Data Fig. 5c,d, and Extended Data Fig. 8a,b,d,e), we consider stochastic dynamics of the form (cf. Eq. S1, but also see Eq. 2):

$$\dot{\mathbf{x}}(t) = \mathbf{J}\,\mathbf{x}(t) + \sigma\,\boldsymbol{\eta}(t) \tag{S31}$$

where $\boldsymbol{\eta}(t)$ is zero-mean, unit-variance white Gaussian noise, so that the total noise injected into the system has variance $\sigma^2$. We assume that the network has attained its (input-free) steady-state distribution before the stimulus arrives at $t = 0$, and the input $\mathbf{x}(0)$ is an instantaneous additive 'push' of the network state from whatever stochastic state it takes at $t = 0$ (rather than directly defining the network's state at $t = 0$ as before).

Considering stochastic network dynamics allows us to meaningfully define optimality in terms of decoding performance (rather than the somewhat arbitrary objective of maximising overlap with a particular direction in state space) — also in closer analogy with the main text. Following Ref. 5, we quantify decoding performance using the Fisher information. For this, we first note that as the dynamics is stochastic, the state of the system at any time is described by a probability distribution. Given that the dynamics is linear and the noise is Gaussian (Eq. S31), this distribution is also Gaussian:

$$\mathbf{x}(t) \sim \mathcal{N}\left(e^{\mathbf{J}\,t}\,\mathbf{x}(0)\,,\mathbf{C}\right) \tag{S32}$$

where $\mathbf{C} = \sigma^2 \int_0^\infty e^{\mathbf{J}\,t'}\,e^{\mathbf{J}^\top t'}\,\mathrm{d}t'$ is the stationary noise covariance of neural responses. (Note that the mean of the state distribution is the same as that for deterministic dynamics, Eq. S7, so that noise only contributes to the covariance, and conversely, the covariance does not depend on the input $\mathbf{x}(0)$.)

The ability to discriminate two inputs in the $\mathbf{x}(0)$ direction, $\mathbf{x}(0)$ and $\mathbf{x}(0)\ (1 + \Delta)$, at a later time $t$, is upper bounded by the Kullback-Leibler (KL) divergence between the two corresponding state distributions at that time. For normal distributions that only differ in their means (Eq. S32), the KL-divergence can be written simply as:

$$\mathrm{D}_{\mathrm{KL}}(t) = \frac{1}{2}\Delta^2\,\mathbf{x}^\top(0)\;\mathcal{I}(t)\;\mathbf{x}(0) \tag{S33}$$

where $\mathcal{I}(t)$ is the Fisher information matrix of the system at time $t$:

$$\mathcal{I}(t) = e^{\mathbf{J}^\top t}\,\mathbf{C}^{-1}\,e^{\mathbf{J}\,t} \tag{S34}$$

Indeed, this upper bound on decoding performance is saturated by a simple linear decoder, $\hat{\mathbf{x}}(t) = \mathbf{W}_{\mathrm{out}}\,\mathbf{x}(t)$ with readout weights

$$\mathbf{W}_{\mathrm{out}} = \mathbf{C}^{-1}\,e^{\mathbf{J}\,t}\,\mathbf{x}(0) \tag{S35}$$

Note that the optimal decoder is thus time-*dependent*.

Finally, to create a single scalar objective that can be optimised, we measure the 'total' decodability of the system over a suitably long (technically, infinite) time horizon:

$$\bar{\mathrm{D}}_{\mathrm{KL}} = \frac{1}{2}\Delta^2\,\mathbf{x}^\top(0)\;\bar{\mathcal{I}}\,\mathbf{x}(0) \tag{S36}$$

where $\bar{\mathcal{I}}$ is the total Fisher information matrix of the system (also called the 'Spatial Fisher memory matrix' in Ref. 5):

$$\bar{\mathcal{I}} = \int_0^\infty e^{\mathbf{J}^\top t}\, \mathbf{C}^{-1}\, e^{\mathbf{J}\, t} \mathrm{d}t \tag{S37}$$

which is symmetric and positive definite by construction. Note that Eq. S36 measures the (upper bound on) the diagonal of what we call the 'cross-temporal decoding matrix' in the main text, i.e. the performance of a time-*dependent* optimal linear decoder, whereas we (optimise and) use time-*independent* decoders for measuring decoding performance in stochastic integrators (roughly corresponding to a horizontal slice of the cross-temporal decoding matrix, Fig. 4a–b). As, in general, the optimal decoder is indeed time-dependent (Eq. S35), a single time-independent linear decoder will not be able to attain optimal performance. Nevertheless, for the integrators we study, $\bar{\mathrm{D}}_{\mathrm{KL}}$ is dominated by persistent activity, which in turn can be decoded by a fixed decoder, and so a time-independent linear decoder is able to achieve near-optimal performance.[1]

We can now define optimal information loading as the $\mathbf{x}(0)$ that maximizes

$$\mathbf{x}^*(0) = \underset{\mathbf{x}(0)}{\operatorname{argmin}}\, \underbrace{\mathbf{x}^\top(0)\, \bar{\mathcal{I}}\, \mathbf{x}(0)}_{\propto \bar{\mathrm{D}}_{\mathrm{KL}}}, \text{ s.t. } \|\mathbf{x}(0)\|_2 = 1 \tag{S38}$$

This is trivially solved by the top eigenvector of $\bar{\mathcal{I}}$, i.e. the eigenvector with the largest eigenvalue. Moreover, because $\bar{\mathcal{I}}$ is symmetric and positive definite (see above), this can be generalized to the first $k$ optimal information loading directions being the top $k$ (orthogonal) eigenvectors of $\bar{\mathcal{I}}$. Importantly, we observe that $\bar{\mathcal{I}}$, as defined in Eq. S37, is the so-called observability Gramian of the system[8], and thus its top $k$ eigenvectors are equivalent to the $k$ most amplifying directions of the system used in our analyses. (Indeed, note the close analogy between Eq. S37 and Eq. S20.) As a technical complication, Eq. S37 defines the observability Gramian with (the square root of) $\mathbf{C}^{-1}$ as the so-called 'readout matrix', whereas for consistency across our analyses (see below), we defined the most amplifying modes as the top eigenvectors of the observability Gramian with simply an identity readout matrix. We confirmed numerically that for the stochastic networks shown in (Fig. 4, Extended Data Fig. 5c,d, and Extended Data Fig. 8a,b,d,e), the top eigenvectors of the observability Gramians defined with these two different readout matrices were highly correlated (average 0.95 for the top $k = 5$ eigenvectors used in our analyses). For all other analyses of most amplifying modes (Fig. 3, Fig. 5f, Fig. 6e, Extended Data Fig. 3e Extended Data Fig. 7, and Extended Data Fig. 10f–h) we used deterministic networks (constructed de-novo, or fitted to stochastic nonlinear networks, or to experimental recordings; Methods 1.4.1 and 1.4.3) and so the choice of an identity readout matrix was justified by a lack of a meaningful definition of $\mathbf{C}$.

### S2.8 Optimal information loading: minimising energy with fixed persistent overlap

In our analyses of optimal information loading so far (Eqs. S25, S29 and S38), we have been considering a constraint on the norm of initial conditions in order to make optimization meaningful (as without some constraint, scaling the initial condition will always scale our chosen measures of performance due to the linearity of the dynamics). We now consider a biologically more relevant constraint that is better aligned with the optimization objectives used for most of our numerically optimized neural networks (Fig. 6 and Extended Data Figs. 3 and 11–13) and also standard in previous studies optimising recurrent neural networks[12–15]: the energy of the system as defined in Eq. S14. Using the duality of objective functions and constraints in constrained optimization, we seek the initial condition that minimizes energy, $\varepsilon_t$ while maintaining a fixed amount of overlap with the persistent mode, $p_t$ (Eq. S22), at the end of a time period of length $t$. This means we want to solve the following optimization problem:

$$\mathbf{x}^*(0) = \underset{\mathbf{x}(0)}{\operatorname{argmin}}\, \varepsilon_t, \text{ s.t. } p_t = 1 \tag{S39}$$

the solution to which is

$$\mathbf{x}^*(0) \propto \mathbf{V}\, \boldsymbol{\Omega}^{-1}(t)\, \boldsymbol{\rho}(t) \tag{S40}$$

$$\text{where } \rho_i(t) = \Xi_{i1}\, e^{\lambda_i t}, \text{ and } \boldsymbol{\Omega}(t) \text{ is defined in Eq. S18} \tag{S41}$$

In the limit when the delay period is very long, $t \to \infty$, we obtain

$$\mathbf{x}^*(0) \propto \mathbf{V}_{\mathrm{Q}}\, \boldsymbol{\Lambda}_{\mathrm{Q}}^{-1}\, \bar{\boldsymbol{\rho}} \tag{S42}$$

$$\text{where } \bar{\boldsymbol{\rho}} = \mathbf{V}_{\mathrm{Q}}^\top\, \mathbf{U}^\top\, \boldsymbol{\rho}_\infty \overset{\approx}{\propto} (1, 0, \dots 0)^\top \tag{S43}$$

[1] In other words, although Ref. 5 found fundamentally feedforward dynamics to be optimal for information maintenance, this was only true in general as long as a time-dependent decoder was allowed. The argument above instead suggests a normative justification for attractor, or integrator, dynamics that can render even a time-independent decoder near-optimal.

where $\mathbf{V}_Q$ and $\mathbf{\Lambda}_Q$ are respectively the matrix of eigenvectors (as columns) and diagonal matrix of associated eigenvalues of $\mathbf{Q}$, both ordered in decreasing order of the eigenvalues (which are all real, as $\mathbf{Q}$ is positive definite, Sections S2.3 and S2.6), and the approximate proportionality in Eq. S43 is due to $\boldsymbol{\rho}_\infty = \lim_{t\to\infty} \boldsymbol{\rho}(t) \propto (1, 0, \ldots 0)^\top$ (in analogy with Eq. S9), $\mathbf{u}_{(1)}$ being identical to the top eigenvector of $\mathbf{Q}$ (Section S2.6), and the eigenvectors of $\mathbf{Q}$ being orthonormal, because it is symmetric positive definite (Eq. S20). Were this proportionality exact, Eq. S42 could be simply solved as

$$\mathbf{x}^*(0) \overset{\approx}{\propto} \mathbf{u}_{(1)} \tag{S44}$$

In other words, the optimal information loading direction of an integrator network that achieves a fixed level of (asymptotic) persistent overlap while minimising the energy (or, conversely, that maximises the asymptotic persistent overlap given a fixed energy budget) is again approximately the top left eigenvector of the dynamics, i.e. (Section S2.6) the most amplifying mode.

In practice, Eq. S43 only holds approximately, with the ratio of the subsequent elements of $\bar{\boldsymbol{\rho}}$ to the first element being very small, but not quite zero. However, the final weighting of the eigenvectors of $\mathbf{Q}$ in expressing $\mathbf{x}^*(0)$ depends on the product of $\bar{\boldsymbol{\rho}}$ with $\mathbf{\Lambda}_Q^{-1}$ (Eq. S42), and the (diagonal) elements of $\mathbf{\Lambda}_Q^{-1}$ (which are the inverses of the diagonal elements of $\mathbf{\Lambda}_Q$) are ordered exactly oppositely to the elements of $\bar{\boldsymbol{\rho}}$. This partially cancels the preferential weighting of the top eigenvector of $\mathbf{Q}$ by $\bar{\boldsymbol{\rho}}$, and also makes Eq. S44 approximate. Nevertheless, in numerical simulations we found that Eq. S44 held to good precision in random linear integrator networks (such as those in Fig. 3d,e, Fig. 4 and Extended Data Figs. 4 and 5). In Extended Data Fig. 4a,b, we show the relationship between the energy produced by random initial conditions, scaled so that they all result in the same asymptotic persistent overlap, and their overlap with the most persistent versus most amplifying mode. We find that, in general, the overlap with the most amplifying mode (but not with the most persistent mode) is strongly and inversely related to the energy.

While our analytical derivations above concerned the asymptotic limit of an infinitely long delay period, Extended Data Fig. 4c–d show numerical simulations with systematically varied finite delay lengths. In particular, Extended Data Fig. 4c shows that the initial conditions that are optimal for different finite delays (Eq. S40), again in the sense of achieving a fixed persistent overlap by the end of the delay with minimal energy measured over the delay (Eq. S39), show an intuitive dependence on the length of the delay (Extended Data Fig. 4c, bottom). For very short delays, it is best to use inputs that are already aligned with the persistent mode, as there is no time for the dynamics of the system to contribute and gradually align the state with the persistent mode. However, for longer delays, the dynamics can have an increasing contribution, thus increasingly preferring the most amplifying mode over the persistent mode, in qualitative agreement with our analytical results. (We find that even for long delays, the overlap of optimal inputs is not all and none with the most amplifying and persistent modes, respectively, in line with our analysis of the approximate nature of Eq. S44.) In line with these results, the persistent mode achieves optimal to near-optimal performance (i.e. minimal energy) at short delays, but the most amplifying mode performs better (uses less energy) at longer delays (Extended Data Fig. 4d, bottom). (For symmetric networks, Extended Data Fig. 4c–d, top, which we only show for the completeness of analogy with Fig. 3d–e, no such trade-off is observed for the trivial reason that the most persistent and amplifying modes are identical.)

### S3 Analysis of canonical nonlinear systems with two stable fixed points

For linear systems that can exhibit persistent activity, we showed that inputs that align with the most amplifying mode generate the most noise-robust (and thus most easily decodable if using multiple inputs) persistent activity (Section S2, see also Figs. 3 and 4 and Extended Data Fig. 5). To develop insights as to how this result might generalise to nonlinear attractor systems, here we study two variants (either symmetric, Extended Data Fig. 6, top, or non-symmetric, Extended Data Fig. 6, bottom) of a canonical minimal nonlinear attractor system, based on the 'unforced Duffing oscillator' [16] (Eqs. 28 and 29; see Extended Data Fig. 6a for flow fields). Specifically, both variants exhibit 3 fixed points: a saddle point at the origin and 2 asymptotically stable fixed points (i.e. attractors) at $(\pm 1, 0)$ (Extended Data Fig. 6a, black crosses). Thus, the (one dimensional) 'coding subspace' of the system along which the two attractors can be best distinguished is the line connecting the two attractors, $\mathbf{c} = (1, 0)^\top$ (Extended Data Fig. 6a, $x_1$ axes). These two attractors divide the state space of the system into two halves, their respective basins of attraction, which are separated by a manifold, the so-called 'separatrix' (which in this case is one-dimensional; Extended Data Fig. 6a, dashed black lines). This means that the attractor to which the system ultimately converges from a particular initial state simply depends on which side of the separatrix that initial state lies.

#### S3.1 A local linearization approach based on linearizing around the ground state instead of an attractor

Standard linearization-based approaches to understand the behaviour of nonlinear attractor systems linearize the system's nonlinear dynamics around an *attractor* [9,17]. These approaches are valuable for understanding the *asymptotic* behaviour of the dynamics, once the state of the system is in the sufficiently close vicinity of the attractor, e.g. to understand its stability[9,16,17]. However, to study information loading, we need to understand the *initial* behaviour of the system. Thus, our approach is based on a linearization around the fixed point at the *origin* (see also Methods 1.4.2), which serves as the 'ground state'—the state which the system is thought to occupy before the 'stimulus' arrives, and thus with respect to which the magnitude of initial conditions is constrained. Indeed, the origin is an ideal ground state in this case (Extended Data Fig. 6a), not only because it is equidistant from the two attractors, but also because it sits on the separatrix. Thus, as long as the system is in the origin, or—the origin being a saddle point—even if it is perturbed away from the origin along the separatrix, it does not 'choose' between the two attractors.

### S3.2 Most persistent and amplifying directions for nonlinear dynamics

Linearizing Eqs. 28 and 29 around the origin yields the Jacobian matrix $\mathbf{J}_s = \begin{pmatrix} 1 & 0 \\ 0 & -1 \end{pmatrix}$ for the symmetric system, and $\mathbf{J}_n = \begin{pmatrix} 1 & 3 \\ 0 & -1 \end{pmatrix}$ for the non-symmetric system. Performing the same analyses on these linearized dynamics as those that we established for genuinely linear dynamics (Fig. 3; see also Methods 1.7.1), reveals that the (locally) 'most persistent' mode of the dynamics, $\mathbf{v}$ (i.e. the direction associated with the top (i.e. largest real) eigenvalue of the Jacobian), in this case represents an unstable direction of the system around the origin (Extended Data Fig. 6a, thin green lines). Although at first it may seem counterintuitive that an unstable direction is called 'persistent', this is actually consistent with our naming convention for linear dynamics not only based on its mathematical definition (see above), but also because—just like for the linear networks we studied—the persistent direction defined in this way corresponds to the coding direction of the system, i.e. $\mathbf{v} = \mathbf{c}$ (in both variants). In addition, these analyses also identify the 'most amplifying' mode, $\mathbf{a}$, (Extended Data Fig. 6a, thin red lines). Intuitively, the (locally) most amplifying mode is orthogonal to the separatrix in both systems (note different scales for the $x_1$ and $x_2$ axes in Extended Data Fig. 6a, causing an apparent lack of orthogonality in the bottom panel), i.e. it is the direction that allows the system to start moving away from the separatrix, and thus choose between the two attractors, the fastest. For the symmetric system, this direction is actually identical to the most persistent direction, $\mathbf{a}_s = \mathbf{v}$. However, for the non-symmetric system, the most amplifying mode is at an (approximately 57°) angle to the most persistent mode, $\mathbf{a}_n \approx (0.55, 0.83)^\top$ (unit normalized).

### S3.3 Locally most amplifying modes predict optimal inputs, analogous to linear networks

Numerical simulations of the original nonlinear dynamics confirmed that the particular information loading strategies we identified and compared for genuinely linear dynamics (Fig. 3a–c) lead to qualitatively similar behaviours in these nonlinear systems. Specifically, in analogy with our analysis of linear dynamics (Fig. 3b), given a fixed magnitude with respect to the origin (Extended Data Fig. 6a, pale blue ellipses), we compared three different initial conditions that were aligned with either the locally most persistent (Extended Data Fig. 6a, pale green arrows and circles) or most amplifying direction at the origin (Extended Data Fig. 6a, pale red arrows and circles), or with a randomly chosen direction (Extended Data Fig. 6a, gray arrows and circles). We then simulated dynamics starting from each of these initial conditions (Extended Data Fig. 6a, dark green, red, and black arrows) and measured how the overlap of the system's state with the most persistent direction evolved over time (Extended Data Fig. 6b). As expected, there were some unavoidable differences from the behaviour of the linear system: with these nonlinear attractor dynamics, eventually all trajectories converge to one of the attractors, and thus the overlap with the most persistent mode ultimately reaches the same asymptote (at unity) in all cases. As a consequence, the differences in trajectories corresponding to different initial conditions, and their overlap with the most persistent mode, can appear smaller than in the case of linear dynamics (this is especially the case for the non-symmetric system; Extended Data Fig. 6a, bottom). Nevertheless, by measuring the average overlap achieved between $t = 100$ and 250 while we systematically varied the direction of the initial conditions in the full 360° range, we found a monotonic relationship between the initial overlap of the state of the system with the most amplifying mode and this measure of 'late' overlap of the system with the most persistent mode Extended Data Fig. 6c, solid lines). This was qualitatively similar to the behavior of the simple linear networks of Fig. 3, where we also found a monotonic (in that case, linear) relationship between initial amplifying and late persistent overlap (Extended Data Fig. 6c, dashed lines; with late overlap measured between $t = 0.8$ and 2 s). This meant that the most amplifying direction was indeed the optimal direction for 'information loading' in both the nonlinear and linear dynamical systems (Extended Data Fig. 6c, pale red circles and squares). In comparison, with both nonlinear and linear dynamics, random directions or the most persistent direction—when it was different from the most amplifying direction (i.e. in the non-symmetric system; Extended Data Fig. 6a–c, bottom)—achieved lower information loading performance by this measure (Extended Data Fig. 6c, gray and pale green circles and squares).

In summary, we have shown that—just like in the case of linear persistent dynamics—the locally most amplifying direction around the ground state of these nonlinear systems represents the optimal information loading strategy, that is better than 'pushing' the system directly in the direction of the desired attractor (most persistent mode), or in random directions. The intuition is that the locally most amplifying mode is the direction that allows the system to leave the vicinity of the separatrix and travel towards the desired attractor the fastest—this gain in speed quickly offsets the initial disadvantage in closeness to the attractor in comparison to inputs using the most persistent mode. While the size of the effects we obtained were small in some cases, these depended on our choice of parameters and were in part due to the simplicity and low-dimensionality of these systems. Thus, as the next step, we studied these effects in high-dimensional neural networks with nonlinear dynamics.

### S4 Analysis of the nonlinear attractor networks of Fig. 2

To further understand how our theoretical insights gained from analysing linear networks (Section S2, see also Fig. 3 and Extended Data Fig. 5) may generalize to nonlinear attractor networks, we performed the same local (around the origin) linearization of the nonlinear attractor networks of Fig. 2 that we introduced in the previous section (Section S3; see also Methods 1.4.2).

#### S4.1 The relationship between subspaces extracted from nonlinear dynamics and by local linearization

First, we linearized the dynamics (Eqs. 1 and 2) of both the symmetric and unconstrained networks of Fig. 2 (Methods 1.4.2) and identified the associated most persistent and amplifying modes around the origin based on this local linearization. We then measured the overlap of the (5-dimensional) subspaces spanned by these modes with the subspaces that we extracted from the activities of the original nonlinear networks without linearizing their dynamics (Methods 1.7.1), analysed in Fig. 2f and l. We found that the locally most *persistent* modes overlapped strongly with the persistent subspace of the nonlinear dynamics in which the attractors lie (Extended Data Fig. 7a, green, 'persistent subspace') and overlapped poorly with the persistent nullspace (Extended Data Fig. 7a, green, 'persistent nullspace'). Subspaces (dimensionally matched) spanned by *random* modes showed the opposite pattern: they had small overlaps with the persistent subspace and large overlaps with the persistent nullspace, simply due to the higher dimensionality of the latter (Extended Data Fig. 7a, black, 'persistent subspace' and 'persistent nullspace'). Critically, for both locally most persistent and random modes, these overlaps were essentially identical between symmetric and unconstrained networks (Extended Data Fig. 7a, compare top and bottom panels). In contrast, the overlap of the locally most *amplifying* modes with the persistent subspace and nullspace of the nonlinear dynamics clearly differentiated between symmetric and unconstrained networks. Specifically, in symmetric networks, the most amplifying modes overlapped strongly with the persistent subspace and poorly with the persistent nullspace—as they were identical to the most persistent modes in this case (Extended Data Fig. 7a, top, red, 'persistent subspace' and 'persistent nullspace'). In unconstrained networks, however, they showed a similarly strong overlap with the persistent nullspace as with the persistent subspace (Extended Data Fig. 7a, bottom, red, 'persistent subspace' and 'persistent nullspace').

#### S4.2 Locally most amplifying modes predict optimal inputs, analogous to linear networks

The results above suggested that our original findings, showing a double dissociation in the efficiency of persistent subspace and nullspace inputs (initial conditions) between symmetric and unconstrained networks (Fig. 2f and l, red and green), are likely explained by how much these inputs align with the locally most amplifying inputs. Indeed, we found that the subspace of numerically optimized inputs (studied in Fig. 2b–e, h–k, and f and l, black) had a substantially larger overlap with the locally most amplifying mode in both classes of networks than with random subspaces, or—in unconstrained networks, in which persistent and amplifying modes are distinguishable—than with the locally most persistent modes (Extended Data Fig. 7a, 'optimal', compare red to black and green). This pattern of overlaps of optimal inputs with the locally most amplifying, persistent and random subspaces was closely analogous to those that we found for the optimized initial conditions in the purely linear networks of Extended Data Fig. 5c,d,g,h (Extended Data Fig. 7a, 'optimal (lin. model)'). We also found a similar pattern of results when performing local linearizations in optimized ring attractor networks (Extended Data Fig. 3e). In sum, these results generalise the results of Section S3 to high dimensional nonlinear neural networks with multiple attractors. They indicate that the locally most amplifying and persistent directions that can be extracted from nonlinear attractor networks around the ground state play functionally similar roles in information loading to the corresponding modes of linear networks of which we have a firm analytical understanding.

#### S4.3 Overlap with most persistent mode in linear networks predicts decoding accuracy in nonlinear networks

In deterministic linear networks, to allow analytical insights, we used the 'overlap with persistent modes' as a measure of performance (Fig. 3c–e, Extended Data Fig. 5a,b,e,f). In contrast, the measure of performance in stochastic nonlinear networks that we regarded as ultimately relevant, and directly applicable to experimental data, was based on the accuracy of a linear decoder (Fig. 2f,l, Fig. 5b,c, Fig. 6c,d). To study the relationship between these two measures, we simulated both the original stochastic nonlinear networks of Fig. 2 and the deterministic linear dynamics obtained from their local linearization (introduced above), and measured the time evolution of each of these metrics in the corresponding networks when their dynamics were started from initial conditions that were optimized for the decoding accuracy of the nonlinear dynamics (Methods 1.3.1 and 1.7.2) while constrained to be within the locally most persistent, most amplifying, or a random subspace, as determined by the local linearization, or without any subspace constraints (i.e. as originally done in Fig. 2b–e, h–k, and f and l, black). For a fair comparison, each of the subspaces used for constraining initial conditions were 5-dimensional.

In general, the overlap with the most persistent modes in the locally linearized networks showed somewhat richer dynamics that in the *de novo* linear networks, in which it remained exactly constant or at most decayed exponentially (Fig. 3c,e). Specifically, the overlap measure we used here showed a brief initial decrease for symmetric networks (Extended Data Fig. 7b, top) and mildly non-monotonic time courses for unconstrained networks (Extended Data Fig. 7b, bottom). These were due to measuring overlap here with a 5-dimensional most persistent subspace (appropriate for networks distinguishing between 6 cue conditions), including some directions that were associated with eigenvalues that could have slightly negative real parts and non-zero complex parts (besides the one that was defined to have a zero eigenvalue, see Methods 1.4.2). In contrast, in Fig. 3, we measured overlap with the single most persistent direction of the dynamics (distinguishing between only two cue conditions) that was constructed to be exactly persistent, i.e. with a zero eigenvalue.

Despite these differences, we found that the overlap measure for the locally linearized networks showed qualitatively similar patterns as in the *de novo* large linear networks of Fig. 3e. For symmetric networks (Extended Data Fig. 7b, top), the overlap was largely constant in time for each initial condition, with most persistent and amplifying inputs achieving identical performances Extended Data Fig. 7b, top, red and green), substantially above that of random inputs (Extended Data Fig. 7b, top, black), and on par—and by this measure, even somewhat better—than that of optimal inputs (Extended Data Fig. 7b, top, blue). For unconstrained networks (Extended Data Fig. 7b, bottom), the overlap for most persistent inputs again remained roughly constant in time (Extended Data Fig. 7b, bottom, green). For most amplifying inputs, overlap started from a lower level initially, but it eventually overtook the level achieved by most persistent inputs (Extended Data Fig. 7b, bottom, red). Random inputs performed worst (Extended Data Fig. 7b, bottom, black), and optimal inputs showed a similar trajectory to that of most amplifying inputs (initially low overlap which then increased to above that achieved by most persistent inputs), though at a somewhat lower performance by this measure (Extended Data Fig. 7b, bottom, blue vs. red).

Not only did these overlap measures of linearized networks retain some important qualitative characteristics of the overlap measures used in *de novo* linear networks, they also showed a clear relationship with our ultimate measure of interest: decoding accuracy in the corresponding stochastic nonlinear networks (using a 'noise-matched' scenario—see Fig. 4a—in which all networks had the same level of noise in their dynamics; Methods 1.4.1). In symmetric networks, most persistent and amplifying inputs achieved maximal performance (as did the optimal inputs), with random inputs performing substantially more poorly (Extended Data Fig. 7c, top). In unconstrained networks, most amplifying (and optimal) inputs outperformed most persistent inputs, with random inputs performing worst again (Extended Data Fig. 7c, bottom). Thus, the overall rank ordering of different inputs for information loading, as measured by eventual decoding accuracy, is well predicted by their rank ordering on the overlap measure.

### S4.4 Fitting linear dynamics reveals information loading strategies in nonlinear networks

We have shown so far that the same principles (and measures) determining the efficiency of different information loading strategies that we analytically identified in linear networks also apply to nonlinear networks. However, our approach for this was based on a local linearization of the original nonlinear dynamics, which required knowledge of the true equations governing the dynamics of these networks. This is obviously not available for experimental data. Thus, in order to be able to apply our theoretically derived measures of optimal information loading without having access to the true dynamics of the system, we analyzed experimental data by fitting neural responses by linear neural networks (Fig. 5d–f; see also Methods 1.4.3). As a consequence, we wanted to validate that this fitting-based approach provides meaningful results, and has the capacity to reveal information loading strategies in nonlinear networks even when

1. we do not have access to the true dynamics but only to samples of activities generated by those dynamics; and
2. we also cannot assume that the true dynamics are linear.

For this, we repeated the same analyses with our simulated nonlinear networks while using 'ground truth' information loading strategies, i.e. inputs confined to the most persistent, or most amplifying, or random, or optimal subspaces as defined above (Section S4.3).

As in the previous section (Section S4.3), we used a 'noise-matched' scenario—see Fig. 4a—in which all networks had the same level of noise in their dynamics; Methods 1.4.1). A more realistic comparison might require 'performance matched' simulations, as in Fig. 3c, in which different networks are allowed to have different amounts of noise in their dynamics such that it is instead the asymptotic level of (decoding) performance that is matched between them. However, we found that our main metric of subspace overlap was robust to changes in noise levels, so we expect that all our results hold to high precision in the performance matched regime as well.

For symmetric networks, we found that linear dynamical systems fitted to responses that resulted from using optimal inputs yielded similar dynamics to those following either most persistent or most amplifying inputs (Extended Data Fig. 7d; top, compare 'persistent', 'amplifying', and 'optimal'). Initial overlap was well above chance with both the most amplifying and persistent modes (though higher for the former), such that the overlap with most amplifying modes remained constant, while the overlap with most persistent modes increased over time. In contrast, the initial overlap resulting from random inputs with either persistent or amplifying modes was close to chance (with overlap with amplifying once again remaining constant, while overlap with persistent increasing over time). Note that all of these were unlike the patterns we saw in experimental data (Fig. 5f).

For unconstrained networks, most amplifying and optimal inputs again resulted in similar dynamics (Extended Data Fig. 7d; bottom, compare 'amplifying' and 'optimal'). Initial overlap was high with most amplifying modes but low (at chance) with persistent modes before the dynamics ultimately overlapped strongly with the persistent modes and weakly with the amplifying modes. This was in contrast to dynamics following persistent or random inputs, in which there was no clear difference in initial overlaps with most amplifying vs. most persistent modes, neither was there a clear decrease in overlap with the most amplifying modes over time (Extended Data Fig. 7d, compare 'persistent' and 'random'). Note that experimental data was only consistent with the former but not the latter pattern of results (Fig. 5f).

### S5 Fisher information in task-optimized ring attractor networks

Here, we note that the cosine term in the cost function (Eq. 4) for the ring attractor networks that we optimize in Methods 1.3.4, quantifying the precision of the decoded angle, is closely related to a cost measuring the population Fisher information about angle. Intuitively, both cost functions quantify the ability of the network to perform fine discrimination of stimulus angles. To see this more formally, recall that the Fisher information in this case is

$$\mathcal{I}_{\theta^{(c)}} = -\int \mathcal{P}\left(\mathbf{r}|\theta^{(c)}\right) \left.\frac{\partial^2}{\partial\theta^2}\right|_{\theta\,=\,\theta^{(c)}} \ln \mathcal{P}\left(\mathbf{r}|\theta\right)\, \mathrm{d}\mathbf{r} \tag{S45}$$

where, as in Methods 1.3.4, $\theta^{(c)}$ is the true stimulus angle on a trial (the target for the decoder), $\mathbf{r}$ is the population response, and $\mathcal{P}\left(\mathbf{r}|\theta\right)$ is the distribution of responses the network generates to some stimulus angle $\theta$. In the limit of a sufficiently large population, the maximum likelihood estimator, $\hat{\theta}_{\mathrm{ML}}$ achieves the same Fisher information as the full population vector, and is distributed as a (circular) Gaussian (von Mises) distribution centered on the true orientation, $\theta^{(c)}$, with some constant (circular) concentration, $\kappa$, so we can write

$$\mathcal{I}_{\theta^{(c)}} \simeq -\int \mathcal{P}\left(\hat{\theta}_{\mathrm{ML}}|\theta^{(c)}\right) \left.\frac{\partial^2}{\partial\theta^2}\right|_{\theta\,=\,\theta^{(c)}} \ln \mathcal{P}\left(\hat{\theta}_{\mathrm{ML}}|\theta\right)\, \mathrm{d}\hat{\theta}_{\mathrm{ML}} \tag{S46}$$

$$= \kappa \int \mathcal{P}\left(\hat{\theta}_{\mathrm{ML}}|\theta^{(c)}\right) \cos\left(\hat{\theta}_{\mathrm{ML}} - \theta^{(c)}\right) \mathrm{d}\hat{\theta}_{\mathrm{ML}} \tag{S47}$$

Assuming that the population vector-based decoder we use for training networks, $\hat{\theta}$ (Eq. 5 in Methods 1.3.4), is an efficient estimator that approximates the maximum likelihood decoder, $\hat{\theta} \simeq \hat{\theta}_{\mathrm{ML}}$[2], and substituting the integral over the distribution of the estimate with an empirical average over its stochastic realizations, we can further rewrite the Fisher information as

$$\mathcal{I}_{\theta^{(c)}} \simeq \kappa \left\langle \cos\left(\hat{\theta}_{\mathrm{ML}} - \theta^{(c)}\right) \right\rangle \tag{S48}$$

Eq. S48 is thus identical to the integrand of the first term in Eq. 4 up to a multiplicative constant (which can be incorporated into $\alpha^{(1)}_{\mathrm{nonlin}}$ in Eq. 4), an additive constant (the 1 inside the square bracket in Eq. 4), which does not matter for optimization, and a sign-flip, because we are maximizing Fisher information in Eq. S48 but minimizing the cost in Eq. 4. Therefore, minimizing (the first term in) Eq. 4 also (approximately) maximizes Eq. S45 (averaged over time points at which the network is decoded and over true stimulus angles), and vice versa.

[2] We also checked empirically that, in line with these assumptions, the empirical distribution of $\hat{\theta}$ was well approximated by a von Mises distribution centered on $\theta^{(c)}$ with a constant concentration across target angles.
